# Multiplexed functional analysis of TAP2 variants in regulating MHC-I cell surface abundance reveals overexpression of PLK1 downregulates antigen presentation

**DOI:** 10.1101/2024.09.30.615919

**Authors:** Gregory Nagy, Rebecca Presley, Hyeongseon Jeon, Lynn Moreira, Shruti Dhar, Olivia Tull, Burak Soysal, Anika S. Yadati, Kayleigh D. Cook, Laura E. Schnell, Arkobrato Gupta, Tapahsama Banerjee, Dongjun Chung, Philip N. Tsichlis, Jeffrey D. Parvin

## Abstract

The abundance of MHC-I on the cell surface depends on its association with antigenic peptides transported to the endoplasmic reticulum by the Transporter Associated with Antigen Processing (TAP), a peptide channel composed of TAP1 and TAP2. We functionally screened over 1400 TAP2 variants for effects on MHC-I cell surface abundance. Amino-acid substitutions of loss of function (LOF) variants clustered in the NTP-binding domain and along the TAP1-TAP2 binding interface, suggesting that these variants interfered with TAP conformational changes associated with peptide transport. Some LOF variants carried potential phosphomimetic substitutions; one such substitution at Ser251 was embedded in a sequence context consistent with the phosphorylation motif of PLK kinases. Inhibition of PLK kinases increased MHC-I surface expression, while overexpression of PLK1 in cells decreased MHC-I in wild type TAP2, but not in TAP2-S251A-expressing cells. Importantly, site-specific phosphorylation of TAP2 in human tumor samples correlated with alterations of gene expression in the cell cycle and antigen presentation pathways, consistent with the notion that phosphorylation downregulates antigenic peptide transport in human tumors. These data strongly support the hypothesis that in cancer cells, TAP2 phosphorylation by PLK1, and perhaps other kinases, can downregulate the antigen presentation process.

## INTRODUCTION

T-cell mediated immunity of virus-infected cells or of tumor cells depends on the T cell receptor (TCR) on CD8+ T lymphocytes recognizing antigenic peptides bound by the Major Histocompatibility Complex Type I (MHC-I). MHC-I consists of one of the HLA-A, HLA-B, or HLA-C proteins in complex with β2-microglobulin. The antigenic peptides are derived from proteolysis of cytoplasmic proteins, which gives rise to peptides 8 to 13 residues in length. The peptides must be transported from the cytoplasm to the lumen of the endoplasmic reticulum (ER) whereupon the Peptide Loading Complex (PLC) mediates their stable binding to the different isoforms of the HLA-A/B/C protein in MHC-I. Once a stable peptide-MHC-I complex is formed, glucosidase II cleaves a glucose moiety in an N-linked glycan on MHC-I and the complex is released for relocalization to the cell surface where it can be recognized by the TCR [1–3].

The Transporter Associated with Antigen Processing (TAP), a central component of the antigen presentation machinery, is composed of two polypeptides, TAP1 (*ABCB2*) and TAP2 (*ABCB3*). Pathogenic germline mutations of either TAP1 or TAP2 result in a severe immunodeficiency, called Bare Lymphocyte Syndrome [1,4]. In the absence of TAP1 or TAP2 function, peptides are not transported from the cytoplasm to the ER lumen and the MHC-I remains unbound and intracellularly retained in the ER.

The process of antigen presentation has several steps, beginning with the proteasome-mediated degradation of cellular proteins, which generates peptides that are transported into the endoplasmic reticulum by TAP. TAP controls the first step in peptide loading to MHC-I, a process that depends on the peptide loading complex (PLC), which also includes Tapasin, TAPBPR, Calreticulin, and ERp57 [1]. The peptide/TAP complex produced by the binding of immunogenic peptides to TAP, binds ATP and undergoes a major conformational change, which alters its orientation from inward facing toward the cytoplasm, to outward ER lumen facing, releasing the peptide into the ER lumen. TAP1 and TAP2 each have similar architectures with a globular NTP-binding domain next to the transmembrane domain. Binding to ATP causes the two NTP-binding domains to interact, closing the gap between the transmembrane domains on the cytoplasmic side, and opening the channel on the side of the ER lumen [5] (Figure 1). The PLC loads peptides onto MHC-I, and stable MHC-I-peptide complexes are released by the action of glucosidase II. The released MHC-I/peptide complexes relocalize to the cell surface where they can be recognized by the TCR of CD8+ T cells (Figure 1) [1].

**Figure 1.**
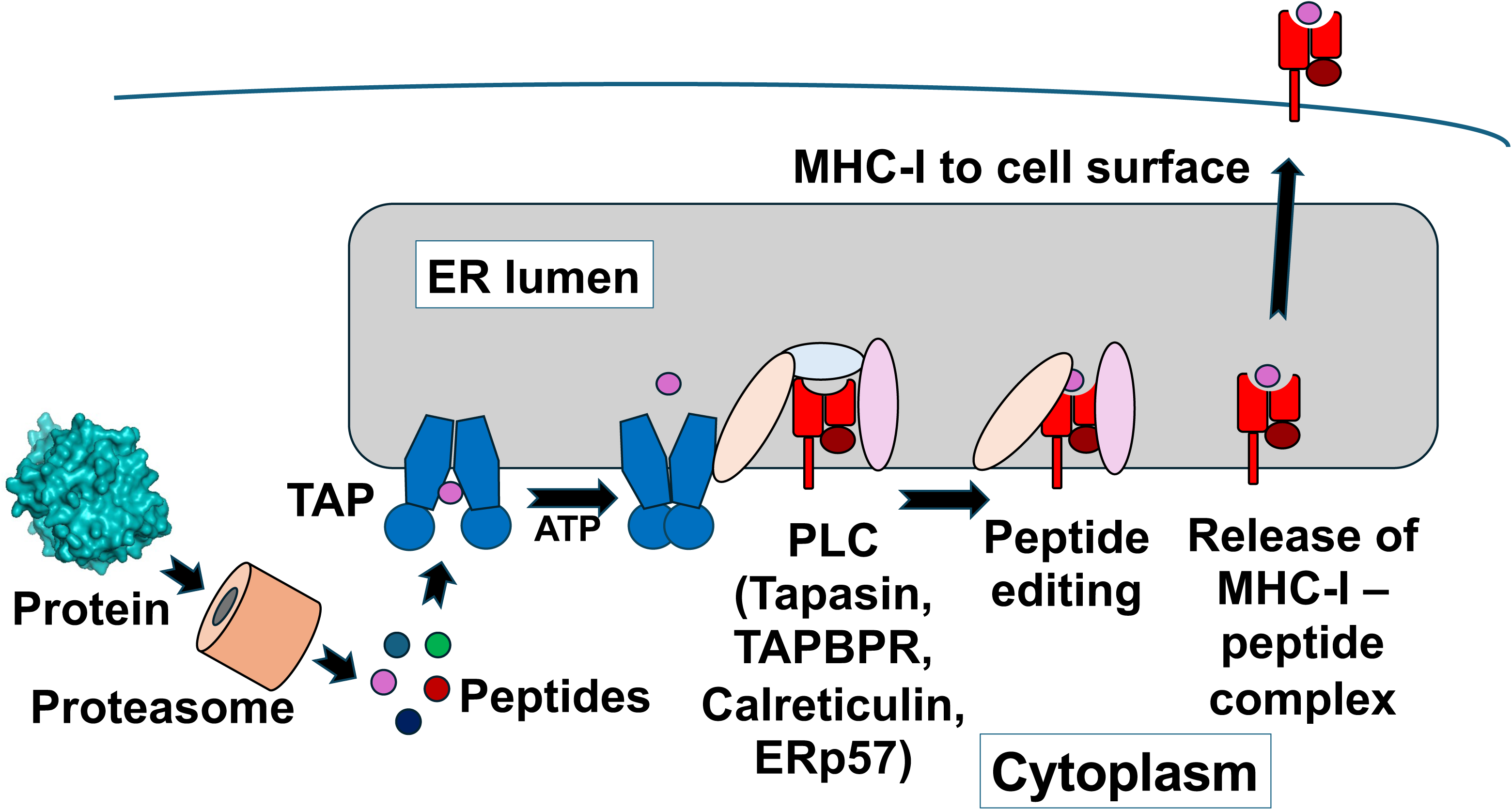
Role of TAP in antigen presentation. A schematic is shown for the pathway by which proteins expressed in a cell are subjected to antigen presentation. Cellular proteins are degraded by the proteasome into peptides of length 8-13 residues, and peptides are actively transported from the cytoplasm to the lumen of the endoplasmic reticulum (ER) by TAP. Peptides in the ER lumen bind to various isoforms of HLA A/B/C in complex with β2-microglobin in concert with other Peptide Loading Complex components that modify the peptide and ensure a stable peptide-MHC-I interaction. Stable MHC-I-peptide complexes are released from the ER lumen and relocate to the cell surface.

Several observations suggest that the TAP peptide transporter is a critical component of the antigen presentation machinery. First, in the absence of functional TAP, there are no peptides available to bind MHC-I in the ER lumen, and MHC-I remains in the ER. Consistent with this observation, germline loss of function (LOF) mutants of TAP1 or TAP2 give rise to a severe immuno-deficiency, the Bare Lymphocyte Syndrome [1,3]. The central regulatory role of TAP in the antigen presentation process is highlighted by the mechanisms employed by herpes viruses to evade host detection of the virus-infected cells. Specifically, all viruses of the herpes family express a protein that binds TAP and blocks its peptide transporting function, thereby inhibiting MHC-I expression on the cell surface and antigen presentation [3,6–8]. Importantly, the activity of TAP in antigen presentation appears to be regulated by phosphorylation [9].

In this study, we used a multiplexed assay to score the effects of missense substitutions of TAP2 on the function of the TAP peptide channel. To this end, we generated a library of variants of TAP2, and we established an assay to score their impact on the localization of MHC-I on the cell surface. Although in this screen we examined the effects of substitution in many TAP2 amino acids predicted to be functionally significant, we paid special attention to potential phosphorylation sites, both because prior evidence had suggested that TAP activity may be regulated by phosphorylation [9], and because the phosphorylation of such sites could be therapeutically targeted. Our results, based on the screen of 1404 TAP2 variants confirmed the functional importance of ATP and peptide binding residues, and suggested that posttranslational modifications, including phosphorylation, functionally regulate TAP2. Follow up validation assays and bioinformatic analyses of publicly available cancer data, confirmed the hypothesis that antigen presentation in human cancer can be regulated by TAP2 phosphorylation. For several of the putative TAP2 phosphorylation sites indicated by the multiplexed analysis, we also identified the kinases that are likely to phosphorylate them. One of these sites (Ser251) was identified as a likely target of PLK1. The predicted role of PLK1 was confirmed by experiments showing that PLK1 overexpression downregulates the abundance of MHC-I on the cell surface in cells engineered to express the wild type TAP2, but not in cells expressing the TAP2-S251A-mutant protein. Importantly, pharmacological inhibition of this kinase increased MHC-I expression on the surface of a cancer cell line with constitutively low MHC-I cell surface expression.

## RESULTS

### Multiplexed assay to measure the effects of TAP2 variants on the regulation of MHC-I abundance on the cell surface

To explore the mechanisms by which TAP2 regulates antigen presentation, we constructed a multiplex library of TAP2 variants, and we screened it to identify variants with altered function. To minimize phenotypic variabilities due to differences in the level of expression between variants, we established a cell culture system which enforced the library plasmid to integrate only once in the same single genomic site in all the cells. This was done by integrating a single copy of a retroviral vector containing the attP recombination site downstream of the RSV-LTR promoter [10] (Figure S1) into the genome of the MHC-I-positive U2OS cells (Figure 2A). We then knocked out the endogenous *TAP2* gene using CRISPR (Figure 2A). To select for TAP2 knockout cells, CRISPR treated and control cells were stained with FITC-conjugated anti-MHC-I (HLA A/B/C) antibody and analyzed by flow cytometry. Cells with reduced cell surface staining were flow sorted as knock-out cells (U2OS-TAP2-KO), and these cells had no detectable TAP2 protein on immunoblot analysis (Figure 2B). TAP2 knock-out cells were viable, morphologically unchanged, and they continued to grow similarly to the unmodified cells. The next step in the development of the multiplex assay was to rescue TAP2 function by integrating a single copy of the TAP2 construct in the attP site of the U2OS-TAP2-KO cells. To this end, we transfected the cells with the TAP2-WT construct (Figure S1). The TAP2 sequence in this construct carried silent nucleotide substitutions that rendered it resistant to targeting by the sgRNAs. The transfected TAP2 construct was promoterless and contained an attB site upstream of an open reading frame encoding hygromycin resistance, a P2A translation re-start signal, and the sgRNA-resistant TAP2-WT sequence with a polyadenylation signal (Figure S1). AttB/AttP recombination by the Bxb1 enzyme (“Int” in Figure S1) placed the Hygro-P2A-TAP2 sequences under the control of the single copy doxycycline-regulated RSV LTR. In the presence of doxycycline (dox), wild type TAP2 was re-expressed (Figure 2B). Analysis of parental, KO, and WT rescue cells for MHC-I cell surface abundance by flow cytometry showed that whereas MHC-I cell surface expression was greatly reduced in the KO cells, its expression was restored by the re-expressed wild type TAP2 (Figure 2B).

**Figure 2.**
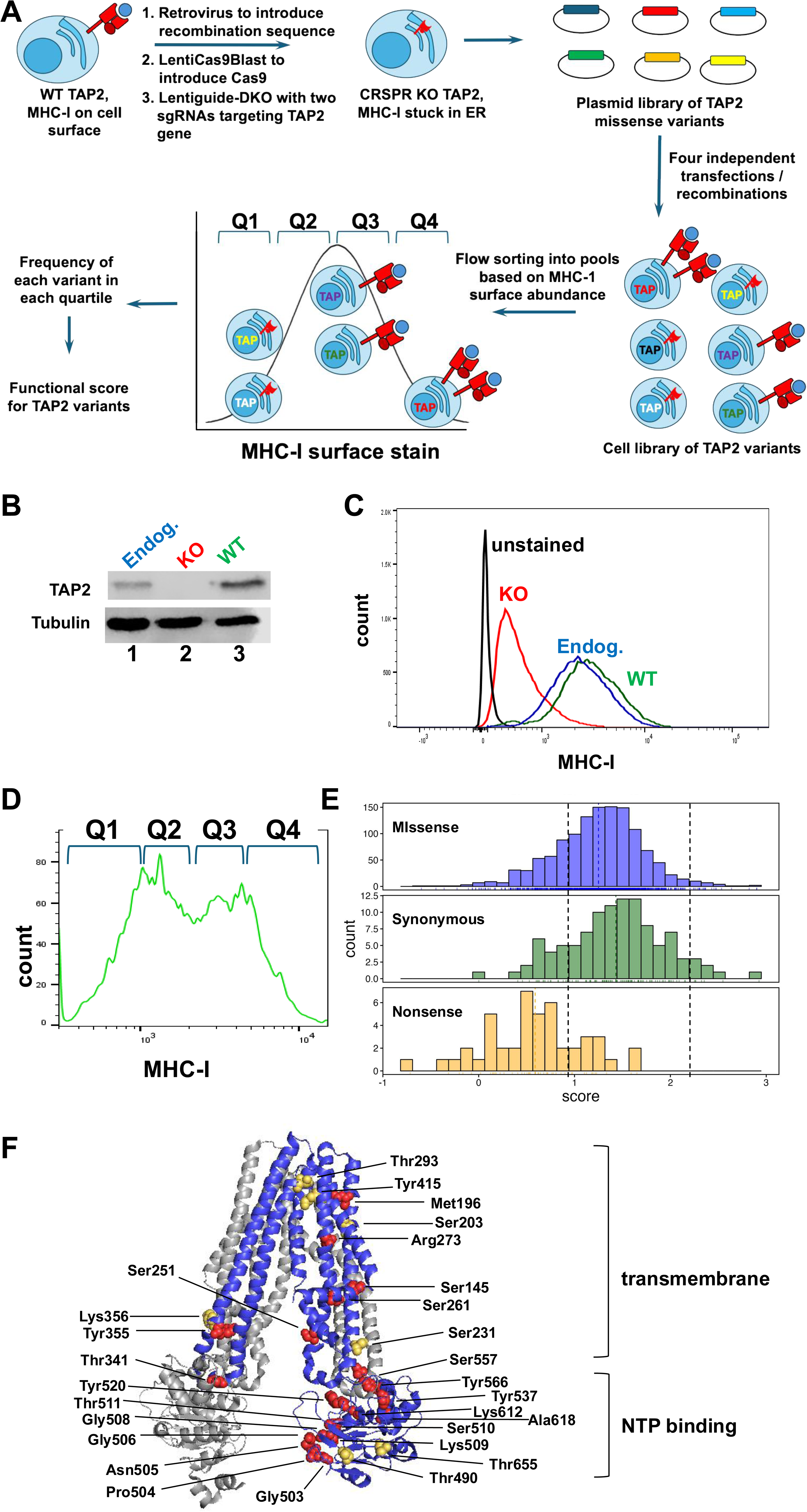
The multiplexed assay for TAP2 function. **A.** A schematic of the multiplexed assay is shown. U2OS cells express on the cell surface MHC-I in association with antigenic peptide. In cells with TAP2 knocked out, the MHC-I is retained intracellularly in the ER and with low detection on the cell surface. The TAP2-KO cells are rescued with a multiplexed library of TAP2 genes/proteins, and each rescued TAP2 contains a single missense substitution at a targeted codon/amino acid residue. For loss of function (LOF) variants of TAP2, the MHC-I is retained intracellularly. For TAP2 variants with normal function, restoration of localization of MHC-I on the cell surface will be at the same abundance as observed with wild-type TAP2. Some of the TAP2 variants will be hyperfunctional and have a higher abundance of MHC-I on the cell surface than observed with cells expressing wild-type TAP2. On staining cells for HLA-A/B/C and flow cytometry, the population of cells is separated according to the level of MHC-I on the cell surface. Cells are collected in quartiles based on the intensity of MHC-I stain, and the frequency of detection of each TAP2 variant with the high, medium, or low quartiles of cells is converted to a functional score. **B.** Immunoblot stained for TAP2 in the parental U2OS cell line (Endog.), in the TAP2 knock out (KO), and the rescue with wild-type TAP2 (WT). Tubulin is shown as a loading control. **C.** A histogram is shown for a flow cytometry experiment measuring the intensity of HLA-A/B/C stain (MHC-I, x-axis) and the count of cells at each staining intensity (y-axis). The results are for parental U2OS cells (Endog.), TAP2-KO cells (KO), and TAP2-KO cells rescued with wild-type (WT) TAP2. Unstained U2OS cells are shown. Results from three experiments were combined in this histogram. **D.** U2OS TAP2-KO cells were rescued by recombination of a library of TAP2 missense variants into a single genomic site; each cell expresses a different TAP2 variant from the multiplex library. The abundance of MHC-I on the cell surface was measured by flow cytometry and shown as a histogram (*bottom*) with MHC-I abundance (x-axis) versus cell count (y-axis). **E.** Histogram of functional scores and results for TAP2 missense variants (*top*), synonymous variants (*middle*) and nonsense variants (*bottom*). A dashed line indicated a functional score of 0.92, which is based on the mean functional score in the population of synonymous variants minus one standard deviation. A second dashed line at a functional score of 2.2 indicated the threshold for classification as hyperfunctional variants. **F.** The 3D structure of TAP1 and TAP2 is shown (PDB 5U1D). TAP1 is shown as gray, and TAP2 is shown as blue. Residues at which four or more variants were LOF are shown in red, and residues at which four or more variants were either LOF or hyperfunctional are shown in yellow. Drawn with PyMOL.

To identify TAP2 variants with altered function, we generated and screened a barcoded library of TAP2 variants, which was constructed by targeting the codons encoding 96 of the 686 amino acid residues of the TAP2 protein (Table S1). The library was cloned in the promoterless plasmid described above (Figure S1A). Library clones transfected in the genetically modified U2OS cells described above, would not be expressed unless integrated by AttB/AttP recombination downstream of the Dox-regulated RSV LTR. Since all expressed variants would be integrated at the same site in each cell, their level of mRNA expression would be expected to be similar in all transduced cells. To select the residues to target for the multiplexed library, we included a residue known to form a salt bridge to Tapasin (Asp16) and five residues involved in known peptide interactions (R368, R369, R380, M413, and E417) [11]. Other target residues included four phosphorylated residues and eleven ubiquitinated residues (PhosphositePlus website, https://www.phosphosite.org/), 12 encoding variants listed as variants of unknown significance (VUS) in ClinVar [12], two residues with variants listed in ClinVar as benign, and four substitutions correlated with human cancer [11]. We tested 56 conserved serine, threonine, and tyrosine residues; conservation was determined by comparison of the TAP2 proteins of 10 different species. The targeted codons also included those encoding 19 of the residues in the peptide binding domain (residues 301-389 and 414-443) [8] and all the residues in the Walker A NTP-binding motif (residues 503-511) [5,13].

To construct the library we employed an inverse PCR strategy [14], which was designed to replace each of the selected codons with up to 20 variants (19 missense changes and 1 nonsense mutation at 96 codons = 1920 possible variants). The plasmid library contained 1786 variants, or 93% of the potential variants (Figure S2). In addition, the library contained some variants arising from missense substitutions at codons we did not target. These variants were presumably the products of PCR errors, and some of them were sufficiently abundant to be scored in the functional assay. As will be discussed below, out of all the tested variants in the plasmid library, we obtained functional scores for 1404 TAP2 variants.

The TAP2 variant library was transfected into the U2OS TAP2-KO cells; four independent transfections/recombinations were separately analyzed. Following selection in hygromycin, cells were stained using FITC-conjugated anti-MHC-I antibody and sorted into four quartiles of staining intensity (diagrammed in Figure 2A, D). The genomic DNA from each quartile of sorted cells was isolated, and the barcodes were retrieved by PCR. Sequencing of the barcodes revealed the frequency of all variants in each quartile. Results from the four independent replicates were combined and averaged. A functional score was calculated based on frequency of a variant in each quartile relative to the total pool of variants (see Methods), and the functional scores of missense, synonymous, and nonsense variants were compared. In all, the functional impacts of 1252 missense variants, 106 synonymous variants, and 46 nonsense mutants were scored for MHC-I cell surface abundance (Tables S2-S4). Histograms showing the distribution of functional scores of TAP2 missense, synonymous, and nonsense variants are shown (Figure 2E). Synonymous variants had a mean functional score of 1.43, and nonsense variants had a mean functional score of 0.59. The functional scores from the missense variants (mean = 1.25) were shifted to the left relative to the synonymous variants. Based on the distributions of the functional scores of synonymous and nonsense variants, we classified variants with scores less than one standard deviation from the mean of the synonymous variants as loss of function (LOF; < 0.92). Of the 1252 missense variants for which a functional score was calculated, 273 were classified as LOF. Variants with a functional score > 2.2 (top 2% of missense variant scores) were classified as hyperfunctional.

Among the residues tested in the multiplexed functional assay, 12 had variants listed as VUS in the ClinVar database. Of these 12, ten were scored, with nine scoring in the functionally normal range, and one, S219F, scoring as LOF (Table S5). Two variants listed in ClinVar as benign, R313H and T665A, were functionally normal in the multiplexed assay, and one nonsense variant list in ClinVar as pathogenic, R273*, scored as LOF (Table S5).

### TAP2 residues with multiple LOF substitutions

We identified amino acid residues which were the target of multiple substitutions that gave rise to LOF missense variants. Twenty-one residues were the target of four or more such LOF substitutions (Figure 2F). These residues with multiple LOF variants include Asp16, which is known to form a salt bridge with Tapasin [11]. Of interest, four missense substitutions at Asp16 were loss of function, and in two variants classified as functionally normal, Asp was replaced by basic residues (D16R and D16H), suggesting that although the salt bridge contact of TAP2 with Tapasin might be functionally significant, it was not essential. Met196, Thr341, and Tyr355 are known to contribute to the binding of the peptide cargo [11], and multiple substitutions at these residues gave rise to LOF variants. Many residues in the Walker A motif / NTP binding domain [15] had four or more LOF substitutions, including Gly503, Pro504, Asn505, Gly506, Gly508, Lys509, Ser510, and Thr511. Lys612, a ubiquitin acceptor (PhosphositePlus) had five detected LOF variants. Ala618 and Arg273 were included in the list of 96 targeted codons since they had VUS listed in ClinVar. Interestingly, the A618S variant listed as VUS in ClinVar had a score consistent with normal function, but inclusion of this residue in the analysis revealed four LOF variants (A618T, A618D, A618V, and A618R). The R273Q variant listed in ClinVar as VUS had normal function, but ten other missense variants at this position were LOF.

At seven residues there were four missense variants that included both LOF and hyperfunctional variants. These were Leu111, Ser203, Thr293, Lys356, Tyr415, Thr490, and Thr655. Interestingly, for those 21 residues that each had four or more substitutions giving rise to LOF variants, the predictions from the Polyphen2 algorithm [16] were all predicted to be damaging, but the PolyPhen2 predictions for five of the seven residues with four or more variants with mixed functional scores (LOF and hyperactive) had Polyphen2 predictions of benign (Table S1).

Two other residues (Ser251 and Ser231) deserve discussion and are included among the residues with four or more variants that were either LOF or both LOF and hyperfunctional. Substitutions at Ser251 gave rise to three LOF variants, including the potentially phospho-mimetic variant S251E. The other phosphomimetic substitution at this residue, S251D was not included in the final table of missense variants because of high variation in measurements, but it was confirmed as LOF in a singleton assay (see below, Figure 3). Substitutions at the Ser231 residue gave rise to one LOF (S231P) and one borderline LOF phosphomimetic variant (S231E; score 0.924). The S231D potentially phosphomimetic variant was not scored. Importantly, two variants with non-phosphorylatable substitutions at the same site (S231G and S231N) scored as hyper-functional. Both Ser251 and Ser231, are on the surface of the protein and they are potential phosphorylation acceptors; we further describe these variants below with the singleton assays.

**Figure 3.**
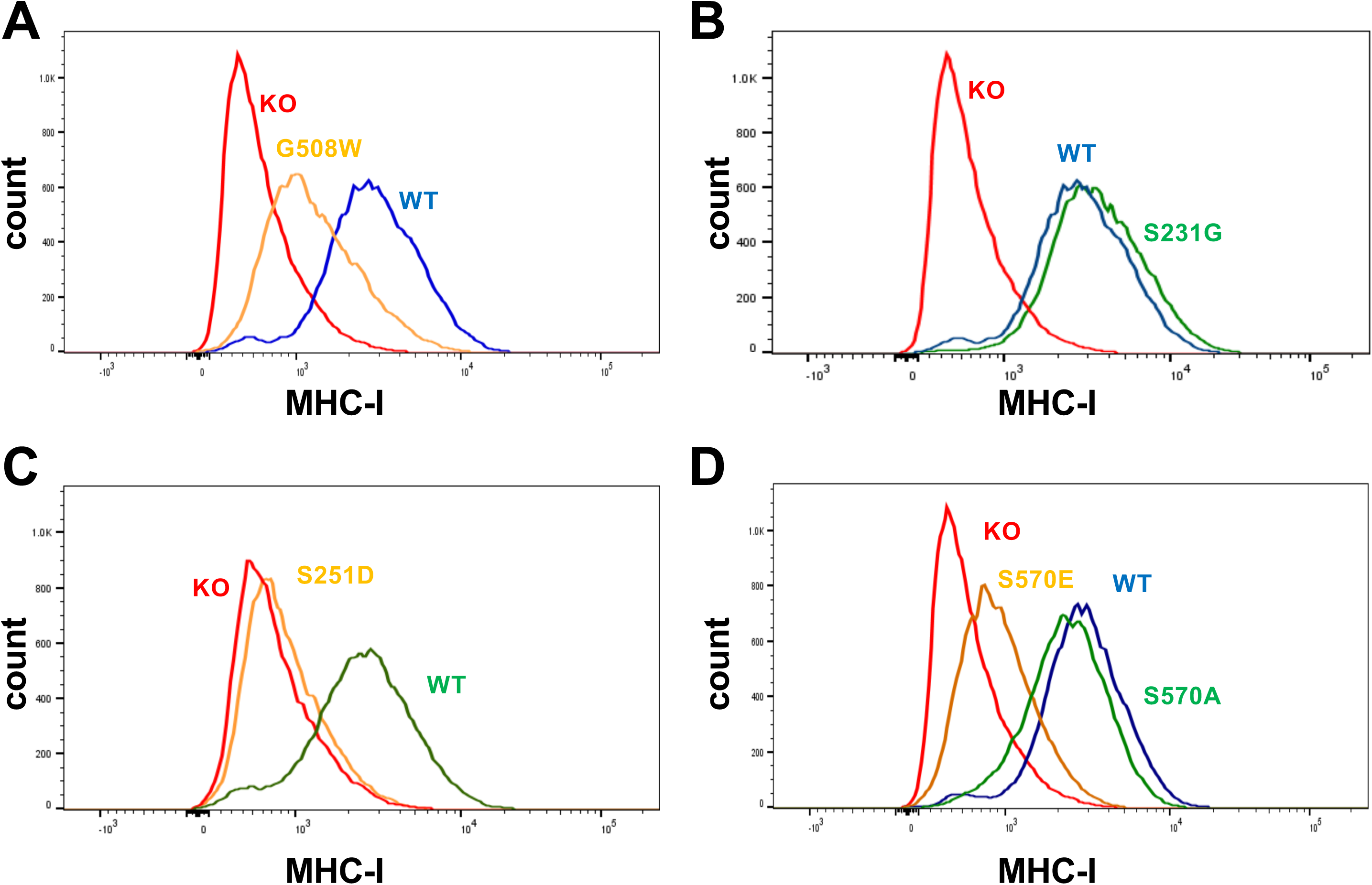
Results from selected singleton assays of TAP2 variants for function in regulating MHC-I cell surface abundance. **A.** U2OS TAP2-KO cells (KO) were rescued by recombination into a single genomic site of a plasmid encoding wild-type TAP2 (WT), or LOF variant TAP2-G508W. Cells were stained with antibody specific for HLA-A/B/C and subjected to flow cytometry. Results from three independent experiments were combined in the histogram, with MHC-I cell surface abundance (x-axis) and cell count (y-axis). **B.** U2OS TAP2-KO cells (KO) were rescued by recombination into a single genomic site of a plasmid encoding wild-type TAP2 (WT), or hyperfunctional variant TAP2-S231G and analyzed as in panel A. Results from three independent experiments were combined in the histogram, with MHC-I cell surface abundance (x-axis) and cell count (y-axis). The WT and KO results were taken from the same results as used in panel A. **C.** U2OS TAP2-KO cells were rescued with TAP2-S251D and measured for function in regulating the abundance of MHC-I on the cell surface. Results from three independent experiments were combined in the histogram. **D.** U2OS TAP2-KO cells were rescued with TAP2-S570E and TAP2-S570A and measured for function in regulating MHC-I abundance on the cell surface. Results from three independent experiments were combined in the histogram.

The structure of TAP in association with the Herpes simplex virus protein ICP47 has been determined with cryo-EM [7,8]. A ribbon diagram and a space filling model of the TAP heterodimer (Figure 2F and Figure S3, respectively) show how TAP1 (gray) and TAP2 (blue) interact. Residues whose substitution gave rise to four or greater LOF variants are shown in red, while residues whose substitution gave rise to both four or more LOF and hyperfunctional variants are shown in yellow. The surface exposed residues that have multiple LOF are primarily in the NTP binding domain or in the peptide channel. Several of these important residues were revealed by digitally removing TAP1 from the structure (Figure S3), indicating that these residues are at the TAP2 interface with TAP1. TAP is an ABC transporter that undergoes a major conformational shift in the presence of peptide cargo and ATP, and many of the residues highlighted by the functional analysis are at positions that would be expected to influence this allosteric conformational change.

### Singleton functional analyses of selected TAP2 variants confirmed their role in the regulation of MHC-I surface localization

Several variants, which were identified as LOF or hyperfunctional in the multiplexed screen, were re-tested in singleton assays. In the singleton assay, a missense substitution was cloned into the same promoterless plasmid used for the construction of the library (Figure S1) and transfected in the same genetically modified U2OS cells, which were used for the multiplex screen and tested one-at-a-time for effects on MHC-I surface abundance measured by flow cytometry. Since all the variants were recombined into the same genomic location with the same Dox-inducible RSV promoter, they were all transcribed at similar levels in all the transfected cells. First, we tested the NTP-binding pocket variant G508W, which was one of the variants with the lowest functional score (0.16) in the multiplex screen. The singleton assay showed that this variant dramatically downregulated MHC-I expression on the cell surface and was consistent with the results of the screen (Figure 3A). Given that the bulky tryptophan residue is likely to block ATP binding, this observation suggests that interfering with the binding of ATP profoundly impairs the TAP functional activity.

Subsequent singleton assays focused on the characterization of substitutions targeting potential phosphorylation sites. The variant S231G was among the hyperfunctional variants in the multiplexed screen, and its expression in the MHC-I-high U2OS cells, in the singleton assay, resulted in a slight increase of MHC-I on the cell surface (Figure 3B), confirming the results of the screen. These observations suggest that the Ser231 site may undergo phosphorylation and that its phosphorylation may functionally inhibit TAP. This was supported by the observation that the potentially phosphomimetic S231E substitution, gave rise to a borderline LOF TAP2 variant (functional score 0.924) in the multiplex screen.

As mentioned in the preceding section, a putative phosphomimetic substitution in Ser251 (S251E) gave rise to a LOF TAP2 variant in the multiplex screen. Here, we targeted the same TAP2 site, to generate another potentially phosphomimetic variant (S251D), which we functionally tested in a singleton assay in U2OS cells. The results confirmed the multiplex observation by showing a dramatic downregulation of MHC-I in the S251D-transduced cells (Figure 3C). Multiple substitutions at Ser251 would block phosphorylation, and these were functionally normal (Table S2). These data with the putative phospho-mimetic substitutions at Ser251 suggest that this residue may also undergo phosphorylation and that its phosphorylation might functionally inhibit TAP.

Three of the sites in TAP2 were reported to be phosphorylated in the proteomic data commons (https://proteomic.datacommons.cancer.gov/pdc/): Thr455, Thr458, and Ser570. The functional significance of their phosphorylation was addressed with the characterization of TAP2 variants carrying phosphomimetic or phosphoinhibitory substitutions at these sites. The phosphomimetic T455E variant was functionally normal (Table S2), suggesting that the phosphorylation of this site does not modulate the activity of TAP2. Both phosphomimetic variants at Thr458 were scored, and T458D was a LOF variant and T458E was functionally normal (Table S2). These data suggest that the phosphorylation of Thr458 modulates TAP2 function, however, because of the discrepancy of the T458D and T458E functional scores, we cannot determine whether the phosphorylation of this site stimulates or inhibits the function of TAP2.

The Ser570 variants were not included in the multiplex library of TAP2 variants. However, phosphomimetic and phosphoinhibitory substitutions at this site were tested in singleton assays in U2OS-TAP2-KO cells. The results showed that whereas the S570E missense variant dramatically downregulated the surface expression of MHC-I, the S570A variant was functionally normal (Figure 3D). The functional neutrality of the S570A phosphoinhibitory variant was not unexpected, given that the assay was performed in the U2OS cells, which have relatively high levels of expression of MHC-I on the cell surface (Figure S4A). These data suggest that TAP2 phosphorylation at Ser570 inhibits TAP activity.

### Candidate TAP2 kinases that when overexpressed could repress TAP2 function

Matching the amino acid sequences surrounding Ser570 and Ser251 with known kinase phosphorylation motifs [17], identified candidate TAP2 kinases that could potentially phosphorylate these sites. Using this strategy, we identified FAM20C [17] as a candidate Ser570 kinase and the PLK family of kinases (PLK1, PLK2, and PLK3) as candidate Ser251 kinases. If these kinases indeed phosphorylate these sites, their inhibition should increase the abundance of MHC-I on the cell surface. To address the effects of the inhibition of these kinases on MHC-I therefore, we chose to target them in two cancer cell lines (MCF7 and T47D) which express low levels of MHC-I on the cell surface (Figure S4A).

FAM20C is one of the most active kinases in the Golgi compartment [18] and is frequently overexpressed in tumors [19,20]. Knocking down FAM20C in T47D cells with three different shRNAs (Figure S4B), resulted in a 1.4-to 2.5-fold increase in the median value of MHC-I surface expression (Figure 4A), suggesting that this kinase indeed inhibits the TAP2/MHC-I axis.

**Figure 4.**
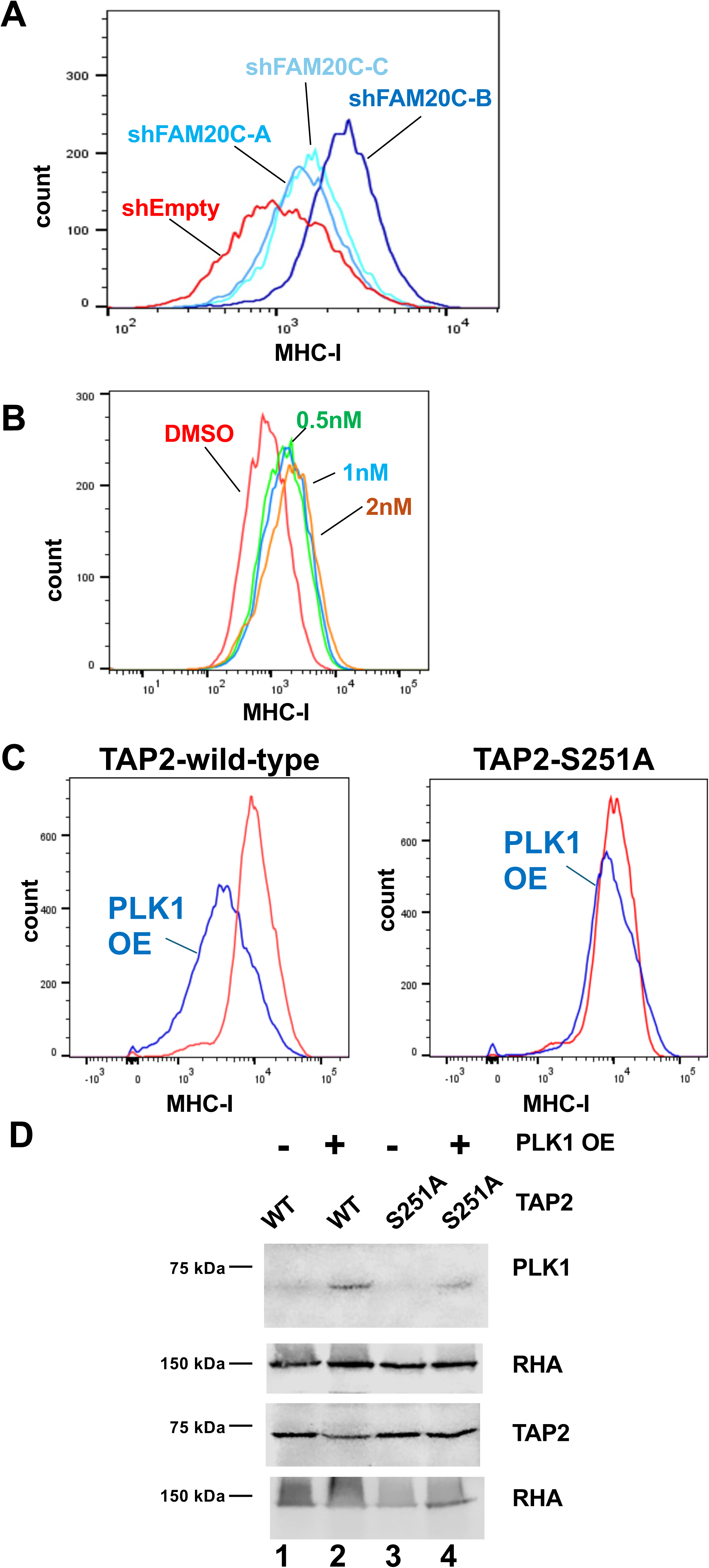
Kinases that potentially phosphorylate TAP2 affect MHC-I surface expression. **A.** The kinase FAM20C was depleted in T47D cells by stable expression of three different shRNAs targeting FAM20C, as indicated; cells were stained with anti-HLA-A/B/C antibody and analyzed by flow cytometry. **B.** MCF-7 cells were treated by the inclusion in the media of 0.5 nM BI-2536 (green), 1 nM BI-2536 (blue), 2 nM BI-2536 (brown), or vehicle control (DMSO, red). The abundance of MHC-I on the cell surface was measured five days later and scored by flow cytometry. **C.** U2OS TAP2-KO cells were rescued with TAP2-WT or TAP2-S251A. These two cell lines were used to overexpress PLK1. Expression of both, the PLK1 and the TAP2 variant, were induced by the inclusion of dox in the medium for six days followed by the measurement of the abundance of MHC-I at the cell surface using flow cytometry. Shown on the left are cells expressing TAP2-WT without PLK1 overexpression (red) and TAP2-WT with PLK1 overexpression (blue). Shown on the right are cells expressing TAP2-S251A without PLK1 overexpression (red), and TAP2-S251A with PLK1 overexpression (blue). Results are combined from three independent experiments. **D.** Protein samples from the cells in panel C were analyzed by immunoblot for the expression of PLK1 (*top*, lanes 2 and 4) and TAP2. Antibody specific for RNA Helicase A (RHA) was used as a loading control (*bottom*).

Among the kinases predicted [17] to phosphorylate Ser251 the PLK kinase family (PLK1, PLK2, and PLK3) scored highly. We treated MCF7 cells, which have relatively low concentration of MHC-I on the cell surface (Figure S4A) with the PLK inhibitor, BI-2536, and we observed a dose– and time-dependent increase in cell surface expression of MHC-I in the presence of this compound (Figure 4B, S4C). Using BI-2536 concentrations as low as 0.5 nM, maximum MHC-I upregulation was observed at five days. BI-2536 inhibits all three PLK isoforms, but at 0.5 nM it is likely to selectively target PLK1 whose IC50 falls in this concentration range. To determine whether the PLK kinases regulate MHC-I surface expression by phosphorylating TAP2 at Ser251, we overexpressed PLK1 in U2OS-TAP2-KO cells that had been rescued with either wild-type TAP2 or TAP2-S251A. Flow cytometric analysis revealed a dramatic downregulation of MHC-I surface expression in PLK1-transduced cells expressing wild type TAP2, but not in PLK1-transduced cells expressing TAP2-S251A (Figure 4C-D). This observation provides indirect evidence for the phosphorylation of TAP2 on Ser251, and strongly supports the hypothesis that PLK1 inhibits MHC-I surface expression by phosphorylating TAP2 at this site.

### Phosphorylation of TAP2 impacts the biology of human tumors

Phosphorylation of several sites of TAP2 have been detected in tumor samples in the Clinical Proteomic Tumor Analysis Consortium (CPTAC) database. To determine whether the level of phosphorylation of different TAP2 sites correlated with differences in the biology of the tumors, we compared the transcriptomes of tumors with high and low levels of TAP2 phosphorylation. The results showed that differences in the level of TAP2 phosphorylation at Thr458 in lung cancer correlated with differences in the expression of genes involved in the antigen presentation and cell cycle pathways (Figure 5A, B). Similarly, Thr458 phosphorylation detected in breast cancer (Figure 5C) and glioblastoma (Figure 5D) was correlated with changes in the antigen presentation pathway. Additionally, differences in the level of TAP2 phosphorylation at Ser570 in endometrial cancer also correlated with transcriptomic differences in the antigen presentation and cell cycle pathways (Figure 5E, F). Among the genes with high expression in the presence of TAP2 phosphorylation are: TAP1, TAP2, calreticulin, proteasome subunits, and multiple other genes in the antigen presentation pathway (Figure 5). The upregulation of cell cycle driving proteins indicated that the cells in these tumors with TAP2 phosphorylation had altered control of cell divisions. Given that the tumor transcriptome can be viewed as a surrogate of the tumor phenotype, this analysis revealed that phosphorylation of TAP2 on these residues affects the phenotype of the tumors.

**Figure 5.**
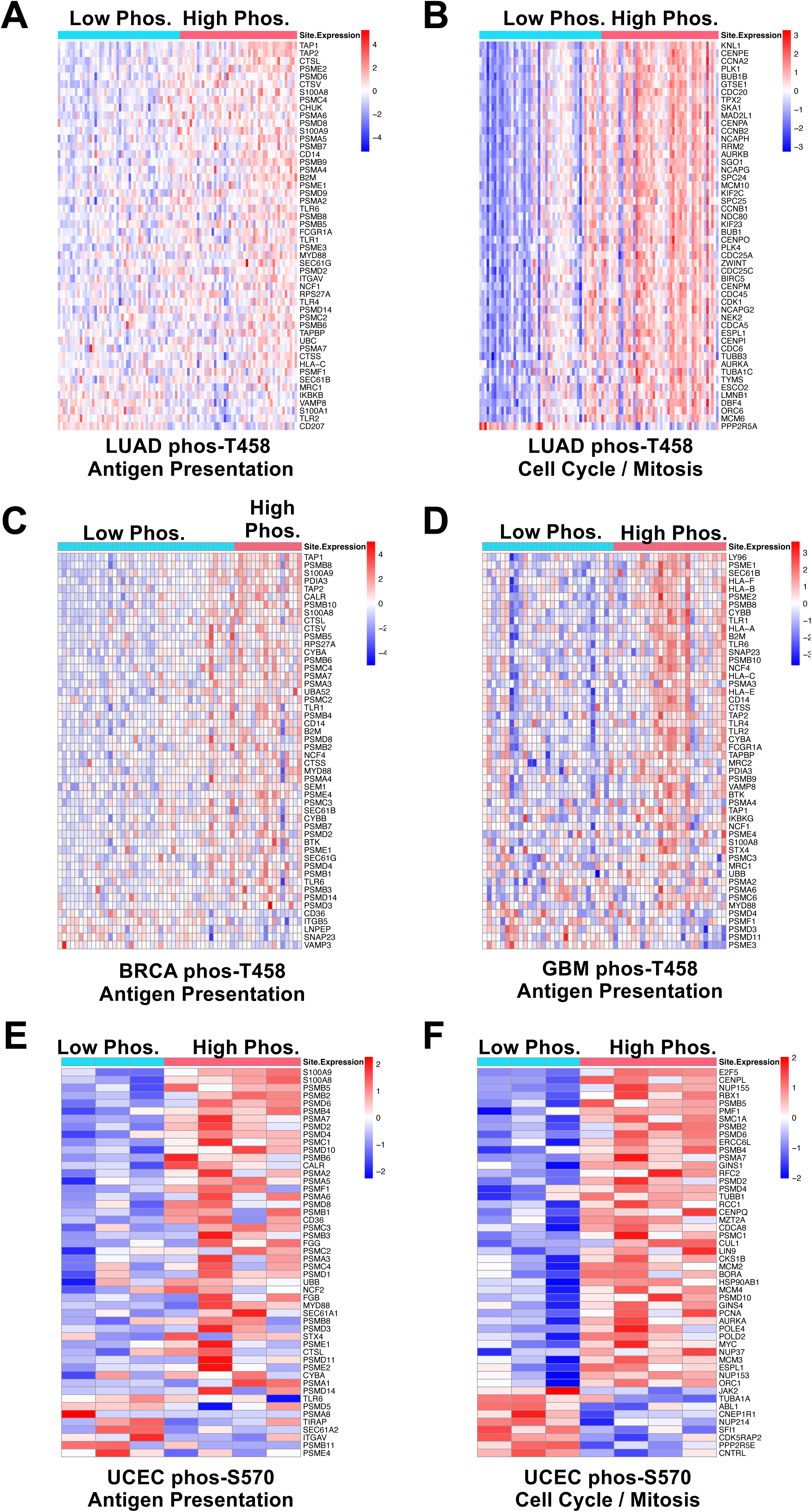
Phosphorylation of TAP2 in tumor samples was correlated with changes in pathways regulating antigen presentation and the cell cycle. **A.** Phosphorylation data in the CPTAC database include detected phosphorylations in tumors for which transcriptome data are available. In lung adenocarcinoma, phosphorylation of TAP2-Thr458 was correlated with an increase in RNA abundance in genes encoding antigen presentation (False discovery rate (FDR) = 0). In the heatmap, columns represent tumor samples with low, or undetectable, phosphorylation at the residue on the left, and high phosphorylation levels on the right. Rows represent individual genes in the pathway. **B.** Using the same lung cancer dataset as in panel A, phosphorylation of TAP2-Thr458 was correlated with an increase in RNA abundance from genes encoding proteins that regulate mitosis (FDR = 0). **C.** Phosphorylation data in the CPTAC database for a breast cancer tumor dataset with detected phosphorylation of TAP2-Thr458 are shown with transcriptome results. Tumor samples with low detected phosphorylation at Thr458 are represented by columns on the left, and samples with high detected phosphorylation at this residue are represented by columns on the right. The relative expression of genes in the antigen presentation pathway are shown. (FDR = 0.00166) **D.** Phosphorylation data in the CPTAC database for a glioblastoma tumor dataset with detected phosphorylation of TAP2-Thr458 are shown with transcriptome results. Tumor samples with low detected phosphorylation at Thr458 are represented by columns on the left, and samples with high detected phosphorylation at this residue are represented by columns on the right. The relative expression of genes in the antigen presentation pathway are shown. (FDR = 0) **E.** In an endometrial cancer dataset, phosphorylation of TAP2-Ser570 was correlated with increased expression of RNAs in the antigen presentation pathway (FDR=0). **F.** In the endometrial cancer dataset, phosphorylation of TAP2-Ser570 was correlated with increased expression of RNAs in the pathway regulating mitosis. (FDR=0)

## DISCUSSION

### Multiplex assay for function of TAP2

In this study, we employed a multiplexed assay to determine the effects of missense substitutions of TAP2 residues on the regulation of the abundance of MHC-I on the cell surface. TAP transports peptides from the cytoplasm to the ER, promoting the translocation of MHC-I to the cell membrane. We therefore targeted 96 codons for mutagenesis, and we have functional scores for 1252 missense variants, 106 synonymous variants, and 46 nonsense variants. Among the missense variants, 273 were classified as LOF.

We measured the effect of 10 VUS that were listed in ClinVar, and we found that one was a LOF variant and nine were functionally normal. The VUS with measured LOF variant was S219F, and we infer that it is likely pathogenic. We would predict that a germline TAP2-S219F would lead to severe immunodeficiency, and a somatic TAP2-S219F variant in a tumor cell would enable the tumor cell to evade immuno-surveillance. Interestingly, phosphorylation of Ser219 has been detected (PhosphositePlus website), but since thirteen of the fifteen variants at Ser219 scored in the functionally normal range, the results of the multiplex assay did not definitively address how the phosphorylation of this residue impacts TAP function.

The American College of Medical Genetics has developed tools for evaluating the functional results from in vitro assays with regard to pathogenicity of a variant [21], but to make these measurements, known pathogenic and benign variants must be available. For TAP2, there are few variants with clinical classifications. Two variants we assayed, R313H and T665A, were listed as benign in ClinVar, and they scored in the functionally normal range in the multiplexed assay used in this study. The variant R273* is listed as pathogenic in ClinVar, and it was LOF in this assay. That the two known benign variants of TAP2 were scored as functionally normal and the one known pathogenic variant was scored as LOF was encouraging, but we cannot assess the accuracy of the functional test based on these three variants.

Twenty-one TAP2 residues had four or more variants with scores indicating LOF. These residues clustered in the NTP binding domain and on TAP2 surfaces that interface with TAP1 in the transmembrane domain. Both, TAP1 and TAP2 have NTP binding domains that are spatially separated in the absence of bound ATP. When both are bound to ATP, the two globular domains bind and switch the conformation of TAP from cytoplasmic facing in the unbound state to ER lumen facing in the ATP-bound state. Since many LOF variants had substitutions in the NTP binding domain or at residues along the TAP1-TAP2 interface, we suggest that many of these LOF variants affect the allosteric change of TAP, which is required for the transport of peptides across the ER membrane into the ER lumen. Importantly, the function of these variants is consistent with the results of earlier studies addressing the mechanisms by which proteins encoded by several members of the herpes virus family enable the virus infected cell to evade the host immunosurveillance. These studies had shown that the HSV1 ICP47 protein binds the peptide binding site of TAP2 and, along with blocking peptide binding, the ICP47 freezes the TAP conformation in the cytoplasmic facing orientation [22]. Proteins encoded by other members of the herpes virus family include US6, BNLF2a, CPXV012, and UL49.5 which all appear to block the ATP binding domain of TAP [22]. Therefore, many of the LOF variants in the current study have substitutions at amino acid residues in TAP2 domains targeted by herpes virus proteins that enable these viruses to inhibit MHC-I expression on the cell surface and antigen presentation.

### Regulation of TAP2 by post-translational modification (PTM)

Multiple sites of ubiquitination and phosphorylation are listed in the online databases PhosphositePlus and Protein Data Commons. Although these PTMs have been detected and listed in publicly available databases, there is essentially nothing known about their functional impact. The functional screen in this study presents, for the first time, the opportunity to address the important question of the role of the PTMs on the regulation of TAP activity in the antigen presentation process.

Since polyubiquitination of lysine often targets a protein for proteasome-mediated degradation, we anticipated that missense substitutions of lysines that are acceptors for ubiquitin would give rise to more stable, hyperactive variants. Eleven TAP2 lysines that were known ubiquitination acceptors were targeted in the multiplex library, and only one of these, Lys356 (K356A) was scored as hyperactive. Overall, the functionality of the lysine-substituted variants suggests that lysine ubiquitination may not target TAP2 for degradation. It is indeed known that proteasomal degradation is only one of the potential outcomes of lysine ubiquitination and that different forms of this modification may be recognized by signaling proteins in many signaling pathways.

Alternatively, it is possible that any of several lysine residues, would suffice to mark the protein for degradation following ubiquitination; in this study we only make one missense substitution per molecule and would miss the effect on protein stability if multiple lysine acceptors needed to be changed to stabilize the protein.

Our multiplex assay identified substitutions indicative of phosphomimetic changes that resulted in LOF, suggesting that phosphorylation of these sites would functionally inactivate TAP2, lowering the abundance of MHC-I on the cell surface and inhibiting antigen presentation. The eleven possible serine and threonine phosphomimetic sites are shown in Figure S5A, and these localize primarily to the NTP-binding domain and to the cytoplasmic portion of the transmembrane domain. There are five observed phosphorylation sites in TAP2, and four of these cluster in the NTP binding domain and one is in the transmembrane domain (Figure S5B). Of the five previously observed phosphorylation sites, only two were potentially LOF in this analysis. Clearly, since the eleven LOF serine and threonine residues are potential phosphomimetic sites, some may in fact be phosphorylated in a condition-specific manner but not previously observed. An excellent candidate for phosphorylation is the site at Ser251. Both phosphomimetic changes at Ser251 (S251D and S251E) were LOF. The amino acid motif surrounding the phosphorylatable residue was analyzed using an online database [17], and the top hits for Ser251 were PLK and COT. We tested inhibitors against these two kinases, and we found that the inhibitor specific for the PLK did result in an increase in MHC-I abundance in MCF7 cells. This inhibitor had previously been found to increase MHC-I abundance in other cell lines [23]. We found that the effective concentration of the chemical inhibitor was consistent with inhibition of PLK1, and overexpression of PLK1 did downregulate MHC-I surface abundance in cells expressing wild-type TAP2, but cells expressing only TAP2-S251A were resistant to the downregulation by PLK1 expression. These results strongly suggest that when PLK1 is overexpressed, it phosphorylates proteins that are not normal substrates for the kinase, and this includes TAP2-Ser251. We conclude that PLK1 phosphorylation of TAP2-Ser251 downregulates its activity, reducing the availability of antigenic peptides in the endoplasmic reticulum, and a reduction in MHC-I expression on the cell surface. PLK1 is overexpressed in many tumors, and an informatics analysis suggested that the overexpression of PLK1 could suppress MHC-I signaling [23].

Similarly, in T47D cells, which we found express low amounts of MHC-I on the cell surface, inhibition of FAM20C, the putative kinase that can phosphorylate Ser570, resulted in an increase in MHC-I surface expression. The FAM20C kinase is overexpressed in many tumor types [18,19], consistent with the notion that the overexpression of the FAM20C kinase in a tumor cell could decrease TAP2 activity with consequent downregulation of the antigen presentation process and evasion of the host immune surveillance.

If the apparent regulation of the abundance of MHC-I on the cell surface by TAP2 phosphorylation is clinically relevant, one would expect that differential phosphorylation of TAP2 sites in different types of human cancer would affect antigen presentation and other cancer relevant functions. This important question was addressed with bioinformatic analyses correlating the levels of TAP2 phosphorylation at Thr458 and Ser570 with the transcriptome in individual tumors of several tumor types (Lung adenocarcinomas, Breast Cancer, Glioblastoma and Endometrial Cancer). These analyses revealed significant concordance between phosphorylation at these sites and transcriptomic changes in the antigen presentation and cell cycle pathways, suggesting that their phosphorylation may be clinically relevant. The changes in the cell cycle could suggest that tumors with phosphorylated TAP2 are more aggressive.

Tumors evolve over time to counter the challenges they meet in an unfavorable microenvironment. One of the challenges is anti-tumor immunity. Cells succeeding to evade the anti-tumor immune response are positively selected. One mechanism commonly employed by the tumor cells to avoid anti-tumor immunity is the downregulation of MHC-I surface expression, which diminishes antigen presentation, and the T cell mediated host defense. However, if MHC-I surface expression is too low, the cancer cells may become vulnerable to NK cell-mediated lysis [1]. Achieving a low level of expression of MHC-I sufficient to evade the T cell cytolysis but sufficiently high to avoid lysis by NK cells could be achieved by multiple mechanisms [1]. To survive both these host defense mechanisms therefore, the cancer cells need to downregulate MHC-I on the cell surface to levels low enough to diminish recognition by T cells, but not too low, to elicit NK cell-mediated lysis. This can be achieved by mechanisms that regulate the expression and activity of the components of the peptide transporter TAP and the peptide loading complex (PLC). Regulation of TAP2 phosphorylation, which is addressed in this report, is one of these mechanisms. By taking advantage of the power of gene variant screening to explore the molecular mechanisms by which the activity of TAP2 in the antigen presentation pathway is regulated, this study identified Ser251 phosphorylation as a regulator of anti-tumor immunity and as a new potential precision biomarker for cancer immunotherapy. Additionally, it identified PLK1 inhibitors as potential novel immunotherapeutic agents.

## METHODS

### Plasmids and viral vectors

The plasmid for expressing wild-type TAP2 and TAP2 variants was constructed from pUC19. The pUC-attB-HygroR-P2A-TAP2 plasmid had no eukaryotic promoter and included the following elements: attB sequence, hygromycin resistance, P2A translation re-start sequence, TAP2, and the bGH polyadenylation sequence (Figure S2A). When this plasmid was recombined into cells containing integrated pLenti-TetBxb1BFP-Int-Blast [10], the hygromycin resistance and TAP2 would be expressed when 10 µg/ml Dox was included in the medium. The TAP2 gene sequence contained silent mutations that rendered it resistant to CRISPR. Details of the plasmid construction, its sequence, and variants generated by site-directed-mutagenesis are available from the authors. The sequences of all plasmids used in this study were confirmed by Sanger sequencing using the OSU Comprehensive Cancer Center Genomics Shared Resource.

### Cell lines

The U2OS cell line (ATCC #HTB-96) was modified to incorporate in its genome a single attP recombination site by infecting cells with the virus encoded by the shuttle plasmid pLenti-TetBxb1BFP-Int-Blast [10]. Cells were colony selected in 15 µg/ml blasticidin and 10 µg/ml dox. Colonies grown from single cells were tested for a single integration site by transfecting both pUC19-attB-PuroR-2A-dsRED and pUC19-attB-PuroR-2A-eGFP, selected in 1.5 µg/ml puromycin, and then analyzed by flow cytometry. Acceptable colonies had cells that were either dsRED-positive or eGFP-positive but not both, indicating a single integration site. The appropriate cell clone, called U2OS-G310a, was infected with LentiCas9Blast, a Cas9-expressing lentivirus [24] and selected in 15 µg/ml blasticidin in the absence of dox. U2OS-G310a-Cas9 cells were then infected with a lentivirus from pLentiguide-DKO [25] containing two sgRNAs targeting TAP2 and cells were selected in 1.5 µg/ml puromycin. Two weeks after selection, cells were stained for MHC-I on the cell surface, and cells with low abundance stain were collected using an ARIA III cell sorter in the OSU Comprehensive Cancer Center Flow Cytometry Shared Resource. These cells, now called U2OS-TAP2-KO were confirmed to stably have low concentration of MHC-I on the cell surface following repeated analysis by flow cytometry. Transfection of U2OS-TAP2-KO cells with plasmid pUC-attB-HygroR-2A-TAP2-wt in the presence of 10 µg/ml dox to induce expression of the Bxb1 integrase, resulted in the cell line U2OS-TAP2-WT with rescued expression of MHC-I on the cell surface (Figure 2).

### Generation of a library of variants in TAP2

The plasmid, pUC-attB-HygroR-2A-TAP2-wt, was subjected to inverse PCR using a strategy [14] to generate degenerate sequences at the targeted codons of TAP2. In brief, the forward primer had NNK sequence (where N = A, C, G, T and K = G, T) for the targeted codon and the next 27 bases were the wild-type sequences of the nine codons downstream of the targeted codon. The reverse primer had wild-type sequence of the reverse-complement sequence of the ten codons upstream of the targeted codon. Ninety-six primer pairs used in 96 separate PCR reactions using high fidelity polymerase copied the entire pUC-attB-HygroR-2A-TAP2 plasmid with the targeted codon at the 5’ end and the upstream sequences at the other end of the plasmid. Following quantitation of PCR products by agarose gel electrophoresis, the 96 PCR reactions were mixed to obtain equimolar DNAs from each PCR product. The PCR product was subjected to DpnI digest to cleave template DNA, phosphorylation with T4 PNK (New England Biolabs) and ligated with T4 DNA Ligase (New England Biolabs). Ligated DNA was transformed into DH5alpha cells (New England Biolabs) and 9,200 colonies were obtained. Barcodes were added to the library DNA by digesting the plasmid with XhoI and NsiI, each of which had a single cut site in the DNA sequence of the 3’-UTR, barcode oligonucleotides (Table S6) were annealed and ligated into the DNA, and > 10,000 colonies were obtained. The plasmid library was sequenced, and the 18 bp barcode was linked to each variant using PacBio long-read sequencing service at the Drexel University Sequencing Core.

### Integration of TAP2 library of variants into U2OS-TAP2-KO cells

U2OS-TAP2-KO cells were cultured in 10 µg/ml doxycycline overnight before transfection. Cells were plated in a 6-well plate such that they would be ∼80% confluent for transfection. Each well of the 6-well plate received 1 ml normal media without pen/strep with 10 µg/ml Dox for the duration of the transfection. Transfections were left on cells for approximately 24 hours. Each transfection consisted of 500 ng TAP2 library and 5 µl lipofectamine-2000 (Thermo-Fisher). Two wells of pUC19-AttB-PuroR-2A-eGFP plasmid, one well with dox and one well without, were transfected simultaneously with the TAP2mut variant library. Approximately 48 hours after transfection, before any antibiotic selection, the wells containing eGFP expressing cells were collected and analyzed by flow cytometry to obtain the percentage of cells that were fluorescent.

The wells of library transfection were collected, cell count was taken, and then calculated a number of integrated cells based on the eGFP results. This was then divided by 2^3^, which conservatively assumes a doubling time for U2OS at under 20 hours. The replicate would continue as long as this number was >100,000 estimated integrated cells. Hygromycin (50 µg/ml) selection then began for the U2OS.

### Staining for MHC-I Surface Expression

Tissue Culture cells were removed from plates using TrypleE (Thermo-Fisher) followed by resuspending cells in medium containing 10% fetal bovine serum. Cell counts were normalized to 1 × 10^6^ cells per sample and resuspended in 200 µl sorting buffer (PBS, 6% fetal bovine serum, 25 mM HEPES, pH 7.2, 5 mM EDTA) plus 5% BSA, followed by rotation at room temperature for 15 minutes. Cells were collected by centrifugation (1000 rpm, 5 minutes) and the pellet was resuspended in 200 µl of sorting buffer with 1 µl of FITC-conjugated Anti-Hu HLA-ABC Ab (Thermo-Fisher, 11-9983-42) per stained sample. Samples were rotated for 30 minutes at room temp, followed by three washes using sorting buffer. After the third wash, samples were resuspended in 1 mL sorting buffer and fluorescence was measured on either Fortessa or ARIA III flow cytometers in the OSU Comprehensive Cancer Center Flow Cytometry Shared Resource. Flow cytometry experiments all included a wild-type TAP2 sample, a TAP2 KO sample, and an unstained sample.

### Multiplexed assay for function of TAP2

The plasmid containing the library of TAP2 variants was based on pUC19; it had no eukaryotic promoter but did contain downstream of the attB recombination site the hygromycin resistance gene, the P2A translation re-start sequence, the TAP2 gene, a barcode in the 3’UTR, and a polyadenylation sequence. When recombined in the single attP site in the genome, the hygromycin and TAP2 coding sequences were downstream of the RSV-LTR and the Tet-operator sequences. For the four multiplexed assays, we used four separate transfections of the multiplex library, and each transfection yielded > 100,000 independent integrations/recombinations. For each multiplex assay, 10 × 10^6^ cells grown in the presence of 10 µg/ml Dox were collected and stained for cell surface HLA-A/B/C. Cells were subjected to flow sorting using the Aria III cytometer in the OSU Comprehensive Cancer Center Flow Cytometry Shared Resource and four quartiles of cells were collected based on the intensity of antibody stain; 2 × 10^6^ cells were collected in each quartile. The genomic DNA was isolated from the cells in each quartile using the DNeasy kit (Qiagen), and the barcodes were recovered by two rounds of PCR using Kapa2G Robust Polymerase (Thermo-Fisher). The first round of PCR for 18 cycles using Nest1 primers (1:1 mix of Nest1-Fwd-N1 and Nest1-Fwd-N2 and a 1:1 mix of Nest1-Rev-N1 and Nest1-Rev-N2; Table S6). These Nest1 primers were a mixture of two primers, one with a single N, and one with two Ns as the first two bases that would be read using the Illumina sequencing; these bases were inserted to increase cluster identification in NGS sequencing. PCR products were purified using AmpurePB beads (PacBio) according to manufacturer’s instructions and were then subjected to a second round (8 cycles) with indexing primers (Table S6). PCR products were gel purified, quantified using Qubit, and samples were prepared as per the instructions of the sequencing facility (Memorial Sloan-Kettering Comprehensive Cancer Center Sequencing core).

### Variant scoring

The plasmid library containing TAP2 variants and linked barcodes were processed using Enrich2 software [26] and using algorithms described previously [27–29]. To measure the effect on function of TAP2 variants, the abundance of each variant in each quartile from the flow sorting was determined from the abundance of the linked barcodes in the samples. A read count was calculated for each variant for each replicate, with the read count cutoff representing the sum of the reads of a variant in the input sample plus the four quartiles for a given replicate. A read count cutoff of 250 was applied to the dataset. If the read count of a variant was above the cutoff for all four replicates, the functional score for each variant was determined based on the formula: score = log2[(Q4 / (Q1 + Q2 + Q3 + Q4)) / (Q1 / (Q1 + Q2 + Q3 + Q))], that is to say, the log base two of the percentage of quartile reads of a given variant in the fourth quartile divided by the percentage of quartile reads of that variant in the first quartile. After calculating the score for each variant, the variance of the scores from the four replicates was calculated, and any variant with a variance greater than 3.0 was excluded from the final dataset.

All algorithms used are available on the GitHub site: https://github.com/dongjunchung/parvin2024.

### shRNA-Mediated Knockdown of FAM20C

Three shRNA constructs targeting FAM20C were obtained from Sigma-Aldrich’s MISSION® shRNA library: TRCN0000128164, TRCN0000128300, and TRCN0000128442. The shRNA constructs were packaged into lentiviral particles using the second-generation lentiviral packaging system. HEK293T cells were transfected with the shRNA plasmids along with the psPAX2 packaging plasmid and pMD2.G envelope plasmid using Lipofectamine 3000 (Invitrogen) according to the manufacturer’s protocol. The viral supernatants were collected 48 hours post-transfection, filtered through a 0.45 µm filter, and used to transduce the target cells in the presence of 8 µg/ml Polybrene. T47D cells were infected with these three viruses carrying shRNAs targeting FAM20C as well as a lentivirus preparation that did not contain an encoded shRNA. T47D cells were selected in puromycin; stably integrated cells were analyzed for FAM20C mRNA expression by qRT-PCR, using standard procedures and were analyzed for expression of MHC-I on the cell surface by flow cytometry as described above.

### Overexpression of PLK1

The human PLK1 gene was cloned into the pInducer20 vector (Addgene plasmid #44012) [30] using the Gateway® LR Clonase® II Enzyme Mix (Invitrogen). The pDONR223-PLK1 plasmid (Addgene plasmid #23867) served as the entry clone containing the PLK1 insert. The lentivirus was infected into U2OS-TAP2-KO cells rescued with either wild-type TAP2 or TAP2-S251A. Infected cells were selected by inclusion of 125 µg/ml G418 in tissue culture medium.

Expression of both the PLK1 and the TAP2 protein were induced in the presence of Dox, and six days later, cells were stained for cell surface MHC-I and analyzed by flow cytometry, and cell lysates were analyzed by immunoblot with an antibody specific for PLK1 (ThermoFisher) using standard methods.

## ACKNOWLEDGMENTS

Thank you to the genomics and flow cytometry cores (P30CA016058) at the Ohio State University Comprehensive Cancer Center. This work was funded in part by NIH R01 CA228083 to JDP and a Pelotonia Fellowship to GN.

## Legends to Supplementary Figures

**Figure S1: Constructs used to express the TAP2 multiplexed library.**

The U2OS cell line had inserted at a single genomic locus the Landing Pad [10] containing the RSV LTR upstream of a Tet-operator sequence, the attP recombination sequence, blue fluorescent protein (BFP), the Bxb1 integrase (Int) and blasticidin resistance gene (Bsd) as a single mRNA. A CMV promoter drove the constitutive expression of the Tet-repressor protein. The TAP2 plasmid was based on a pUC19 backbone with an insertion including the attB recombination sequence followed by the coding sequence for hygromycin resistance, an in-frame P2A translation re-start sequence and an in-frame human TAP2 sequence with its stop codon. In the 3’UTR the 18 bp barcode was inserted, and the bGH polyadenylation sequence from the pcDNA3 plasmid was inserted downstream. This plasmid lacked a promoter for eukaryotic cells. Recombination mediated by the Bxb1 protein via the attB and attP sequences resulted in the insertion of the plasmid library downstream of the RSV-LTR and Tet-operator.

**Figure S2. The TAP2 multiplexed library**.

The TAP2 coding sequence is diagrammed for codons 2-344 (*top*) and 345-687 (*bottom*) with the wild-type residue indicated by the dark purple rectangles. At each position, possible missense substitutions are listed in the vertical axis. Nonsense mutations are indicated in the top row by the asterisk. Codons targeted for substitution are indicated by black ovals at the top of the matrix. Variants detected using long-read sequencing are indicated by the color from low number (white) to higher abundance of each variant in the pool indicated by blue and violet. Grey indicates substitutions not targeted and also missense substitutions that were targeted but not present in the library. Of the possible 1920 variants (19 missense substitutions and 1 nonsense substitution at 96 codons), 93% were present in the plasmid library.

**Figure S3. Evaluation of TAP2 missense variants for protein surface exposure**.

Four views of TAP are shown to evaluate the surface exposure of residues that had multiple LOF variants (drawn with PyMOL). The cryo-EM structure [7] of TAP1 (gray) in complex with TAP2 (blue) is shown. Residues for which four or more substitutions resulted in LOF are indicated in red and residues for which four or more substitutions resulted in either LOF or hyperfunctional changes are indicated in yellow. The full complex is shown (*top left*) or rotated (*top right*). The two views are shown on the bottom with the TAP1 protein digitally removed.

**Figure S4. Effects of kinase inhibition on MHC-I cell surface expression**.

**A.** Some commonly used cancer-derived cell lines were assayed for the abundance of MHC-I on the cell surface. Shown are the results for U2OS, which has a comparatively high level of MHC-I surface expression, along with breast cancer cell lines T47D and MCF7. Shown are three independent experiments combined in a single histogram.

**B.** Evaluation of depletion of FAM20C RNA from T47D cells expressing shRNAs targeting FAM20C. The effect of shRNAs targeting FAM20C in T47D cells was evaluated by qRT-PCR. The shFAM20C-768, shFAM20C-1488, and shFAM20C-1498 were shRNAs A, B, and C, respectively, in Figure 4A.

**C.** Time course of changes in MHC-I cell surface expression on MCF7 cells after treatment with the BI-2536 PLK inhibitor for the indicated times. Cells were treated with vehicle control (red), 1 mM (blue), or 2 mM (brown) BI-2536.

**Figure S5. Locations of LOF phosphomimetic substitutions on the TAP protein structure**.

A. The locations on the TAP protein structure of the LOF phosphomimetic substitutions are shown (red). The structure is rotated 180° at the right. These figures were drawn with PyMOL.

B. The TAP1 (gray) and TAP2 (blue) proteins are shown with phosphorylation sites that had been observed in publicly available databases (PhosphositePlus and Proteomic Data Commons) are shown (yellow). The globular NTP binding domain and the transmembrane domain are indicated. The structure is rotated 180° at the right.

## LEGENDS TO SUPPLEMENTARY TABLES

**Table S1. Codon selections**.

TAP2 codons targeted for mutagenesis are listed along with known VUS, known interactions or PTMs, conservation among 10 species (human, rat, mouse, chicken, chimpanzee, golden hamster, dog, gorilla, wild boar, and platypus), and domain localizations. Results from PolyPhen-2 [16] for substitutions of indicated residue to alanine or of known VUS are shown with likely damaging changes (red), possibly damaging substitutions (pink or yellow) or likely benign substitution (green).

**Table S2. Functional results of missense substitutions**.

Results for function in regulation of MHC-I cell surface localization are given for 1252 missense substitutions in TAP2. The functional scores from four separate transfections and integration into the U2OS-TAP2-KO cells are indicated, and the average score is highlighted (yellow).

Variants classified as loss of function (LOF) are highlighted in orange, and variants classified as hyperactive are highlighted in blue. Possible phosphomimetic substitutions are highlighted in gray. Read count constitutes the sum of read counts from the four quartiles plus an input library sample.

**Table S3. Functional results of synonymous substitutions**

Results for function in regulation of MHC-I cell surface localization are given for 106 synonymous substitutions in TAP2. As in Table S2, functional scores for variants in four replicate experiments are given, and the average functional score is highlighted in yellow.

**Table S4. Functional results of nonsense substitutions**

Results for function in regulation of MHC-I cell surface localization are given for 46 nonsense variants in TAP2. As in Table S2, functional scores for variants in four replicate experiments are given, and the average functional score is highlighted in yellow.

**Table S5. Variants of Unknown Significance**

Twelve codons with known VUS that were listed in ClinVar are indicated with the functional impact of the missense substitution. Also listed are two known benign variants and one pathogenic variant from ClinVar and the respective functional scores and one variant that was reported in literature as possibly cancer associated.

**Table S6. DNA primers used**

Table S6 lists the primers used for constructs and their sequences. Mutagenic oligonucleotides for multiplex libraries and for singleton experiments are available from the authors.

