## Supplemental Figures for "Multiplexed functional analysis of TAP2 variants in regulating MHC-I cell surface abundance reveals overexpression of PLK1 downregulates antigen presentation"

Figure S1

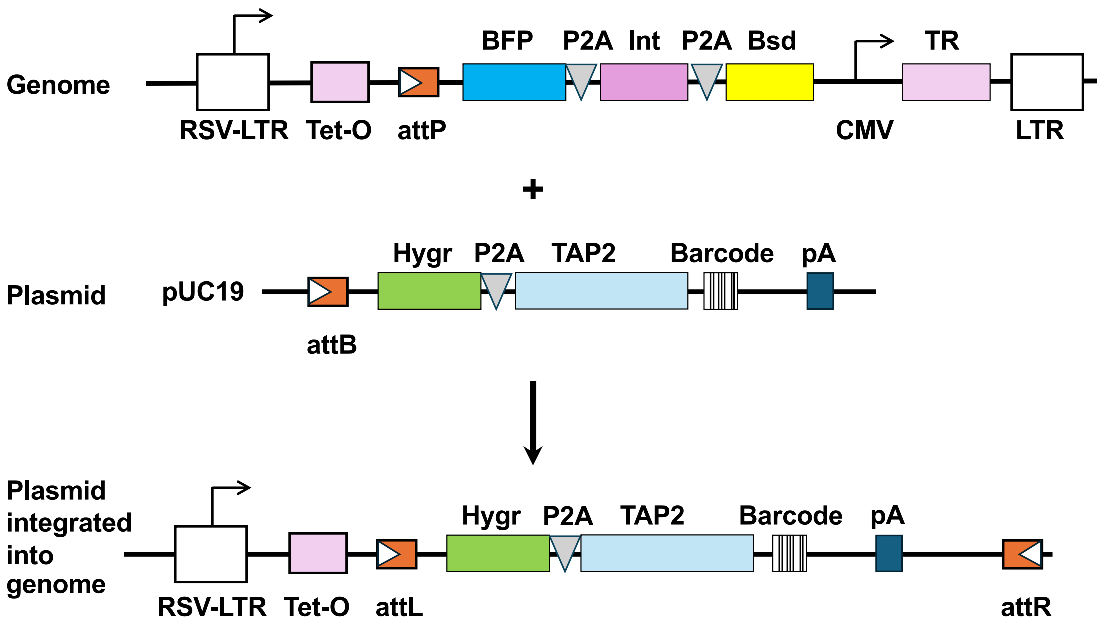

Genome
BFP
P2A
Int
P2A
Bsd
TR
RSV-LTR
Tet-O
attP
CMV
LTR
+
Plasmid
pUC19
Hygr
P2A
TAP2
Barcode
pA
attB
Plasmid integrated into genome
Hygr
P2A
TAP2
Barcode
pA
RSV-LTR
Tet-O
attL
attR

#### Figure S2

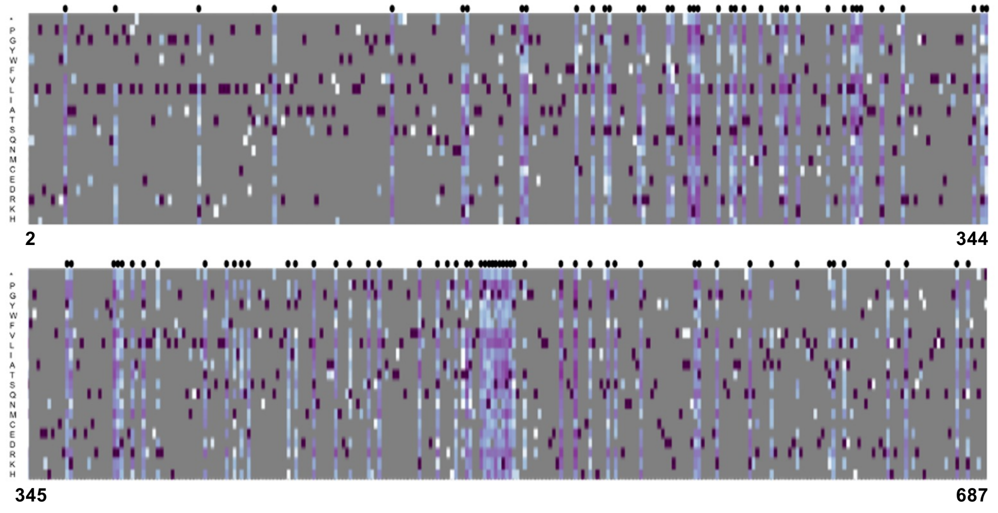

**Figure S3**

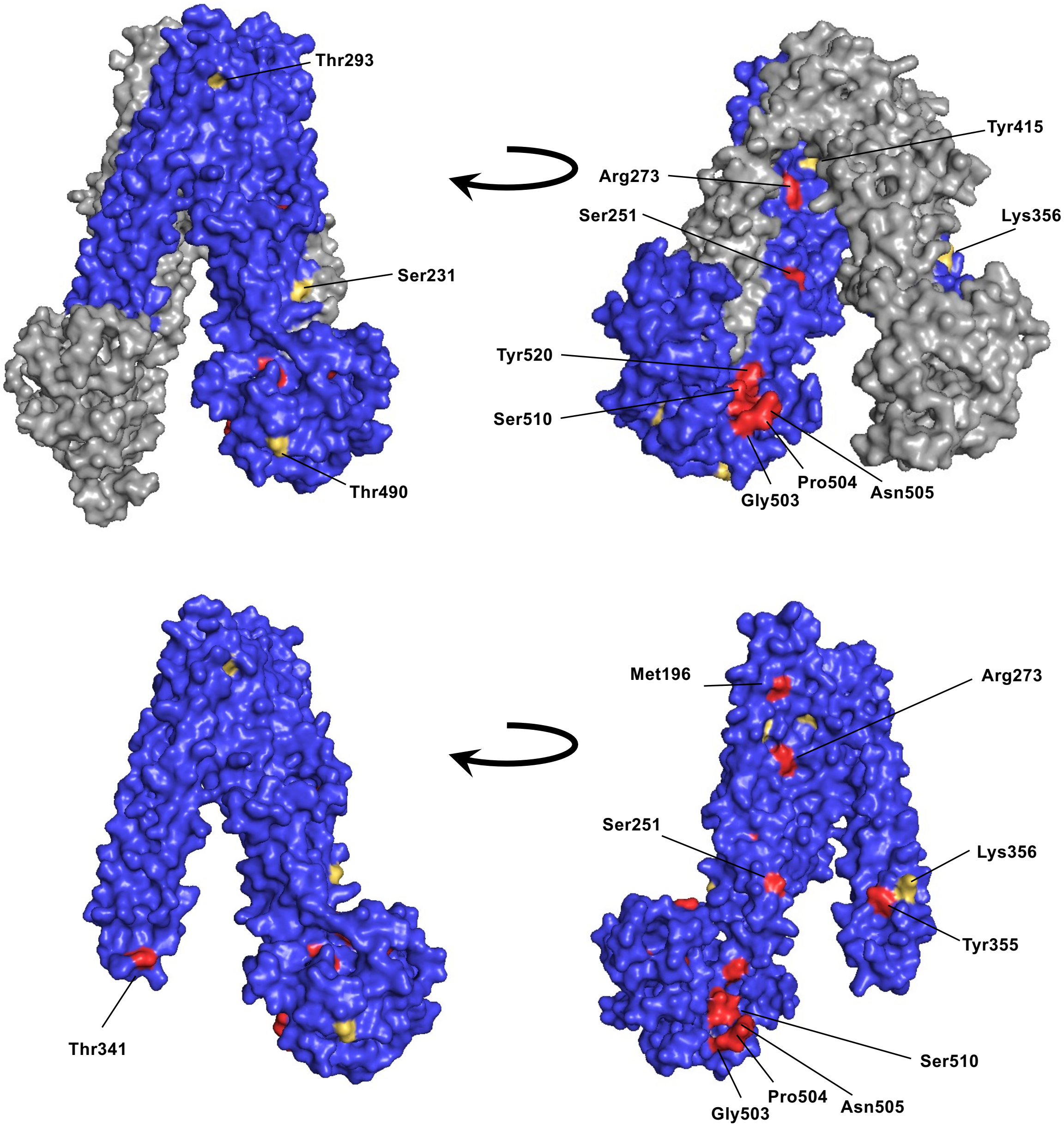
Thr293
Ser231
Thr490
Tyr415
Arg273
Ser251
Lys356
Tyr520
Ser510
Pro504
Asn505
Gly503
Thr341
Met196
Arg273
Ser251
Lys356
Tyr355
Ser510
Pro504
Asn505
Gly503

#### Figure S4

A

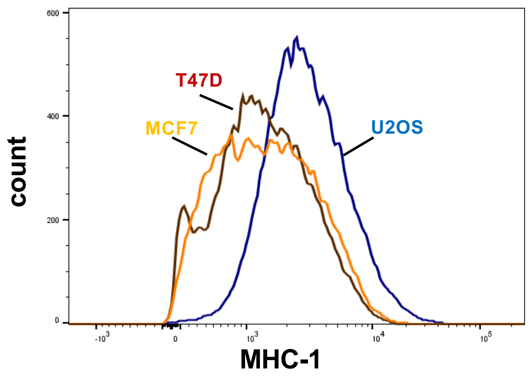


B

| | Average | St. Dev. |
|---|---|---|
| Control | 1 | 0 |
| shFAM20C-768 | 0.103 | 0.047 |
| shFAM20C-1488 | 0.252 | 0.030 |
| shRAM20C-1498 | 0.240 | 0.131 |

C

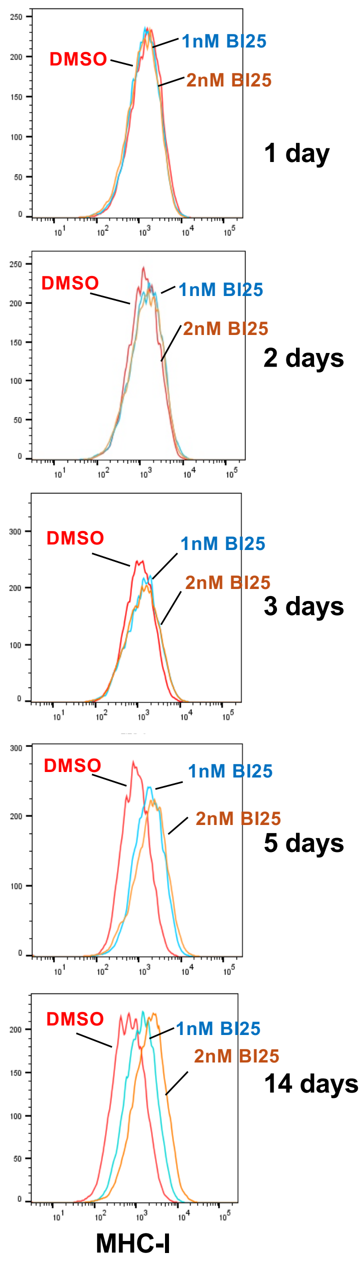


D

### Figure S5

**A** **TAP2 LOF phosphomimetic sites**

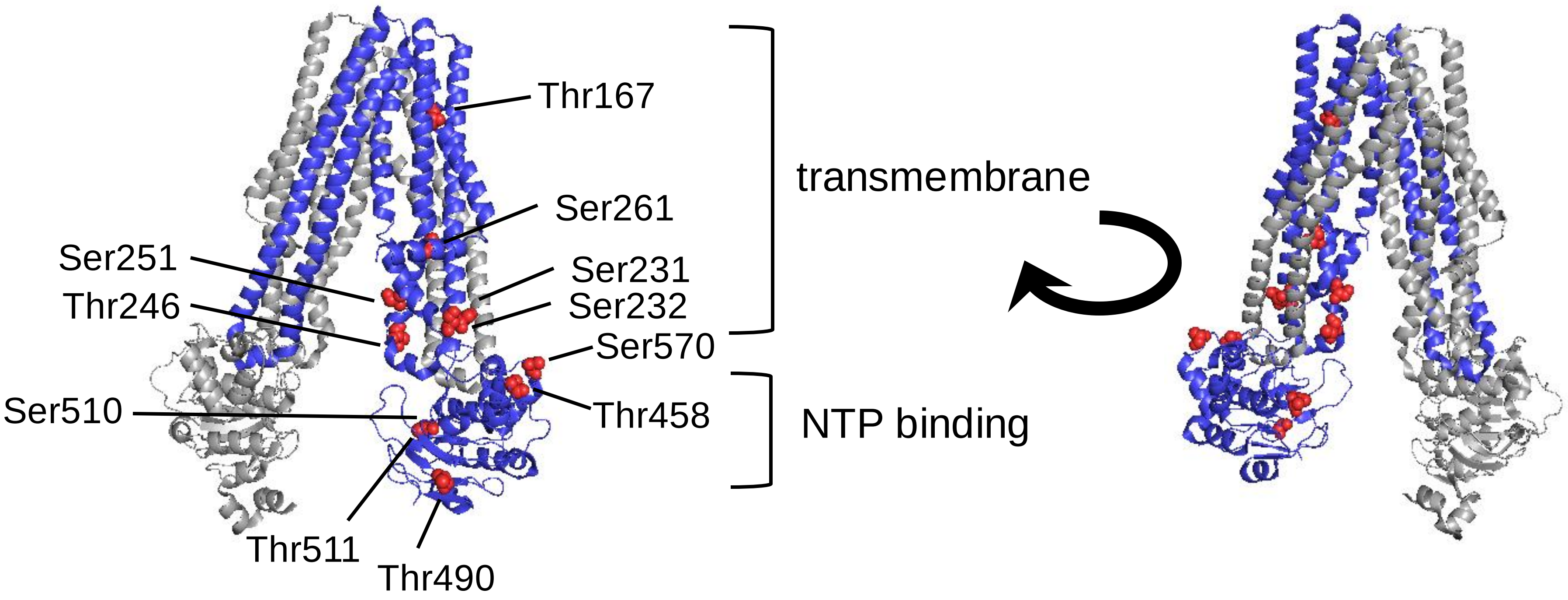


**B** **Known TAP2 phosphorylation sites**

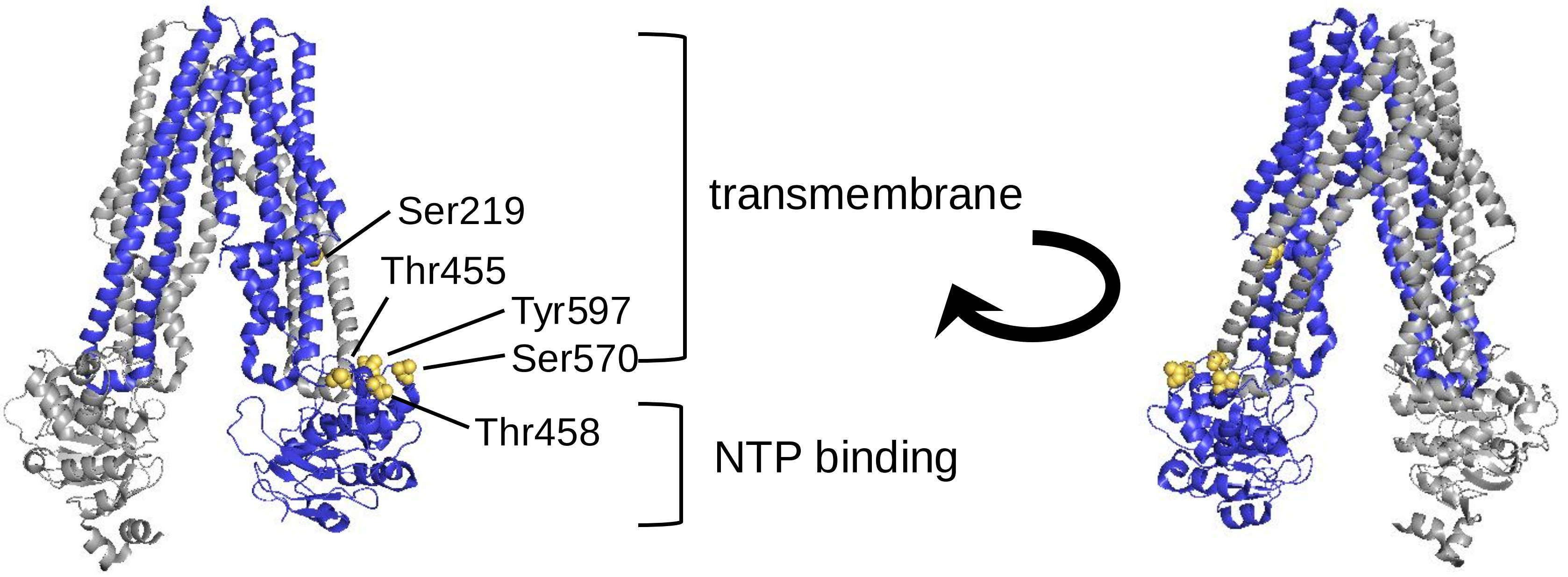
