## Supplemental Tables for "Multiplexed functional analysis of TAP2 variants in regulating MHC-I cell surface abundance reveals overexpression of PLK1 downregulates antigen presentation"

### Table S1 (TAP2 codon selection)

| codon | Site | PTM or Interaction | ClinVar | Conservation * | nucleotide binding region 503-510 | Peptide binding site 301-389, 414-443 | PolyPhen-2 prediction |
|---|---|---|---|---|---|---|---|
| 16 | D16 | salt bridge with TAPBP | | 10 | | | D->A is predicted to be **PROBABLY DAMAGING** with a score of **0.994** ( |
| 34 | G34R | | VUS | 9 | | | G->R is predicted to be **PROBABLY DAMAGING** with a score of **0.997** ( |
| 54 | K54 | Ubiquination | | 7 | | | K->R is predicted to be **BENIGN** with a score of **0.000** (sensitivity: **1.00**; s |
| 78 | S78 | | | 9 | | | S->A is predicted to be **PROBABLY DAMAGING** with a score of **0.982** (s |
| 111 | L111F | | VUS | 9 | | | L->F is predicted to be **PROBABLY DAMAGING** with a score of **1.000** (s |
| 135 | K135 | Ubiquination | | 4 | | | K->R is predicted to be **BENIGN** with a score of **0.000** (sensitivity: **1.00**; s |
| 145 | S145 | | | 9 | | | S->A is predicted to be **POSSIBLY DAMAGING** with a score of **0.896** (se |
| 167 | T167 | | | 10 | | | T->A is predicted to be **POSSIBLY DAMAGING** with a score of **0.771** (se |
| 172 | Y172F | | VUS | 9 | | | Y->F is predicted to be **POSSIBLY DAMAGING** with a score of **0.562** (se |
| 173 | S173 | | | 8 | | | S->A is predicted to be **BENIGN** with a score of **0.060** (sensitivity: **0.94**; s |
| 196 | M196 | peptide interaction | | 10 | | | M->A is predicted to be **PROBABLY DAMAGING** with a score of **0.981** (s |
| 200 | S200 | | | 10 | | | S->A is predicted to be **POSSIBLY DAMAGING** with a score of **0.727** (se |
| 203 | S203 | | | 10 | | | S->A is predicted to be **PROBABLY DAMAGING** with a score of **0.972** (s |
| 204 | S204 | | | 10 | | | S->A is predicted to be **POSSIBLY DAMAGING** with a score of **0.866** (se |
| 219 | S219F | Phosphorylation | VUS | 8 | | | S->F is predicted to be **POSSIBLY DAMAGING** with a score of **0.899** (se |
| 220 | R220Q | | | 9 | | | R->Q is predicted to be **PROBABLY DAMAGING** with a score of **0.999** ( |
| 231 | S231 | | | 9 | | | S->A is predicted to be **PROBABLY DAMAGING** with a score of **1.000** (s |
| 232 | S232 | | | 9 | | | S->A is predicted to be **PROBABLY DAMAGING** with a score of **0.959** (s |
| 244 | T244 | | | 9 | | | T->A is predicted to be **POSSIBLY DAMAGING** with a score of **0.735** (se |
| 245 | K245 | Ubiquination | | 9 | | | K->R is predicted to be **PROBABLY DAMAGING** with a score of **0.999** (s |
| 246 | T246 | | | 9 | | | T->A is predicted to be **POSSIBLY DAMAGING** with a score of **0.953** (se |
| 251 | S251 | | | 10 | | | S->A is predicted to be **PROBABLY DAMAGING** with a score of **0.999** (s |
| 254 | S254 | | | 9 | | | S->A is predicted to be **BENIGN** with a score of **0.149** (sensitivity: **0.92**; s |
| 255 | S255 | | | 9 | | | S->A is predicted to be **POSSIBLY DAMAGING** with a score of **0.777** (se |
| 257 | T257 | | | 9 | | | T->A is predicted to be **PROBABLY DAMAGING** with a score of **0.998** (s |
| 261 | S261 | | | 10 | | | S->A is predicted to be **PROBABLY DAMAGING** with a score of **1.000** (s |
| 273 | R273Q | | VUS | 10 | | | R->Q is predicted to be **PROBABLY DAMAGING** with a score of **1.000** ( |
| 274 | S274 | | | 9 | | | S->A is predicted to be **POSSIBLY DAMAGING** with a score of **0.913** (se |
| 277 | K277 | | | 9 | | | K->R is predicted to be **PROBABLY DAMAGING** with a score of **0.998** (s |
| 289 | S289 | | | 10 | | | S->A is predicted to be **PROBABLY DAMAGING** with a score of **0.999** (s |
| 293 | T293 | | | 9 | | | T->A is predicted to be **BENIGN** with a score of **0.222** (sensitivity: **0.91**; s |
| 295 | L295V | | VUS | 10 | | | L->V is predicted to be **PROBABLY DAMAGING** with a score of **0.994** (s |
| 296 | S296 | | | 9 | | | S->A is predicted to be **POSSIBLY DAMAGING** with a score of **0.613** (se |
| 303 | T303 | | | 8 | | peptide | T->A is predicted to be **BENIGN** with a score of **0.005** (sensitivity: **0.97**; s |
| 308 | K308 | Ubiquination | | 10 | | peptide | K->R is predicted to be **PROBABLY DAMAGING** with a score of **1.000** (s |
| 313 | R313H | | benign | 10 | | peptide | R->H is predicted to be **PROBABLY DAMAGING** with a score of **1.000** (s |
| 341 | T341 | | | 9 | | peptide | T->A is predicted to be **POSSIBLY DAMAGING** with a score of **0.817** (se |
| 343 | R343H | | VUS | 10 | | peptide | R->H is predicted to be **PROBABLY DAMAGING** with a score of **1.000** (s |
| 344 | S344 | | | 9 | | peptide | S->A is predicted to be **BENIGN** with a score of **0.052** (sensitivity: **0.94**; s |
| 355 | Y355 | | | 10 | | peptide | Y->F is predicted to be **PROBABLY DAMAGING** with a score of **0.999** (s |

| | | | | | | | |
|---|---|---|---|---|---|---|---|
| 356 | K356 | Ubiquination | | 9 | | peptide | K->R is predicted to be **BENIGN** with a score of **0.003** (sensitivity: **0.98**; s |
| 368 | R368P | assoc, peptide length selection | VUS | 8 | | peptide | R->P is predicted to be **PROBABLY DAMAGING** with a score of **1.000** (s |
| 369 | R369Q | peptide length selection | | 10 | | peptide | R->Q is predicted to be **POSSIBLY DAMAGING** with a score of **0.947** (se |
| 370 | D370N | cancer assoc | | 10 | | peptide | D->N is predicted to be **POSSIBLY DAMAGING** with a score of **0.494** (se |
| 373 | R373H | cancer assoc | | 6 | | peptide | R->H is predicted to be **BENIGN** with a score of **0.118** (sensitivity: **0.93**; s |
| 376 | Y376 | | | 9 | | peptide | Y->F is predicted to be **BENIGN** with a score of **0.264** (sensitivity: **0.91**; s |
| 380 | R380 | peptide interaction | | 8 | | peptide | R->A is predicted to be **POSSIBLY DAMAGING** with a score of **0.949** (se |
| 405 | T405 | | | 10 | | | T->A is predicted to be **PROBABLY DAMAGING** with a score of **0.996** (s |
| 411 | S411 | | | 9 | | | S->A is predicted to be **POSSIBLY DAMAGING** with a score of **0.718** (se |
| 413 | M413 | peptide interaction | | 3 | | | M->A is predicted to be **PROBABLY DAMAGING** with a score of **0.962** (s |
| 415 | Y415 | | | 10 | | peptide | Y->F is predicted to be **BENIGN** with a score of **0.151** (sensitivity: **0.92**; s |
| 417 | E417 | peptide interaction | | 9 | | peptide | E->A is predicted to be **POSSIBLY DAMAGING** with a score of **0.905** (se |
| 428 | Y428 | | | 10 | | peptide | Y->F is predicted to be **BENIGN** with a score of **0.353** (sensitivity: **0.90**; s |
| 430 | Y430 | | | 8 | | peptide | Y->F is predicted to be **BENIGN** with a score of **0.017** (sensitivity: **0.95**; s |
| 435 | S435 | | | 10 | | peptide | S->A is predicted to be **POSSIBLY DAMAGING** with a score of **0.906** (se |
| 446 | Y446 | | | 10 | | | Y->F is predicted to be **PROBABLY DAMAGING** with a score of **0.999** (s |
| 450 | Q450 | Ubiquitination | | 4 | | | Q->A is predicted to be **BENIGN** with a score of **0.222** (sensitivity: **0.91**; s |
| 455 | S455 | Phosphorylation | | 3 | | | S->A is predicted to be **BENIGN** with a score of **0.000** (sensitivity: **1.00**; s |
| 458 | T458 | Phosphorylation | | 8 | | | T->A is predicted to be **POSSIBLY DAMAGING** with a score of **0.623** (se |
| 469 | K469 | Ubiquination | | 4 | | | K->R is predicted to be **BENIGN** with a score of **0.029** (sensitivity: **0.95**; s |
| 474 | S474 | | | 8 | | | S->A is predicted to be **BENIGN** with a score of **0.002** (sensitivity: **0.99**; s |
| 477 | Y477 | | | 10 | | | Y->F is predicted to be **PROBABLY DAMAGING** with a score of **1.000** (s |
| 490 | T490 | | | 10 | | | T->A is predicted to be **BENIGN** with a score of **0.027** (sensitivity: **0.95**; s |
| 492 | T492 | | | 9 | | | T->A is predicted to be **PROBABLY DAMAGING** with a score of **0.984** (s |
| 499 | T499M | | VUS | 10 | | | T->M is predicted to be **PROBABLY DAMAGING** with a score of **1.000** (s |
| 500 | A500V | | VUS | 9 | | | A->V is predicted to be **PROBABLY DAMAGING** with a score of **0.981** (s |
| 503 | G503 | | | 10 | NTP | | G->A is predicted to be **PROBABLY DAMAGING** with a score of **0.999** (s |
| 504 | P504 | | | 10 | NTP | | P->A is predicted to be **PROBABLY DAMAGING** with a score of **0.992** (s |
| 505 | N505 | | | 10 | NTP | | N->A is predicted to be **PROBABLY DAMAGING** with a score of **0.999** (s |
| 506 | G506 | | | 10 | NTP | | G->A is predicted to be **PROBABLY DAMAGING** with a score of **0.999** (s |
| 507 | S507 | | | 9 | NTP | | S->A is predicted to be **BENIGN** with a score of **0.039** (sensitivity: **0.94**; s |
| 508 | G508 | | | 10 | NTP | | G->A is predicted to be **PROBABLY DAMAGING** with a score of **0.999** (s |
| 509 | K509 | | | 10 | NTP | | K->A is predicted to be **PROBABLY DAMAGING** with a score of **1.000** (s |
| 510 | S510 | | | 10 | NTP | | S->A is predicted to be **PROBABLY DAMAGING** with a score of **0.996** (s |
| 511 | T511 | | | 10 | NTP | | T->A is predicted to be **PROBABLY DAMAGING** with a score of **0.964** (s |
| 520 | Y520 | | | 10 | | | Y->F is predicted to be **POSSIBLY DAMAGING** with a score of **0.922** (se |
| 523 | T523 | | | 9 | | | T->A is predicted to be **BENIGN** with a score of **0.438** (sensitivity: **0.89**; s |
| 533 | P533H | | VUS | 10 | | | P->H is predicted to be **PROBABLY DAMAGING** with a score of **1.000** (s |
| 537 | Y537 | | | 10 | | | Y->F is predicted to be **PROBABLY DAMAGING** with a score of **1.000** (s |
| 541 | Y541 | | | 10 | | | Y->F is predicted to be **PROBABLY DAMAGING** with a score of **1.000** (s |
| 557 | S557 | | | 10 | | | S->A is predicted to be **PROBABLY DAMAGING** with a score of **0.986** (s |
| 559 | S559 | | | 10 | | | S->A is predicted to be **PROBABLY DAMAGING** with a score of **0.992** (s |

| | | | | | |
|---|---|---|---|---|---|
| 566 | Y566 | | | 10 | Y->F is predicted to be **PROBABLY DAMAGING** with a score of **1.000** (s |
| 597 | Y597 | Phosphorylation | | 5 | Y->F is predicted to be **BENIGN** with a score of **0.009** (sensitivity: **0.96**; s |
| 598 | T598 | | | 10 | T->A is predicted to be **POSSIBLY DAMAGING** with a score of **0.876** (se |
| 603 | K603 | Ubiquination | | 8 | K->R is predicted to be **BENIGN** with a score of **0.019** (sensitivity: **0.95**; s |
| 612 | K612 | Ubiquination | | 10 | K->R is predicted to be **PROBABLY DAMAGING** with a score of **0.999** (s |
| 618 | A618S | | VUS | 10 | A->S is predicted to be **PROBABLY DAMAGING** with a score of **0.994** (s |
| 625 | P625L | | | 10 | P->L is predicted to be **PROBABLY DAMAGING** with a score of **1.000** (s |
| 634 | T634 | | | 10 | T->A is predicted to be **PROBABLY DAMAGING** with a score of **0.999** (s |
| 635 | S635 | | | 10 | S->A is predicted to be **PROBABLY DAMAGING** with a score of **0.983** (s |
| 638 | D638A | Innactive peptide transport | | 10 | D->A is predicted to be **PROBABLY DAMAGING** with a score of **1.000** (s |
| 655 | T655 | | | 10 | T->A is predicted to be **BENIGN** with a score of **0.170** (sensitivity: **0.92**; s |
| 665 | T665A | | benign | 9 | T->A is predicted to be **BENIGN** with a score of **0.101** (sensitivity: **0.93**; s |
| 679 | K679 | Ubiquination | | 3 | K->R is predicted to be **BENIGN** with a score of **0.000** (sensitivity: **1.00**; s |
| 682 | K682 | Ubiquination | | 3 | K->R is predicted to be **BENIGN** with a score of **0.022** (sensitivity: **0.95**; s |

*Conservation includes: human, rat, mouse, chicken, chimp, golden hamster, dog, gorilla, wild boar

Table S2 (TAP2 Missense variants)

| index | codon | mut | Score integration 1 | Score integration 2 | Score integration 3 | Score integration 4 | Score average | Surviving Replicates | Variance | read count average |
|---|---|---|---|---|---|---|---|---|---|---|
| p.Asp16Met | 16 | M | 0.38950241 | -0.4666425 | 2.01400068 | -1.5557506 | 0.095277493 | 4 | 2.269905 | 13622.25 |
| p.Asp16Lys | 16 | K | -1.4191042 | 0.49444446 | 1.87490478 | 0.38834645 | 0.334647863 | 4 | 1.825485 | 22823.5 |
| p.Asp16Trp | 16 | W | -0.1422895 | 0.52230461 | 0.93711229 | 0.31073954 | 0.406966726 | 4 | 0.201766 | 6040.5 |
| p.Asp16Val | 16 | V | -0.0926636 | 0.18867618 | 2.95044188 | 0.58558208 | 0.908009135 | 4 | 1.931426 | 24155 |
| p.Asp16Gly | 16 | G | 0.34815047 | 0.10050247 | 2.9891646 | 0.42288317 | 0.965175176 | 4 | 1.839664 | 18662.5 |
| p.Asp16Glu | 16 | E | 0.15804811 | -0.0083517 | 1.6820542 | 2.25326896 | 1.021254883 | 4 | 1.253243 | 20957.25 |
| p.Asp16Arg | 16 | R | -0.217714 | 0.09098396 | 2.64643781 | 1.61062535 | 1.032583269 | 4 | 1.796171 | 25164.25 |
| p.Asp16Ile | 16 | I | -0.3095185 | -0.129509 | 2.65662349 | 1.97212682 | 1.047430691 | 4 | 2.223688 | 16006 |
| p.Asp16Pro | 16 | P | 0.01702847 | 0.19247348 | 2.48618435 | 1.5509717 | 1.061664498 | 4 | 1.371812 | 26448.25 |
| p.Asp16Thr | 16 | T | -0.0681869 | 0.33002393 | 2.55879044 | 1.48057743 | 1.075301221 | 4 | 1.409331 | 34137.75 |
| p.Asp16Ser | 16 | S | -0.0319597 | 0.51788637 | 2.75561928 | 2.00034346 | 1.31047234 | 4 | 1.664896 | 15356.5 |
| p.Asp16Gln | 16 | Q | 0.07211858 | 0.23566478 | 2.60004683 | 2.35067515 | 1.314626334 | 4 | 1.811229 | 22162 |
| p.Asp16Asn | 16 | N | 0.27095123 | -0.3285565 | 2.29411944 | 3.15236829 | 1.347220614 | 4 | 2.707253 | 12211 |
| p.Asp16Ala | 16 | A | 0.5609096 | 0.41153836 | 2.36864459 | 2.27778099 | 1.404718387 | 4 | 1.129937 | 13811.5 |
| p.Asp16Tyr | 16 | Y | 0.33386264 | 0.0191511 | 2.89129374 | 2.46151193 | 1.426454853 | 4 | 2.130453 | 3895.75 |
| p.Asp16His | 16 | H | 0.22902551 | 0.31187074 | 2.85431068 | 2.63682787 | 1.508008701 | 4 | 2.051102 | 5695.5 |
| p.Gly25Asp | 25 | D | -0.6046525 | -0.0505489 | -0.6966079 | -0.0906025 | -0.360602972 | 4 | 0.113831 | 3422.5 |
| p.Leu30Ser | 30 | S | 0.01109771 | 1.99587211 | 2.82781902 | 2.54189378 | 1.844170657 | 4 | 1.612517 | 1169 |
| p.Gly34Met | 34 | M | 0.78384894 | 0.54196928 | 2.30580843 | -0.303278 | 0.832087153 | 4 | 1.182468 | 8150.5 |
| p.Gly34Thr | 34 | T | 0.56968198 | -0.140385 | 3.53630669 | 0.20290692 | 1.042127645 | 4 | 2.848921 | 6520.75 |
| p.Gly34Asp | 34 | D | 0.88348599 | 1.36979441 | 2.10128334 | 1.0577155 | 1.353069808 | 4 | 0.289282 | 3270 |
| p.Gly34Gln | 34 | Q | 0.32235298 | 1.20884589 | 2.18322182 | 2.19526613 | 1.477421706 | 4 | 0.80659 | 5995 |
| p.Gly34His | 34 | H | 0.62857952 | 0.56484714 | 2.91169158 | 2.77783186 | 1.720737524 | 4 | 1.688237 | 6246 |
| p.Gly34Ile | 34 | I | 1.25955769 | -0.0695331 | 2.70043972 | 3.15541084 | 1.761468788 | 4 | 2.143075 | 7398.75 |
| p.Leu44Arg | 44 | R | 0.77957713 | -0.4446712 | 2.44057259 | 2.74272486 | 1.379550856 | 4 | 2.223922 | 2454.25 |
| p.Arg45Gly | 45 | G | 0.22014067 | 0.42191295 | 1.48542683 | 3.2643406 | 1.347955262 | 4 | 1.940317 | 2540.5 |
| p.Lys54His | 54 | H | -0.4560997 | 0.81379664 | 2.19010288 | -0.0593436 | 0.622114049 | 4 | 1.374087 | 7649.5 |
| p.Lys54Ala | 54 | A | -0.6057211 | 1.65975307 | 2.26303441 | -0.7090988 | 0.651991905 | 4 | 2.348483 | 1744 |
| p.Lys54Pro | 54 | P | -0.1008265 | 0.30870238 | 2.5315814 | 1.14345279 | 0.970727525 | 4 | 1.350868 | 7081.5 |
| p.Lys54Leu | 54 | L | 0.06906919 | 0.68520417 | 2.50779464 | 1.69486806 | 1.239234015 | 4 | 1.164361 | 7940.5 |
| p.Lys54Ser | 54 | S | 0.50335239 | 0.94802766 | 0.93236128 | 3.3081223 | 1.422965907 | 4 | 1.621921 | 4969.5 |
| p.Lys54Gly | 54 | G | 0.34599585 | 0.45674675 | 2.4780473 | 3.02174538 | 1.575633819 | 4 | 1.889836 | 7467.75 |

| | | | | | | | | | | |
|---|---|---|---|---|---|---|---|---|---|---|
| p.Lys54Arg | 54 | R | 0.79951502 | 0.54255219 | 2.73696559 | 3.13793489 | 1.804241922 | 4 | 1.750016 | 4169 |
| p.Pro74Thr | 74 | T | 0.72900887 | 1.89851868 | 2.96123043 | 3.73799572 | 2.331688426 | 4 | 1.71008 | 2142.75 |
| p.Thr76Ile | 76 | I | 0.68312598 | 0.97397454 | 2.57469417 | -0.4195389 | 0.953063947 | 4 | 1.529009 | 3307.5 |
| p.Ser78Ala | 78 | A | 0.9489414 | 0.19881045 | 2.72246602 | -0.4408335 | 0.857346094 | 4 | 1.868667 | 2148.5 |
| p.Ser78Trp | 78 | W | -0.9923545 | 0.87126669 | 2.5849625 | 1.11783649 | 0.895427784 | 4 | 2.1561 | 1947.5 |
| p.Ser78Val | 78 | V | 0.7438444 | -0.0940664 | 2.55799545 | 1.02840842 | 1.059045476 | 4 | 1.225603 | 7294.5 |
| p.Ser78Asn | 78 | N | 0.6588536 | 1.3441138 | 1.50355417 | 1.10103278 | 1.151888587 | 4 | 0.13543 | 12107.5 |
| p.Ser78Pro | 78 | P | 0.08248942 | 0.83014024 | 3.38148932 | 0.79944596 | 1.273391236 | 4 | 2.094473 | 10613 |
| p.Ser78Leu | 78 | L | 0.31135423 | 0.50381477 | 2.84308913 | 1.72040102 | 1.344664788 | 4 | 1.387071 | 21571 |
| p.Ser78Arg | 78 | R | 1.06518417 | 0.69075716 | 3.20681241 | 1.0988283 | 1.51539551 | 4 | 1.305713 | 8644 |
| p.Ser78Ile | 78 | I | 0.35920911 | 0.79095427 | 2.54019263 | 2.5672605 | 1.564404128 | 4 | 1.336201 | 6943.5 |
| p.Ser78Thr | 78 | T | 1.36707295 | 1.82127288 | 3.36315076 | 0.22936198 | 1.695214641 | 4 | 1.684768 | 5149 |
| p.Ser78Met | 78 | M | 1.00942941 | 0.81904264 | 2.8268511 | 2.75661012 | 1.852983319 | 4 | 1.181859 | 4057.5 |
| p.Ser78His | 78 | H | 0.01073971 | 1.3498221 | 3.40421601 | 3.00500068 | 1.942444626 | 4 | 2.449495 | 5103.25 |
| p.Ala92Ser | 92 | S | 0.36066049 | 0.51435086 | 2.19730144 | 3.59562098 | 1.666983443 | 4 | 2.345307 | 4249.75 |
| p.Ala108Asp | 108 | D | 0.04059799 | -0.7143473 | 3.06959027 | 0.79646661 | 0.798076895 | 4 | 2.673659 | 2918 |
| p.Leu111Arg | 111 | R | 0.02801813 | 0.15445666 | 2.1500597 | -0.0622484 | 0.56757151 | 4 | 1.120908 | 29252.75 |
| p.Leu111Pro | 111 | P | -0.2777742 | 0.06441905 | 1.29420649 | 1.46905807 | 0.637477349 | 4 | 0.762967 | 22533.25 |
| p.Leu111Asn | 111 | N | 0.80540314 | 0.3594471 | 2.28950662 | 0.07110107 | 0.88136448 | 4 | 0.97252 | 8381.75 |
| p.Leu111Asp | 111 | D | -0.0210452 | 0.33600055 | 1.87555263 | 1.39286986 | 0.895844454 | 4 | 0.786991 | 15133.75 |
| p.Leu111Gln | 111 | Q | 0.81717178 | 0.08103142 | 1.94467044 | 1.02007223 | 0.965736468 | 4 | 0.58868 | 34565 |
| p.Leu111Lys | 111 | K | -0.512577 | 0.28149577 | 2.45731468 | 2.08893942 | 1.078793212 | 4 | 2.02962 | 26427.25 |
| p.Leu111Glu | 111 | E | 0.31765062 | -0.1340304 | 2.75050107 | 1.48726583 | 1.10534677 | 4 | 1.669639 | 9088.25 |
| p.Leu111Ala | 111 | A | 0.8132461 | 1.03415124 | 1.68249304 | 1.18178943 | 1.177919951 | 4 | 0.136088 | 16985.75 |
| p.Leu111Thr | 111 | T | 0.40091488 | 0.4241892 | 3.03924163 | 1.60534032 | 1.367421507 | 4 | 1.55847 | 24216.25 |
| p.Leu111Met | 111 | M | 0.77020001 | -0.6200353 | 3.00646466 | 2.57531233 | 1.432985435 | 4 | 2.811642 | 15359 |
| p.Leu111Ser | 111 | S | 0.41777598 | 0.39627944 | 2.7837681 | 2.40064421 | 1.499616934 | 4 | 1.616209 | 14797.25 |
| p.Leu111Gly | 111 | G | 0.68655512 | 0.24701239 | 2.92109229 | 2.60088272 | 1.613885628 | 4 | 1.803746 | 21989.25 |
| p.Leu111Val | 111 | V | 1.13524848 | 0.49194232 | 3.03603065 | 2.22896627 | 1.723046931 | 4 | 1.280335 | 19641.25 |
| p.Leu111Tyr | 111 | Y | 1.31252259 | 1.63374487 | 2.60665757 | 4.34999201 | 2.475729262 | 4 | 1.863997 | 1583 |
| p.Asp130Asn | 130 | N | 0.85878058 | 1.02245485 | 3.05889369 | 1.21439963 | 1.538632188 | 4 | 1.04832 | 2483.75 |
| p.Lys135Met | 135 | M | 0.18956406 | 0.15082254 | 2.9381653 | -0.0199617 | 0.814647541 | 4 | 2.012431 | 22738.5 |
| p.Lys135His | 135 | H | -0.0978176 | 0.77273071 | 2.49322253 | 0.8099948 | 0.994532613 | 4 | 1.174184 | 16065.5 |
| p.Lys135Gln | 135 | Q | -0.1570853 | 1.39167261 | 1.57210035 | 1.52901737 | 1.083926247 | 4 | 0.690413 | 16918.25 |
| p.Lys135Thr | 135 | T | 0.44396964 | 0.82587059 | 2.52750245 | 0.73226494 | 1.132401903 | 4 | 0.891439 | 31950.25 |

| | | | | | | | | | | |
|---|---|---|---|---|---|---|---|---|---|---|
| p.Lys135Glu | 135 | E | 0.93751647 | 1.23473187 | 1.84855729 | 0.97977497 | 1.250145149 | 4 | 0.17639 | 10990.75 |
| p.Lys135Ser | 135 | S | -0.4809132 | 0.45012134 | 3.29566244 | 2.00998409 | 1.318713661 | 4 | 2.793097 | 11825.5 |
| p.Lys135Leu | 135 | L | 0.45338179 | 0.21016066 | 2.99903272 | 2.01285773 | 1.418858225 | 4 | 1.747627 | 32651 |
| p.Lys135Asp | 135 | D | 0.160593 | 1.05546278 | 3.16619228 | 1.32595396 | 1.427050506 | 4 | 1.592276 | 6249.25 |
| p.Lys135Ala | 135 | A | 0.33581178 | 0.31409489 | 2.4490649 | 2.72996278 | 1.457233589 | 4 | 1.722641 | 21424 |
| p.Lys135Arg | 135 | R | 0.3949346 | 0.27960193 | 3.15650449 | 2.80169131 | 1.65818308 | 4 | 2.34962 | 10680.5 |
| p.Lys135Pro | 135 | P | 0.32038867 | 0.74364979 | 2.78272655 | 3.86564299 | 1.928102002 | 4 | 2.824039 | 24135.25 |
| p.Ser145Cys | 145 | C | -0.4583985 | -0.4074612 | 1.94937393 | 0.58740155 | 0.417728928 | 4 | 1.274421 | 3370.5 |
| p.Ser145Pro | 145 | P | 0.3109778 | 0.79827817 | 1.57853623 | -0.5916078 | 0.524046095 | 4 | 0.825745 | 9351 |
| p.Ser145Val | 145 | V | 0.59747212 | 0.15868986 | 1.71006623 | 0.99294524 | 0.864793363 | 4 | 0.433651 | 18561.75 |
| p.Ser145Leu | 145 | L | 0.20445555 | -0.3267413 | 2.10926222 | 1.67787964 | 0.916214026 | 4 | 1.351679 | 62442.25 |
| p.Ser145Asn | 145 | N | -0.5475735 | 0.76640937 | 2.26614032 | 1.57447013 | 1.01486158 | 4 | 1.460597 | 3601.25 |
| p.Ser145Arg | 145 | R | -0.1318419 | 0.65509436 | 2.52561037 | 1.08655782 | 1.033855155 | 4 | 1.243474 | 15856 |
| p.Ser145Gly | 145 | G | 0.78366482 | -0.5097903 | 2.82765108 | 1.61863021 | 1.180038962 | 4 | 1.973208 | 11844.25 |
| p.Ser145His | 145 | H | 0.56845987 | 0.47730825 | 2.59636726 | 1.10626319 | 1.187099643 | 4 | 0.959696 | 18978.25 |
| p.Ser145Thr | 145 | T | -0.2856396 | 0.34138966 | 1.98200357 | 2.84949893 | 1.221813143 | 4 | 2.091603 | 10664 |
| p.Ser145Gln | 145 | Q | 0.37047067 | 0.52692619 | 2.79147188 | 1.64515952 | 1.333507066 | 4 | 1.266933 | 23935.75 |
| p.Ser145Tyr | 145 | Y | 0.24626382 | -0.3810117 | 2.95756473 | 2.62314569 | 1.361490646 | 4 | 2.806424 | 6105.5 |
| p.Ser145Asp | 145 | D | 0.77782615 | 0.46613549 | 2 | 3.3485223 | 1.648120986 | 4 | 1.723229 | 4554 |
| p.Ser145Phe | 145 | F | 0.36883502 | 0.50650126 | 2.84645474 | 2.92913602 | 1.662731761 | 4 | 2.005339 | 3420.5 |
| p.Ser145Met | 145 | M | 0.83013239 | 1.21002165 | 3.04942971 | 1.79096048 | 1.720136058 | 4 | 0.941454 | 8788.5 |
| p.Ala162Asp | 162 | D | -0.9764453 | -0.0630522 | 2.53138146 | 0.97936023 | 0.617811063 | 4 | 2.265899 | 4125.25 |
| p.Thr167Asp | 167 | D | 0.66935469 | -1.3148733 | 2.38277365 | 0.1440256 | 0.470320151 | 4 | 2.330159 | 3743 |
| p.Thr167Val | 167 | V | -0.6829946 | 0.25933562 | 2.35395695 | 0.87294326 | 0.700810311 | 4 | 1.624113 | 3920.25 |
| p.Thr167Gln | 167 | Q | -0.5533617 | 1.08719844 | 2.22443157 | 3.21244962 | 1.492679485 | 4 | 2.61459 | 8502.75 |
| p.Tyr172Pro | 172 | P | -0.5354753 | -0.0333905 | 1.30800114 | 2.46240033 | 0.800383919 | 4 | 1.833224 | 27175 |
| p.Tyr172Met | 172 | M | 0.24487248 | 0.84757271 | 1.87328707 | 0.80683343 | 0.943141422 | 4 | 0.460155 | 38494.5 |
| p.Tyr172Arg | 172 | R | -0.0933539 | 0.39404373 | 3.03115659 | 1.44522752 | 1.194268477 | 4 | 1.911823 | 26628.75 |
| p.Tyr172Thr | 172 | T | -0.1973733 | 1.21780311 | 2.68477937 | 1.4363726 | 1.285395434 | 4 | 1.394747 | 12868.5 |
| p.Tyr172Lys | 172 | K | 0.24827395 | 0.68198961 | 2.311678 | 1.93815134 | 1.295023225 | 4 | 0.972898 | 12207.75 |
| p.Tyr172His | 172 | H | -0.0834096 | 0.80232397 | 3.04614178 | 1.53051472 | 1.323892713 | 4 | 1.753789 | 9100.75 |
| p.Tyr172Ala | 172 | A | 0.32486761 | 0.70001705 | 2.75861059 | 2.18811395 | 1.492902298 | 4 | 1.359436 | 6223.75 |
| p.Tyr172Phe | 172 | F | 1.02483185 | 1.95078703 | 2.93859946 | 0.56259469 | 1.619203255 | 4 | 1.106818 | 1113.5 |
| p.Tyr172Leu | 172 | L | 0.57871207 | 0.6313966 | 3.2400169 | 2.49169567 | 1.735455311 | 4 | 1.797535 | 20829.25 |
| p.Tyr172Val | 172 | V | 0.99772026 | 0.6741462 | 2.57173342 | 2.99159929 | 1.808799791 | 4 | 1.30879 | 14882.5 |

| | | | | | | | | | | |
|---|---|---|---|---|---|---|---|---|---|---|
| p.Tyr172Cys | 172 | C | 1.15200309 | 1.63226822 | 2.37851162 | 2.46712601 | 1.907477236 | 4 | 0.393854 | 1958.75 |
| p.Tyr172Gly | 172 | G | 0.62746027 | 1.38448391 | 2.07496206 | 4.23095443 | 2.079465169 | 4 | 2.406748 | 5937.5 |
| p.Tyr172Gln | 172 | Q | 1.21631791 | 1.02397886 | 2.86561432 | 3.31550183 | 2.105353229 | 4 | 1.33407 | 6359.25 |
| p.Ser173Phe | 173 | F | -0.5493386 | -1.2658941 | 1.32192809 | -0.4088055 | -0.225527525 | 4 | 1.205142 | 431.75 |
| p.Ser173Gln | 173 | Q | 0.59263031 | 0.54238768 | 2.5849625 | 0.29540802 | 1.003847128 | 4 | 1.127952 | 20612.5 |
| p.Ser173His | 173 | H | 0.80831098 | 0.3071601 | 2.80735492 | 0.65167184 | 1.143624461 | 4 | 1.274041 | 5385.75 |
| p.Ser173Leu | 173 | L | 0.20559043 | 0.50500252 | 2.28337635 | 1.61185328 | 1.151455647 | 4 | 0.935258 | 28805.25 |
| p.Ser173Pro | 173 | P | 0.06520867 | 0.7412502 | 2.4886229 | 1.35813951 | 1.163305319 | 4 | 1.059458 | 57333.5 |
| p.Ser173Glu | 173 | E | 0.34806602 | 0.49196266 | 2.81540259 | 1.26867151 | 1.231025697 | 4 | 1.279166 | 19659 |
| p.Ser173Asp | 173 | D | 0.5277311 | 0.74846732 | 2.75232453 | 1.02519209 | 1.26342876 | 4 | 1.026668 | 27210.5 |
| p.Ser173Ile | 173 | I | -0.0647269 | 0.1714079 | 3.05415577 | 1.99154834 | 1.288096267 | 4 | 2.230312 | 18630.5 |
| p.Ser173Val | 173 | V | 0.47999294 | -0.1260787 | 2.86881234 | 1.9439115 | 1.291659511 | 4 | 1.860543 | 22443.25 |
| p.Ser173Arg | 173 | R | 0.46703452 | 0.08665847 | 2.79750714 | 2.09862803 | 1.362457038 | 4 | 1.67692 | 5811 |
| p.Ser173Thr | 173 | T | 0.03023506 | 0.28137247 | 2.90122185 | 2.68844909 | 1.475319614 | 4 | 2.339553 | 41241.25 |
| p.Ser173Trp | 173 | W | 0.36681234 | 0.145008 | 3.05062607 | 2.39623779 | 1.489671052 | 4 | 2.109125 | 3759.75 |
| p.Ser173Ala | 173 | A | 0.58865784 | 0.41929581 | 1.93733354 | 3.11243254 | 1.514429933 | 4 | 1.596277 | 20064.5 |
| p.Ser173Lys | 173 | K | 0.60513644 | 1.23152628 | 2.70790956 | 1.55541823 | 1.524997626 | 4 | 0.777492 | 15392 |
| p.Ser173Gly | 173 | G | 0.26922262 | 0.5566762 | 2.8987012 | 2.42368159 | 1.537070403 | 4 | 1.736243 | 39664.5 |
| p.Ser173Met | 173 | M | 0.57448895 | 0.35402509 | 3.07610298 | 2.15251892 | 1.539283983 | 4 | 1.691179 | 18046.25 |
| p.Ser173Asn | 173 | N | 0.77634887 | 0.39208917 | 2.92617765 | 3.5240407 | 1.904664095 | 4 | 2.40895 | 14332.5 |
| p.Ala188Ser | 188 | S | 0.46464143 | 1.23331371 | 3.22463089 | 3.30530178 | 2.056971951 | 4 | 2.045228 | 5259.5 |
| p.Met196Asp | 196 | D | 0.28214323 | 0.95751902 | 0.43459977 | -1.675309 | -0.000261748 | 4 | 1.330661 | 5707.75 |
| p.Met196Arg | 196 | R | -0.3065178 | -0.4740135 | 1.5740536 | 0.14886339 | 0.235596412 | 4 | 0.865475 | 8924.5 |
| p.Met196Glu | 196 | E | 0.90633239 | 0.65042289 | 1.72964225 | -0.7269041 | 0.639873348 | 4 | 1.042263 | 8123 |
| p.Met196Gln | 196 | Q | 0.83380986 | -0.1641592 | 1.80735492 | 0.28669343 | 0.690924766 | 4 | 0.720468 | 9729.25 |
| p.Met196Cys | 196 | C | -0.005416 | 0.40075596 | 3.24302038 | 0.15391268 | 0.948068247 | 4 | 2.368724 | 7954.5 |
| p.Met196Asn | 196 | N | 0.46826628 | 0.40352351 | 1.94144782 | 1.19864382 | 1.002970356 | 4 | 0.521424 | 8841.5 |
| p.Met196His | 196 | H | 0.58985224 | -0.2596851 | 2.61223543 | 1.14689362 | 1.022324038 | 4 | 1.457972 | 10645.75 |
| p.Met196Gly | 196 | G | 0.26771804 | 0.0808736 | 2.66389638 | 2.42075592 | 1.358310985 | 4 | 1.884861 | 11700.25 |
| p.Met196Pro | 196 | P | 0.45950982 | 0.54205212 | 2.19950599 | 2.59843462 | 1.449875638 | 4 | 1.2287 | 13845 |
| p.Met196Leu | 196 | L | 0.15035855 | 0.32076581 | 2.71044161 | 2.6387474 | 1.45507834 | 4 | 1.988656 | 38014.75 |
| p.Met196Ser | 196 | S | 0.15040199 | 0.40298875 | 3.10507386 | 2.63567869 | 1.573535821 | 4 | 2.289749 | 9225.25 |
| p.Met196Lys | 196 | K | 1.37513719 | 0.12376756 | 2.55654682 | 2.48254693 | 1.634499624 | 4 | 1.306312 | 4905.25 |
| p.Ser200Glu | 200 | E | 0.32553935 | 0.19634056 | 2.51592086 | -0.9717417 | 0.516514756 | 4 | 2.117172 | 4538.75 |
| p.Ser200Pro | 200 | P | -0.4761518 | 0.23244807 | 2.44522259 | 1.27176077 | 0.868319901 | 4 | 1.620441 | 18610.25 |

| | | | | | | | | | | |
|---|---|---|---|---|---|---|---|---|---|---|
| p.Ser200Tyr | 200 | Y | 0.65795224 | 0.87372118 | 3.11769504 | -0.6672109 | 0.995539389 | 4 | 2.465696 | 4348.25 |
| p.Ser200Trp | 200 | W | 0.93720487 | 1.93555129 | 1.03606925 | 0.52442096 | 1.108311595 | 4 | 0.35325 | 1846 |
| p.Ser200His | 200 | H | 1.12588444 | 0.10299712 | 2.52685955 | 0.72676347 | 1.120626142 | 4 | 1.056072 | 9641.75 |
| p.Ser200Gln | 200 | Q | 0.16033145 | 0.41434506 | 2.44914865 | 1.61911502 | 1.160735046 | 4 | 1.142676 | 8368.75 |
| p.Ser200Phe | 200 | F | 0.1768414 | 1.06863844 | 3.13577678 | 0.59825932 | 1.244878985 | 4 | 1.721792 | 5752.75 |
| p.Ser200Gly | 200 | G | 1.13475422 | 1.25143858 | 1.94137227 | 0.95850787 | 1.321518235 | 4 | 0.185263 | 4466 |
| p.Ser200Leu | 200 | L | 0.56634682 | 0.95870383 | 2.68189024 | 1.45710156 | 1.416010615 | 4 | 0.845066 | 53664.5 |
| p.Ser200Thr | 200 | T | 0.18715142 | 0.67126479 | 2.69750774 | 2.56887654 | 1.531200122 | 4 | 1.661 | 21236 |
| p.Ser200Arg | 200 | R | -0.0042908 | 0.38660426 | 2.99879725 | 2.8806716 | 1.565445589 | 4 | 2.546019 | 18421.25 |
| p.Ser200Lys | 200 | K | 0.12775555 | 1.52930996 | 2.75344264 | 2.0689918 | 1.619874987 | 4 | 1.240435 | 5814.5 |
| p.Ser200Val | 200 | V | 0.1155648 | 1.15276525 | 2.68392895 | 2.58623281 | 1.634622955 | 4 | 1.515443 | 15683.5 |
| p.Ser200Asp | 200 | D | 0.63095727 | 1.35609033 | 2.5417093 | 2.99939472 | 1.882037904 | 4 | 1.175159 | 5735.25 |
| p.Ser203Val | 203 | V | 0.70140441 | 0.48101154 | 1.90046433 | -0.0882006 | 0.748669917 | 4 | 0.700286 | 3503.5 |
| p.Ser203Met | 203 | M | -0.1381211 | 0.48655261 | 2.85721544 | 0.10477888 | 0.827606466 | 4 | 1.896913 | 4897.25 |
| p.Ser203Thr | 203 | T | 1.31665957 | 0.96716139 | 2.46760555 | -0.7155466 | 1.008969975 | 4 | 1.732665 | 5769.5 |
| p.Ser203Ile | 203 | I | -0.0205038 | 0.68079201 | 2.04975304 | 1.44546309 | 1.038876092 | 4 | 0.812565 | 4657.25 |
| p.Ser203Leu | 203 | L | 0.40821992 | 0.81114592 | 2.61197545 | 0.77791385 | 1.152313784 | 4 | 0.980286 | 8817 |
| p.Ser203His | 203 | H | 0.39940685 | 0.55183026 | 2.13993026 | 3.19430916 | 1.571369132 | 4 | 1.79005 | 3644 |
| p.Ser203Gly | 203 | G | -0.2709978 | 1.1742744 | 2.36632221 | 3.36580779 | 1.658851656 | 4 | 2.457783 | 3702.25 |
| p.Ser203Phe | 203 | F | 1.81341243 | -0.1484396 | 2.81526012 | 2.64310654 | 1.780834868 | 4 | 1.84557 | 3869 |
| p.Ser203Cys | 203 | C | 1.52533852 | 0.85033515 | 2.80735492 | 4.26303441 | 2.361515749 | 4 | 2.265802 | 2111 |
| p.Ser203Ala | 203 | A | 1.75450304 | 1.76873363 | 1.84799691 | 4.4493074 | 2.455135244 | 4 | 1.769124 | 1347.75 |
| p.Ser204Val | 204 | V | 0.25772231 | -0.1274168 | 1.26848884 | 0.81012987 | 0.552231046 | 4 | 0.376065 | 3841.5 |
| p.Ser204Trp | 204 | W | 0.11532414 | 0.23650536 | 2.16803789 | 1.74586412 | 1.066432878 | 4 | 1.089516 | 8953.5 |
| p.Ser204Tyr | 204 | Y | 0.30374985 | -0.5181211 | 2.7398481 | 1.80560726 | 1.082771024 | 4 | 2.146042 | 5121 |
| p.Ser204Met | 204 | M | 0.02345897 | 0.72217587 | 2.76167211 | 0.98158796 | 1.122223727 | 4 | 1.358297 | 13750.25 |
| p.Ser204Ala | 204 | A | 0.0694214 | 0.12780374 | 2.83564397 | 1.80783751 | 1.210176654 | 4 | 1.824065 | 9667.75 |
| p.Ser204Gln | 204 | Q | -0.076589 | 0.23664058 | 3.36380474 | 1.33083021 | 1.213671635 | 4 | 2.41872 | 14765.5 |
| p.Ser204Phe | 204 | F | 0.07860549 | 0.41862029 | 2.75372544 | 1.82746446 | 1.269603922 | 4 | 1.552158 | 19017 |
| p.Ser204Pro | 204 | P | 0.62913053 | 0.45888445 | 2.44036166 | 1.75845428 | 1.321707728 | 4 | 0.888754 | 24481 |
| p.Ser204Leu | 204 | L | 0.5777323 | 0.41595466 | 2.77810598 | 1.71329219 | 1.371271283 | 4 | 1.212832 | 70750.25 |
| p.Ser204Ile | 204 | I | 0.48379501 | 0.69700039 | 2.8696392 | 1.51494932 | 1.391345979 | 4 | 1.168798 | 20688.75 |
| p.Ser204His | 204 | H | 0.97442137 | 0.00196526 | 1.87231036 | 3.04123232 | 1.472482329 | 4 | 1.677108 | 18374.75 |
| p.Ser204Thr | 204 | T | 0.40553403 | 1.2554845 | 2.41169406 | 1.88412198 | 1.48920864 | 4 | 0.745305 | 17497.5 |
| p.Ser204Asn | 204 | N | 0.5723065 | 0.77418186 | 2.8964236 | 1.85096815 | 1.523470026 | 4 | 1.1528 | 13748.5 |

| | | | | | | | | | | | |
|---|---|---|---|---|---|---|---|---|---|---|---|
| p.Ser204Glu | 204 | E | 0.20281688 | 0.12469575 | 3.27753398 | 2.80735492 | 1.603100382 | | 4 | 2.800144 | 1299.75 |
| p.Ser204Cys | 204 | C | 1.61768681 | -0.0303 | 2.23681949 | 2.87114961 | 1.673838967 | | 4 | 1.552581 | 13479.25 |
| p.Ala207Thr | 207 | T | 0.31232381 | 1.90846645 | 3.60203601 | 2.19162007 | 2.003611587 | | 4 | 1.819938 | 931.75 |
| p.Ser219Phe | 219 | F | -0.1418481 | -0.8622775 | -0.4594316 | 1.7331889 | 0.067407922 | | 4 | 1.320163 | 4437.25 |
| p.Ser219Gln | 219 | Q | 0.455051 | -0.25529 | 1.34680276 | -0.1813599 | 0.341300962 | | 4 | 0.551023 | 8632.75 |
| p.Ser219Pro | 219 | P | 0.40171612 | 0.38541884 | 2.15931222 | 0.88633028 | 0.958194365 | | 4 | 0.695196 | 30740.25 |
| p.Ser219Lys | 219 | K | 0.09478436 | 0.8681564 | 1.64356323 | 1.31885823 | 0.981340556 | | 4 | 0.450417 | 16869.5 |
| p.Ser219Thr | 219 | T | 0.69821104 | 0.58685904 | 2.29727632 | 0.43424097 | 1.004146842 | | 4 | 0.754901 | 25830.25 |
| p.Ser219Asn | 219 | N | -0.6189098 | 0.27589362 | 2.72792045 | 2.73039294 | 1.278824295 | | 4 | 2.938065 | 726 |
| p.Ser219Ala | 219 | A | 0.00284649 | 0.75170325 | 2.83634359 | 1.64923483 | 1.310032037 | | 4 | 1.488384 | 15718.5 |
| p.Ser219Gly | 219 | G | 0.64525619 | 0.62047247 | 2.23054787 | 1.8508863 | 1.336790705 | | 4 | 0.684809 | 27434.25 |
| p.Ser219Val | 219 | V | 0.03447846 | 0.0123934 | 2.86760146 | 2.5185751 | 1.358262107 | | 4 | 2.396066 | 24256.75 |
| p.Ser219Glu | 219 | E | 0.55242968 | -0.0100647 | 3.19307303 | 1.88050641 | 1.403986096 | | 4 | 2.050864 | 11106 |
| p.Ser219Cys | 219 | C | 0.34745553 | 0.82497584 | 1.14622074 | 4.01978465 | 1.584609191 | | 4 | 2.743285 | 5639.5 |
| p.Ser219Tyr | 219 | Y | 0.658893 | 1.26216084 | 2.95664363 | 1.74050161 | 1.654549772 | | 4 | 0.949379 | 12695 |
| p.Ser219Leu | 219 | L | 0.46885055 | 0.75331359 | 2.4710258 | 3.06567504 | 1.689716244 | | 4 | 1.62369 | 14481 |
| p.Ser219Arg | 219 | R | 0.39159026 | 1.53486627 | 2.67590044 | 3.11822155 | 1.930144631 | | 4 | 1.497024 | 14486.75 |
| p.Ser219Ile | 219 | I | 0.50327132 | 0.70447001 | 2.61610101 | 3.94222994 | 1.941518072 | | 4 | 2.685584 | 21124.5 |
| p.Arg220Ser | 220 | S | -0.0463842 | 0.52028965 | 1.77949021 | -0.2460074 | 0.501847069 | | 4 | 0.830852 | 11269.75 |
| p.Arg220His | 220 | H | -0.7718406 | 1.18654722 | 1.32192809 | 0.63321767 | 0.592463088 | | 4 | 0.916014 | 2380.75 |
| p.Arg220Gly | 220 | G | 1.22953449 | -0.1831767 | 2.02090899 | -0.2311626 | 0.70902605 | | 4 | 1.223982 | 5041.25 |
| p.Arg220Pro | 220 | P | -0.3841136 | 0.36364391 | 3.02458564 | 1.25157964 | 1.063923886 | | 4 | 2.155538 | 10051.5 |
| p.Arg220Tyr | 220 | Y | -0.2363953 | 0.44800899 | 1.90000421 | 2.97662549 | 1.272060854 | | 4 | 2.084785 | 4069.75 |
| p.Arg220Leu | 220 | L | 0.36978994 | 0.69049031 | 2.68692491 | 1.53412654 | 1.320332925 | | 4 | 1.071172 | 11483.75 |
| p.Arg220Gln | 220 | Q | 0.67201292 | 0.27226953 | 1.31807577 | 3.61549687 | 1.469463773 | | 4 | 2.232526 | 7828.5 |
| p.Arg220Glu | 220 | E | 0.6522444 | 0.99669233 | 2.66073001 | 3.48542683 | 1.948773392 | | 4 | 1.818544 | 3886 |
| p.Ser231Pro | 231 | P | 0.75985017 | 0.14851264 | 1.34456894 | 0.89415365 | 0.786771349 | | 4 | 0.243589 | 19158 |
| p.Ser231Glu | 231 | E | 0.9338937 | -0.0759239 | 2.29852349 | 0.54022211 | 0.924178858 | | 4 | 1.012182 | 10998.25 |
| p.Ser231Phe | 231 | F | -0.8285195 | 1.50919499 | 2.14684139 | 1.15340991 | 0.995231691 | | 4 | 1.647151 | 3011.75 |
| p.Ser231Tyr | 231 | Y | 0.73198219 | 1.28345395 | 2.35049725 | -0.1634987 | 1.050608662 | | 4 | 1.106502 | 1533 |
| p.Ser231Arg | 231 | R | -0.0606471 | 0.41017301 | 2.70593222 | 1.17326071 | 1.057179699 | | 4 | 1.466671 | 17206.75 |
| p.Ser231Ile | 231 | I | 0.00970746 | -0.6761601 | 3.06441201 | 1.92060659 | 1.079641488 | | 4 | 2.958045 | 10353 |
| p.Ser231Lys | 231 | K | 0.19467166 | -0.115966 | 3.50932806 | 1.59991284 | 1.296986638 | | 4 | 2.732584 | 3691.25 |
| p.Ser231Val | 231 | V | 0.13816402 | 0.33771676 | 2.84785919 | 1.89351157 | 1.304312882 | | 4 | 1.674634 | 14795.5 |
| p.Ser231Thr | 231 | T | 0.33935054 | 0.54386823 | 3.01917011 | 1.38056709 | 1.320738994 | | 4 | 1.484966 | 18447 |

| | | | | | | | | | | |
|---|---|---|---|---|---|---|---|---|---|---|
| p.Ser231Ala | 231 | A | 0.30976917 | 0.41161473 | 2.80735492 | 1.97347371 | 1.375553132 | 4 | 1.490879 | 6629.75 |
| p.Ser231His | 231 | H | 0.42808433 | 1.00544906 | 2.081665 | 2.90546652 | 1.605166225 | 4 | 1.221005 | 16585.5 |
| p.Ser231Met | 231 | M | 0.25599748 | 0.74648026 | 2.90000421 | 2.52174682 | 1.606057194 | 4 | 1.69144 | 12377.5 |
| p.Ser231Asn | 231 | N | 1.38541918 | 1.17163537 | 2.89907109 | 3.38593381 | 2.210514862 | 4 | 1.205258 | 9127.75 |
| p.Ser231Gly | 231 | G | 1.02480049 | 1.09231766 | 2.99462282 | 4.06526866 | 2.294252408 | 4 | 2.227724 | 18266.25 |
| p.Ser232Gln | 232 | Q | -0.3089582 | 0.43849952 | 1.55527958 | 1.41378112 | 0.774650514 | 4 | 0.768358 | 9844 |
| p.Ser232Pro | 232 | P | 0.20616115 | 0.23884963 | 2.88041838 | -0.0424353 | 0.820748474 | 4 | 1.901217 | 7800 |
| p.Ser232Glu | 232 | E | 0.17430344 | 0.28432374 | 2.12730432 | 2.38142911 | 1.241840153 | 4 | 1.379727 | 13112.25 |
| p.Ser232Ile | 232 | I | 0.70384271 | 0.67837753 | 2.22723368 | 1.94585375 | 1.388826918 | 4 | 0.662382 | 13260.5 |
| p.Ser232Thr | 232 | T | 0.47708753 | 0.06983832 | 3.06735527 | 1.95074786 | 1.391257243 | 4 | 1.901396 | 10398.25 |
| p.Ser232Arg | 232 | R | -0.6206236 | 0.79279487 | 3.14660148 | 2.29532966 | 1.403525596 | 4 | 2.767933 | 11417 |
| p.Ser232Met | 232 | M | 0.51771684 | 0.65716701 | 2.2451125 | 2.50250034 | 1.480624172 | 4 | 1.077982 | 5922.25 |
| p.Ser232Lys | 232 | K | 0.73696559 | 1.05763863 | 2.54056838 | 1.59344897 | 1.482155392 | 4 | 0.622716 | 3326 |
| p.Ser232Gly | 232 | G | 0.99716841 | 0.98122612 | 3.81090399 | 0.15153308 | 1.4852079 | 4 | 2.55991 | 8176 |
| p.Ser232Leu | 232 | L | 0.9028643 | 0.40930905 | 2.89553073 | 3.00506452 | 1.80319215 | 4 | 1.797067 | 16375.5 |
| p.Ser232Ala | 232 | A | 0.96410334 | 0.88427564 | 3.34314458 | 4.20945337 | 2.350244232 | 4 | 2.837653 | 3530 |
| p.Phe241Leu | 241 | L | 0.10445116 | 1.31719018 | 0.31194401 | 2.10780329 | 0.960347159 | 4 | 0.865659 | 986.5 |
| p.Thr244Val | 244 | V | 0.37693967 | 0.21011844 | 2.1301796 | 1.0689729 | 0.946552652 | 4 | 0.760918 | 64850 |
| p.Thr244Ala | 244 | A | 0.24117793 | 0.38398758 | 2.18831544 | 1.91742773 | 1.182727169 | 4 | 1.025164 | 48290.25 |
| p.Thr244Gln | 244 | Q | 0.26737399 | 0.66731863 | 2.48421267 | 1.51721401 | 1.234029826 | 4 | 0.966245 | 41813.5 |
| p.Thr244Asn | 244 | N | 0.79436214 | 0.61562773 | 3.02834463 | 1.03791623 | 1.369062681 | 4 | 1.253606 | 25213.25 |
| p.Thr244Tyr | 244 | Y | 0.17861403 | 0.71780766 | 2.4069642 | 2.18324484 | 1.371657681 | 4 | 1.193802 | 18698 |
| p.Thr244Cys | 244 | C | 1.20576638 | 0.5443559 | 1.57095564 | 2.17416511 | 1.373810758 | 4 | 0.465222 | 4936.75 |
| p.Thr244Ser | 244 | S | 0.46012228 | 0.72565271 | 2.45995198 | 1.91459307 | 1.390080009 | 4 | 0.908675 | 38417.25 |
| p.Thr244Pro | 244 | P | 0.47361993 | 0.58427801 | 2.75776064 | 1.86784991 | 1.420877122 | 4 | 1.194745 | 61980.75 |
| p.Thr244Met | 244 | M | 0.95004184 | 1.33320717 | 2.92545305 | 0.56786007 | 1.444140533 | 4 | 1.072865 | 9470 |
| p.Thr244Lys | 244 | K | 0.56906779 | 0.57771048 | 2.54217001 | 2.4176694 | 1.526654422 | 4 | 1.214215 | 26281.75 |
| p.Thr244Glu | 244 | E | 0.74118733 | -0.0622984 | 3.10657166 | 2.34089418 | 1.531588694 | 4 | 2.100252 | 17768.75 |
| p.Thr244Asp | 244 | D | 0.49917978 | 0.75758428 | 2.60696641 | 2.39020809 | 1.56348464 | 4 | 1.184849 | 38907.5 |
| p.Thr244Arg | 244 | R | 0.32056841 | 0.63886359 | 2.96936171 | 2.32859183 | 1.564346384 | 4 | 1.65388 | 41672.25 |
| p.Thr244His | 244 | H | 0.4157455 | 0.16741644 | 3.3892744 | 2.30892951 | 1.570341463 | 4 | 2.385107 | 15443.5 |
| p.Thr244Leu | 244 | L | 0.63758161 | 0.2494074 | 3.16374643 | 2.44935124 | 1.62502167 | 4 | 1.971515 | 35313.75 |
| p.Thr244Gly | 244 | G | 0.20249741 | 0.82030291 | 2.54018321 | 3.5051851 | 1.76704216 | 4 | 2.321001 | 75922.5 |
| p.Thr244Ile | 244 | I | 0.82347666 | 0.41405897 | 2.84741879 | 3.4313308 | 1.879071305 | 4 | 2.202583 | 23283.75 |
| p.Lys245Glu | 245 | E | 0.54929563 | 1.07352904 | 2.31525409 | -0.8307848 | 0.776823485 | 4 | 1.696992 | 9225.75 |

| | | | | | | | | | | |
|---|---|---|---|---|---|---|---|---|---|---|
| p.Lys245His | 245 | H | -0.0297473 | 0.23105369 | 3 | 0.11759623 | 0.829725646 | 4 | 2.104774 | 5215.5 |
| p.Lys245Leu | 245 | L | -0.0350916 | 0.73309759 | 2.33647097 | 1.69530661 | 1.182445892 | 4 | 1.093037 | 33042 |
| p.Lys245Arg | 245 | R | 0.1084932 | 0.48821813 | 3.00070153 | 1.29866935 | 1.224020554 | 4 | 1.649325 | 18623 |
| p.Lys245Tyr | 245 | Y | 0.4732576 | 0.80338602 | 2.24444934 | 1.61284153 | 1.283483624 | 4 | 0.639631 | 8545 |
| p.Lys245Ser | 245 | S | 0.9827294 | 0.55869399 | 2.49123245 | 1.31456535 | 1.336805298 | 4 | 0.688008 | 36143.25 |
| p.Lys245Pro | 245 | P | 0.42648129 | 0.59330348 | 2.64346884 | 1.9152299 | 1.394620876 | 4 | 1.136686 | 20681.75 |
| p.Lys245Phe | 245 | F | -0.6834719 | 0.89247154 | 2.36427439 | 3.04836302 | 1.405409262 | 4 | 2.748416 | 2684 |
| p.Lys245Ile | 245 | I | 0.51009458 | 1.18637944 | 3.14505033 | 0.96737858 | 1.452225733 | 4 | 1.353006 | 3296 |
| p.Lys245Val | 245 | V | -0.0453576 | 1.0619056 | 2.70043972 | 2.13611253 | 1.463275057 | 4 | 1.473452 | 13286 |
| p.Lys245Gln | 245 | Q | 0.05476233 | 0.80970507 | 2.38172475 | 2.90595195 | 1.538036025 | 4 | 1.771191 | 14378.25 |
| p.Lys245Thr | 245 | T | 0.53286442 | 0.73467238 | 3.08109332 | 2.51865229 | 1.716820604 | 4 | 1.623514 | 22304 |
| p.Lys245Gly | 245 | G | 1.03663801 | 0.60338617 | 2.39490754 | 3.0088059 | 1.760934403 | 4 | 1.274543 | 17258.25 |
| p.Lys245Met | 245 | M | 1.71043496 | -0.1288517 | 3.42017219 | 2.39359987 | 1.848838842 | 4 | 2.232089 | 12219.25 |
| p.Thr246Glu | 246 | E | 0.43499776 | 0.33837273 | 2.05219621 | -0.1601405 | 0.666356552 | 4 | 0.921583 | 17075.75 |
| p.Thr246Asp | 246 | D | 0.57368824 | 0.61323199 | 2.38522403 | 0.55374377 | 1.031472006 | 4 | 0.81512 | 45532.75 |
| p.Thr246Met | 246 | M | -0.2456094 | 0.46789072 | 2.14043683 | 1.95784457 | 1.080140687 | 4 | 1.342352 | 28745.5 |
| p.Thr246Phe | 246 | F | 1.10119299 | 0.08413534 | 2.51295126 | 0.71234181 | 1.102655347 | 4 | 1.059555 | 2929.75 |
| p.Thr246Leu | 246 | L | 0.28040521 | 0.17028027 | 2.38365086 | 1.66939449 | 1.125932707 | 4 | 1.168465 | 54545.5 |
| p.Thr246Ala | 246 | A | 0.43217444 | 0.19442263 | 2.80735492 | 1.21167827 | 1.161407565 | 4 | 1.392837 | 46989.5 |
| p.Thr246Lys | 246 | K | 0.38144664 | 0.08441134 | 2.49472215 | 2.00848392 | 1.242266013 | 4 | 1.412458 | 42215.75 |
| p.Thr246Pro | 246 | P | 0.30495235 | 0.14089595 | 2.50294113 | 2.08976685 | 1.259639071 | 4 | 1.465975 | 85415.5 |
| p.Thr246Asn | 246 | N | 0.03847028 | 0.57779784 | 2.44051004 | 2.02792734 | 1.271176374 | 4 | 1.31345 | 39137 |
| p.Thr246Ile | 246 | I | -0.5605856 | 0.55913514 | 2.68589141 | 2.4062959 | 1.27268422 | 4 | 2.384087 | 10261.75 |
| p.Thr246Gly | 246 | G | 0.01789891 | 0.26680364 | 3.03813513 | 2.19541075 | 1.379562107 | 4 | 2.169611 | 22845.25 |
| p.Thr246Cys | 246 | C | -0.084213 | 0.98641422 | 2.18807235 | 2.49569516 | 1.396492181 | 4 | 1.398499 | 5653.75 |
| p.Thr246Ser | 246 | S | 0.50812296 | 0.17408709 | 2.32904883 | 2.6233489 | 1.408651944 | 4 | 1.552574 | 38319.75 |
| p.Thr246Gln | 246 | Q | 0.52493941 | 0.39032378 | 2.80766158 | 1.94074365 | 1.415917105 | 4 | 1.352693 | 54074.75 |
| p.Thr246Val | 246 | V | 0.38514444 | 0.75227122 | 2.97003557 | 1.7455107 | 1.463240483 | 4 | 1.339292 | 34563.25 |
| p.Thr246His | 246 | H | -0.2945769 | 0.87842343 | 2.60304064 | 2.78822892 | 1.493779029 | 4 | 2.16098 | 22653 |
| p.Thr246Trp | 246 | W | 1.02959089 | -0.3457614 | 2.75383941 | 2.58096279 | 1.504657914 | 4 | 2.122876 | 5915 |
| p.Thr246Arg | 246 | R | 0.01241036 | 0.63486385 | 2.53630456 | 3.07808174 | 1.565415124 | 4 | 2.169512 | 41914.25 |
| p.Thr246Tyr | 246 | Y | 0.55891271 | 1.04878856 | 3.04092026 | 1.92785021 | 1.644117936 | 4 | 1.187883 | 11708.75 |
| p.Ser251Glu | 251 | E | 0.98727493 | 0.00421225 | 1.57423609 | -1.144705 | 0.355254567 | 4 | 1.419492 | 5348.5 |
| p.Ser251Ile | 251 | I | 0.39007388 | 0.5623483 | 1.79576695 | -0.0595769 | 0.672153046 | 4 | 0.629854 | 12948 |
| p.Ser251Thr | 251 | T | -0.3410091 | 0.54400855 | 2.33799646 | 0.98129236 | 0.880572057 | 4 | 1.246589 | 10626.5 |

| | | | | | | | | | | | |
|---|---|---|---|---|---|---|---|---|---|---|---|
| p.Ser251Pro | 251 | P | 0.0038889 | 0.68080274 | 2.52867619 | 1.14417418 | 1.0893855 | | 4 | 1.139934 | 30959.5 |
| p.Ser251Asn | 251 | N | 0.33721691 | 0.21458798 | 2.51846709 | 1.29837902 | 1.092162752 | | 4 | 1.138983 | 6938 |
| p.Ser251Gly | 251 | G | 0.71989208 | 0.0396957 | 1.85414913 | 2.40237793 | 1.254028711 | | 4 | 1.146252 | 7618.5 |
| p.Ser251Trp | 251 | W | 2.56580755 | 0.02303097 | 2.72869798 | -0.2937312 | 1.255951323 | | 4 | 2.602105 | 2072 |
| p.Ser251Gln | 251 | Q | -0.1751404 | 1.20819502 | 2.14684139 | 2.0233325 | 1.300807116 | | 4 | 1.141605 | 2666.5 |
| p.Ser251Leu | 251 | L | 0.16082569 | 0.41148669 | 3.18067424 | 1.54954329 | 1.325632477 | | 4 | 1.894584 | 10998 |
| p.Ser251His | 251 | H | 0.65533986 | 0.06301721 | 2.72109919 | 1.95170026 | 1.347789131 | | 4 | 1.460271 | 23176 |
| p.Ser251Arg | 251 | R | 0.2663475 | -0.0370461 | 2.50106124 | 3.29347552 | 1.505959542 | | 4 | 2.700982 | 16778 |
| p.Ser251Tyr | 251 | Y | 1.02961877 | -0.0070778 | 2.63420602 | 2.66813288 | 1.581219959 | | 4 | 1.705704 | 2423 |
| p.Ser251Met | 251 | M | 0.94509291 | 1.10351668 | 3.18789155 | 1.33288846 | 1.6423474 | | 4 | 1.086991 | 11720.75 |
| p.Ser254Thr | 254 | T | -0.971711 | 1.18057225 | 0.79731525 | -0.5168398 | 0.12233419 | | 4 | 1.060315 | 3795.25 |
| p.Ser254Leu | 254 | L | -0.1881052 | 0.96868982 | 2.15625006 | 1.41395965 | 1.087698589 | | 4 | 0.963362 | 16441.25 |
| p.Ser254Asn | 254 | N | 1.25876818 | 1.65287782 | 2.47909387 | -0.5779761 | 1.203190949 | | 4 | 1.668597 | 8108.75 |
| p.Ser254His | 254 | H | 0.32060795 | 1.23533728 | 2.33532092 | 1.73108218 | 1.405587085 | | 4 | 0.725506 | 10051.25 |
| p.Ser254Gly | 254 | G | 0.21210716 | 0.46090865 | 2.5514878 | 2.47558233 | 1.425021485 | | 4 | 1.591093 | 13578.25 |
| p.Ser254Ala | 254 | A | 0.60270078 | 0.80365773 | 1.67024769 | 2.85367109 | 1.482569324 | | 4 | 1.050078 | 13115.5 |
| p.Ser254Pro | 254 | P | 0.38417654 | 0.76624847 | 2.36717311 | 2.80735492 | 1.581238259 | | 4 | 1.406074 | 9333.75 |
| p.Ser254Cys | 254 | C | 0.43219222 | 0.47201271 | 3.37125581 | 2.33319921 | 1.652164986 | | 4 | 2.100058 | 2386.75 |
| p.Ser254Val | 254 | V | 0.20288426 | 1.02179938 | 2.98641094 | 2.42064499 | 1.65793489 | | 4 | 1.622805 | 7902.25 |
| p.Ser254Arg | 254 | R | 0.88331439 | 0.71973275 | 3.05784407 | 2.31170825 | 1.743149868 | | 4 | 1.27946 | 7160.5 |
| p.Ser254Asp | 254 | D | 0.30372048 | 1.20994913 | 2.29232163 | 3.1727053 | 1.744674134 | | 4 | 1.567156 | 7598 |
| p.Ser254Lys | 254 | K | 1.29319638 | 0.88101196 | 3.23446525 | 2.4150375 | 1.955927773 | | 4 | 1.146699 | 593.75 |
| p.Ser255Ile | 255 | I | 0.90319571 | 0.36018867 | 2.91235388 | -0.2981021 | 0.969409044 | | 4 | 1.919051 | 6633.5 |
| p.Ser255Gly | 255 | G | -0.0498267 | -0.3500533 | 2.63742992 | 1.81942775 | 1.014244426 | | 4 | 2.092202 | 6464.25 |
| p.Ser255Glu | 255 | E | -0.3797764 | 0.13178987 | 2.44376419 | 2.08956538 | 1.071335763 | | 4 | 1.969608 | 14355.5 |
| p.Ser255Pro | 255 | P | 0.3802342 | 0.31577028 | 2.08447979 | 1.79449936 | 1.143745908 | | 4 | 0.858985 | 29486.75 |
| p.Ser255Val | 255 | V | 0.0805512 | 0.09046282 | 2.4621333 | 2.16838687 | 1.200383547 | | 4 | 1.671664 | 27994 |
| p.Ser255Asp | 255 | D | 0.09028889 | 0.45904238 | 3.03464614 | 1.33846068 | 1.230609522 | | 4 | 1.720609 | 4700.5 |
| p.Ser255Lys | 255 | K | 1.31368793 | 0.12069556 | 2.03420148 | 1.54381652 | 1.253100374 | | 4 | 0.660215 | 13614.5 |
| p.Ser255Thr | 255 | T | 0.3972441 | 0.93043568 | 2.09857739 | 1.74237909 | 1.292159067 | | 4 | 0.594908 | 12831.25 |
| p.Ser255Arg | 255 | R | 0.66587579 | 0.84373179 | 2.72078358 | 1.2717381 | 1.375532315 | | 4 | 0.868966 | 25849 |
| p.Ser255Leu | 255 | L | 0.62828462 | 0.63531061 | 2.48045959 | 1.94039174 | 1.421111642 | | 4 | 0.879309 | 51426.75 |
| p.Ser255Met | 255 | M | 0.28590887 | 0.66878618 | 2.96410633 | 1.98602996 | 1.476207835 | | 4 | 1.514167 | 13873 |
| p.Ser255Asn | 255 | N | 1.46789872 | 0.97389004 | 1.7972661 | 1.82555952 | 1.516153595 | | 4 | 0.157045 | 5634.25 |
| p.Ser255Ala | 255 | A | 0.82010412 | 0.44082153 | 3.00219756 | 2.61551997 | 1.719660793 | | 4 | 1.630699 | 4423 |

| | | | | | | | | | | |
|---|---|---|---|---|---|---|---|---|---|---|
| p.Ser255Gln | 255 | Q | 0.87064526 | 0.52255594 | 3.01342595 | 2.75860572 | 1.791308217 | 4 | 1.628863 | 52108.75 |
| p.Thr257Lys | 257 | K | -0.3783554 | 0.43724838 | 1.90382863 | 0.13142047 | 0.523535526 | 4 | 0.959939 | 17526 |
| p.Thr257Cys | 257 | C | 0.20266179 | 0.83809453 | 2.06547384 | -0.6088883 | 0.624335453 | 4 | 1.273741 | 4383.5 |
| p.Thr257Arg | 257 | R | -0.0316744 | -0.4139584 | 2.462863 | 1.4807753 | 0.874501377 | 4 | 1.790581 | 37520.25 |
| p.Thr257His | 257 | H | -0.2933852 | 0.12921015 | 2.06228428 | 2.44346707 | 1.085394064 | 4 | 1.871332 | 9224 |
| p.Thr257Glu | 257 | E | 0.07520194 | 0.26169506 | 1.8786937 | 2.38883815 | 1.151107214 | 4 | 1.336662 | 14938 |
| p.Thr257Trp | 257 | W | -0.203294 | -0.0936618 | 3.09122967 | 1.85861058 | 1.163221123 | 4 | 2.549301 | 10223.5 |
| p.Thr257Phe | 257 | F | 1.154374 | -0.2502266 | 2.92599942 | 1.33371266 | 1.290964862 | 4 | 1.689698 | 3198 |
| p.Thr257Pro | 257 | P | 0.44288194 | 0.4328259 | 2.78235548 | 2.28005299 | 1.484529076 | 4 | 1.502773 | 38146.25 |
| p.Thr257Ser | 257 | S | 0.38771907 | 0.68543166 | 3.29102051 | 1.57497409 | 1.484786331 | 4 | 1.70438 | 13905.25 |
| p.Thr257Ile | 257 | I | 1.15134022 | 0.78427131 | 1.64519761 | 2.49491828 | 1.518931853 | 4 | 0.547781 | 6249.25 |
| p.Thr257Gly | 257 | G | 0.07512651 | 0.69970748 | 3.03190858 | 2.32111834 | 1.531965229 | 4 | 1.895875 | 54429.75 |
| p.Thr257Ala | 257 | A | -0.4098224 | 1.13434271 | 3.23685778 | 2.17254095 | 1.53347977 | 4 | 2.41521 | 22497.5 |
| p.Thr257Leu | 257 | L | 0.79307671 | 0.65217292 | 2.86352309 | 2.37987196 | 1.672161168 | 4 | 1.244454 | 38699 |
| p.Thr257Asn | 257 | N | 0.83483566 | 0.10985648 | 1.80735492 | 4.06847974 | 1.705131699 | 4 | 2.966061 | 1409.75 |
| p.Thr257Met | 257 | M | 0.5208783 | 0.76065994 | 2.55965683 | 3.17634841 | 1.75438587 | 4 | 1.72649 | 25197.75 |
| p.Thr257Gln | 257 | Q | 0.5083946 | 0.85056502 | 2.11164536 | 3.58611252 | 1.764179372 | 4 | 1.95062 | 10664.5 |
| p.Thr257Val | 257 | V | 0.30526069 | 0.97640562 | 2.97519661 | 3.18619565 | 1.860764641 | 4 | 2.066803 | 15809 |
| p.Ser261Glu | 261 | E | 0.60582111 | -0.6979071 | -0.2294818 | 1.43186212 | 0.277573574 | 4 | 0.882932 | 3624.75 |
| p.Ser261Met | 261 | M | -0.3292025 | 0.10771961 | 2.29923036 | -0.3429679 | 0.433694888 | 4 | 1.590567 | 14900.75 |
| p.Ser261Pro | 261 | P | -0.2520155 | -0.0503952 | 1.96625852 | 0.71010425 | 0.593488017 | 4 | 1.009187 | 33878 |
| p.Ser261Cys | 261 | C | -0.3712558 | 0.53417157 | 2.31070084 | 0.23582736 | 0.67736099 | 4 | 1.327617 | 7470.75 |
| p.Ser261Gln | 261 | Q | -0.290928 | 0.24576217 | 2.30017354 | 0.6158368 | 0.717711139 | 4 | 1.251551 | 24915.25 |
| p.Ser261Asp | 261 | D | -0.0421144 | 0.19177019 | 1.95503611 | 0.78800751 | 0.723174861 | 4 | 0.796581 | 12224.75 |
| p.Ser261Thr | 261 | T | -0.7952974 | 0.29626321 | 2.35198533 | 1.24441873 | 0.774342463 | 4 | 1.800753 | 20094.5 |
| p.Ser261Arg | 261 | R | 0.69210496 | 0.12713876 | 1.76618992 | 0.60904812 | 0.79862044 | 4 | 0.478121 | 17346 |
| p.Ser261Leu | 261 | L | 0.35299136 | 0.67602765 | 1.71776138 | 0.70838414 | 0.863791134 | 4 | 0.349863 | 16371.5 |
| p.Ser261Val | 261 | V | -0.3786306 | -0.3252113 | 2.0836002 | 2.13545278 | 0.878802773 | 4 | 2.020498 | 17532.25 |
| p.Ser261Ile | 261 | I | 0.46158009 | -0.115557 | 1.9609579 | 1.25612604 | 0.890776756 | 4 | 0.825228 | 4737.25 |
| p.Ser261Ala | 261 | A | 0.52380576 | 0.36806043 | 1.86842007 | 2.47007055 | 1.307589202 | 4 | 1.054308 | 34851.75 |
| p.Ser261Lys | 261 | K | 0.33317021 | 0.10050754 | 2.9895984 | 1.87474347 | 1.324504905 | 4 | 1.852071 | 19401 |
| p.Ser261Gly | 261 | G | 0.44303964 | 0.66446927 | 2.61268698 | 1.63371085 | 1.338476685 | 4 | 0.988956 | 22138.25 |
| p.Ser261Asn | 261 | N | 0.07102199 | 0.15366911 | 2.55623821 | 2.73754026 | 1.379617392 | 4 | 2.147921 | 8506 |
| p.Ser261Tyr | 261 | Y | 0.93464356 | 0.85996526 | 2.81690923 | 1.5740906 | 1.546402161 | 4 | 0.820133 | 9756.75 |
| p.Arg273Gly | 273 | G | -0.3467953 | -0.6814523 | 1.87316998 | 0.46484106 | 0.327440871 | 4 | 1.293539 | 10210.25 |

| | | | | | | | | | | |
|---|---|---|---|---|---|---|---|---|---|---|
| p.Arg273Asn | 273 | N | -0.131167 | 0.23968503 | 2.30078933 | -0.5485205 | 0.465196715 | 4 | 1.601176 | 3142.25 |
| p.Arg273Trp | 273 | W | 1.02089522 | -0.4484237 | 0.74075717 | 0.64198245 | 0.488802795 | 4 | 0.416154 | 5680.25 |
| p.Arg273Pro | 273 | P | -0.3544567 | -0.3966829 | 2.08869224 | 1.04253151 | 0.595021038 | 4 | 1.438768 | 19206 |
| p.Arg273Phe | 273 | F | -0.5277412 | 0.32884548 | 1.63399084 | 1.03991437 | 0.618752365 | 4 | 0.86886 | 8748 |
| p.Arg273Met | 273 | M | -0.7021792 | 0.0893791 | 1.9177919 | 1.17380247 | 0.61969858 | 4 | 1.340226 | 11957.75 |
| p.Arg273Leu | 273 | L | -0.4171304 | -0.340045 | 1.99887421 | 1.30550255 | 0.636800342 | 4 | 1.455802 | 63748 |
| p.Arg273Val | 273 | V | -0.0329566 | -0.2520831 | 2.1250434 | 1.26459053 | 0.77614856 | 4 | 1.256668 | 27356 |
| p.Arg273Ser | 273 | S | 0.17039227 | -0.3816374 | 2.3070121 | 1.51495846 | 0.902681369 | 4 | 1.510917 | 10993.75 |
| p.Arg273Tyr | 273 | Y | -0.2771941 | -0.0857536 | 3.22106385 | 0.75477867 | 0.903223706 | 4 | 2.588627 | 10856.75 |
| p.Arg273Thr | 273 | T | -1.0291463 | 1.24415294 | 1.39592868 | 2.10302401 | 0.92848982 | 4 | 1.843337 | 4278.5 |
| p.Arg273Gln | 273 | Q | 0.34342503 | 0.16873229 | 1.88772318 | 1.43535748 | 0.958809497 | 4 | 0.697633 | 12730.75 |
| p.Arg273Ile | 273 | I | 0.75812091 | 0.08945733 | 2.35606443 | 0.93605918 | 1.034925463 | 4 | 0.908571 | 11728.75 |
| p.Arg273Ala | 273 | A | 0.55068751 | -0.7810582 | 2.70193862 | 2.58607014 | 1.264409522 | 4 | 2.835538 | 12011.25 |
| p.Ser274Ile | 274 | I | -0.507989 | 0.70368715 | 2.57543973 | 0.34549657 | 0.779158616 | 4 | 1.692378 | 9471.5 |
| p.Ser274Gly | 274 | G | 0.41248992 | 0.3354862 | 3.37280926 | -0.3029732 | 0.954453058 | 4 | 2.702137 | 6772.25 |
| p.Ser274Pro | 274 | P | -0.5119581 | -0.2196547 | 2.71885686 | 1.84744373 | 0.958671942 | 4 | 2.479791 | 36770 |
| p.Ser274Leu | 274 | L | 0.30895284 | 0.50925171 | 2.75141187 | 1.23521646 | 1.20120822 | 4 | 1.226404 | 51309.25 |
| p.Ser274Met | 274 | M | 0.52708104 | 0.38540141 | 2.47143356 | 1.44794124 | 1.207964311 | 4 | 0.931385 | 17962 |
| p.Ser274Gln | 274 | Q | 0.51401626 | 0.19106577 | 2.82156878 | 1.48656844 | 1.253304813 | 4 | 1.396254 | 12753.5 |
| p.Ser274Thr | 274 | T | 0.41498974 | 0.26083791 | 2.15210526 | 2.39065071 | 1.304645904 | 4 | 1.259539 | 30122 |
| p.Ser274Arg | 274 | R | 0.39253751 | 0.51478187 | 2.99044117 | 1.82793672 | 1.431424319 | 4 | 1.502425 | 32701 |
| p.Ser274Asn | 274 | N | 0.29371717 | 1.22483422 | 2.472821 | 2.25776979 | 1.562285545 | 4 | 1.011971 | 10262.25 |
| p.Ser274Cys | 274 | C | 1.06761623 | 0.32852436 | 2.27718978 | 2.61831192 | 1.572910572 | 4 | 1.130898 | 9969.25 |
| p.Ser274Ala | 274 | A | 1.01002572 | 0.93965814 | 2.10449667 | 2.58294803 | 1.65928214 | 4 | 0.663589 | 27378.5 |
| p.Ser274Glu | 274 | E | 0.91960165 | 0.50664879 | 3.09050137 | 2.30078109 | 1.704383224 | 4 | 1.442488 | 28146 |
| p.Ser274Lys | 274 | K | 1.13296079 | 0.57550217 | 2.99262047 | 2.42655817 | 1.781910402 | 4 | 1.252649 | 5624.5 |
| p.Ser274Val | 274 | V | 0.53674384 | 0.81431614 | 2.84931332 | 3.0780884 | 1.819615425 | 4 | 1.766806 | 18511.75 |
| p.Ser274His | 274 | H | 0.83004273 | 1.1304368 | 2.78068697 | 3.32419142 | 2.01633948 | 4 | 1.495609 | 25412.75 |
| p.Lys277Val | 277 | V | 0.17489879 | 0.80847764 | 1.95314619 | 0.43712106 | 0.843410919 | 4 | 0.614904 | 8918 |
| p.Lys277Ser | 277 | S | 0.01931956 | 0.70578331 | 1.71234181 | 1.20291341 | 0.910089522 | 4 | 0.521522 | 11541.75 |
| p.Lys277Arg | 277 | R | 0.08363098 | 0.04787663 | 2.73476852 | 1.38560105 | 1.062969293 | 4 | 1.629507 | 13946 |
| p.Lys277Asn | 277 | N | -0.2589833 | 1.66990241 | 0.44349757 | 2.43937946 | 1.073449032 | 4 | 1.464579 | 5952.75 |
| p.Lys277His | 277 | H | 0.43079793 | 0.90721361 | 1.82359556 | 1.25050401 | 1.103027776 | 4 | 0.343734 | 13789.25 |
| p.Lys277Met | 277 | M | 0.20457938 | 0.39165172 | 3.08855044 | 1.08037978 | 1.191290328 | 4 | 1.741639 | 17684.5 |
| p.Lys277Gly | 277 | G | 0.22212646 | 0.2925926 | 2.92550627 | 1.61713251 | 1.26433946 | 4 | 1.638146 | 6025.5 |

| | | | | | | | | | | |
|---|---|---|---|---|---|---|---|---|---|---|
| p.Lys277Asp | 277 | D | 0.65078127 | 0.55176316 | 2.47108363 | 1.64762777 | 1.330313957 | 4 | 0.823317 | 16687.25 |
| p.Lys277Ala | 277 | A | 1.04154308 | 1.09803684 | 2.05975474 | 1.20220463 | 1.350384822 | 4 | 0.228075 | 22382.75 |
| p.Lys277Pro | 277 | P | 0.10554634 | 1.01431099 | 2.0852626 | 2.24079318 | 1.361478278 | 4 | 0.998316 | 20585.25 |
| p.Lys277Ile | 277 | I | 1.08092 | 1.04892636 | 2.22597231 | 1.2988796 | 1.413674567 | 4 | 0.305591 | 6078 |
| p.Lys277Glu | 277 | E | 0.57352156 | 0.70589735 | 2.88530123 | 1.89140851 | 1.514032162 | 4 | 1.186811 | 14300.5 |
| p.Lys277Thr | 277 | T | 0.54169761 | 0.31801796 | 2.38148705 | 2.92941652 | 1.542654783 | 4 | 1.709466 | 37127 |
| p.Lys277Tyr | 277 | Y | 0.02587489 | 0.47090317 | 3.51096192 | 2.86603083 | 1.718442701 | 4 | 2.983741 | 6026.5 |
| p.Lys277Leu | 277 | L | 1.20273604 | 0.51664113 | 3.31738417 | 3.15087806 | 2.046909851 | 4 | 1.962401 | 13211.25 |
| p.Ser289Pro | 289 | P | 0.36505534 | 0.0485866 | 1.98831819 | -0.9726927 | 0.35731687 | 4 | 1.508155 | 3998.25 |
| p.Ser289Gly | 289 | G | -0.3714588 | 0.00178111 | 2.71010425 | -0.1479506 | 0.548118991 | 4 | 2.100934 | 6229 |
| p.Ser289Trp | 289 | W | 0.05268299 | 0.33985 | 2.65896308 | 1.44452294 | 1.124004753 | 4 | 1.407153 | 8718 |
| p.Ser289Arg | 289 | R | -0.0268985 | 1.07423062 | 2.95659321 | 1.04298317 | 1.261727129 | 4 | 1.53871 | 8357 |
| p.Ser289Gln | 289 | Q | 1.05767277 | 1.33180961 | 2.83241238 | -0.0347654 | 1.296782333 | 4 | 1.396526 | 9592.5 |
| p.Ser289Leu | 289 | L | 0.58746067 | 0.94775291 | 2.02711613 | 1.72317098 | 1.321375173 | 4 | 0.445911 | 11149.75 |
| p.Ser289Lys | 289 | K | -0.4178715 | 0.88599537 | 3.10336384 | 2.03634185 | 1.401957396 | 4 | 2.29174 | 4653.75 |
| p.Ser289Met | 289 | M | 1.09965688 | 1.20424323 | 2.92081297 | 0.99680466 | 1.555379436 | 4 | 0.835798 | 9658.75 |
| p.Ser289Thr | 289 | T | 0.53852207 | 0.55136953 | 3.63630938 | 2.19101461 | 1.729303898 | 4 | 2.218446 | 3302 |
| p.Ser289Glu | 289 | E | 1.0413511 | 0.74644213 | 3.45943162 | 2.45003292 | 1.924314443 | 4 | 1.599991 | 3679.75 |
| p.Ser289Ala | 289 | A | 0.41248179 | 1.4786536 | 3.94381716 | 1.97736903 | 1.953080396 | 4 | 2.187383 | 5626.5 |
| p.Thr293His | 293 | H | -0.0131754 | -0.5032086 | 1.30518483 | 0.45319643 | 0.310499331 | 4 | 0.592216 | 9661 |
| p.Thr293Lys | 293 | K | -0.9794324 | -0.0190908 | 1.45892514 | 0.90262459 | 0.340756649 | 4 | 1.146129 | 6344.5 |
| p.Thr293Arg | 293 | R | -0.0595762 | 0.00741747 | 2.13626569 | 1.56149349 | 0.911400102 | 4 | 1.227632 | 16163.5 |
| p.Thr293Gly | 293 | G | 0.2407431 | 1.0207829 | 2.31256268 | 0.93123132 | 1.12633 | 4 | 0.746872 | 28093.75 |
| p.Thr293Leu | 293 | L | -0.1200666 | 0.90646931 | 2.5849625 | 1.68706069 | 1.264606482 | 4 | 1.322463 | 2819.5 |
| p.Thr293Ile | 293 | I | 0.52371334 | 0.30389505 | 3.04240557 | 1.78379145 | 1.41345135 | 4 | 1.604464 | 7218.5 |
| p.Thr293Tyr | 293 | Y | 1.24731503 | 0.6544914 | 2.3991189 | 2.36462374 | 1.666387269 | 4 | 0.741328 | 9189.75 |
| p.Thr293Ser | 293 | S | 0.42080889 | 0.62884242 | 2.20863512 | 3.4334906 | 1.672944262 | 4 | 2.014827 | 13537.25 |
| p.Thr293Val | 293 | V | 0.48219634 | 0.83484554 | 3.1845347 | 2.75716177 | 1.814684584 | 4 | 1.833454 | 12836.25 |
| p.Thr293Ala | 293 | A | 1.59422542 | 0.32474037 | 3.85315861 | 2.6592947 | 2.107854775 | 4 | 2.264495 | 4319.5 |
| p.Thr293Met | 293 | M | 0.02132851 | 3.02661173 | 2.89450687 | 2.50250034 | 2.111236863 | 4 | 1.990742 | 2490 |
| p.Thr293Pro | 293 | P | 1.61851506 | 1.59708606 | 3.20243781 | 2.82017896 | 2.309554474 | 4 | 0.681042 | 3389.5 |
| p.Leu295Gly | 295 | G | 0.51628128 | 0.26778959 | 2.24814887 | 0.80374467 | 0.958991102 | 4 | 0.786594 | 40213.75 |
| p.Leu295Thr | 295 | T | -0.0161756 | 0.59939262 | 2.3328914 | 1.20017629 | 1.029071181 | 4 | 1.00213 | 54938.25 |
| p.Leu295Pro | 295 | P | -0.1955101 | 0.3713235 | 2.18455692 | 2.02627497 | 1.09666132 | 4 | 1.414507 | 27549.75 |
| p.Leu295Asn | 295 | N | 0.35811419 | 0.55076905 | 2.41618568 | 1.20402278 | 1.132272925 | 4 | 0.863683 | 14286.5 |

| | | | | | | | | | | |
|---|---|---|---|---|---|---|---|---|---|---|
| p.Leu295Arg | 295 | R | 0.21437409 | 0.07005968 | 2.26932067 | 2.21237571 | 1.191532537 | 4 | 1.472096 | 26385.25 |
| p.Leu295Ala | 295 | A | 0.40039023 | 0.39706359 | 2.83113791 | 1.19935489 | 1.206986655 | 4 | 1.314833 | 26270.75 |
| p.Leu295Ile | 295 | I | 0.81563836 | 0.43993168 | 2.83058723 | 0.8215247 | 1.226920491 | 4 | 1.174866 | 3320 |
| p.Leu295Val | 295 | V | 0.74282198 | 0.60454576 | 2.49000134 | 1.40847374 | 1.311460704 | 4 | 0.740483 | 44104 |
| p.Leu295Met | 295 | M | 0.28497224 | 0.1811911 | 3.06801159 | 2.2967081 | 1.457720757 | 4 | 2.100601 | 24485.25 |
| p.Leu295Trp | 295 | W | 0.24399763 | 0.27154591 | 2.6520767 | 3.21183864 | 1.594864718 | 4 | 2.436105 | 1891.25 |
| p.Leu295Ser | 295 | S | 0.83217943 | 0.5192523 | 2.98523933 | 2.11578375 | 1.613113701 | 4 | 1.313932 | 23357.5 |
| p.Leu295Asp | 295 | D | -0.0076968 | 1.04874044 | 3.63269511 | 2.27510224 | 1.737210247 | 4 | 2.466961 | 10893.25 |
| p.Leu295His | 295 | H | 1.38083602 | 0.54936998 | 1.79646661 | 3.35641647 | 1.770772268 | 4 | 1.386267 | 6133.25 |
| p.Leu295Glu | 295 | E | 1.1333231 | 0.36164079 | 2.81069682 | 3.18153629 | 1.871799249 | 4 | 1.807622 | 26850 |
| p.Ser296Asn | 296 | N | -0.0323068 | -0.2038658 | 2.1004019 | -0.257878 | 0.401587821 | 4 | 1.291901 | 6099.25 |
| p.Ser296Met | 296 | M | -1.9156078 | 0.56660068 | 1.65507253 | 1.61009871 | 0.479041027 | 4 | 2.80145 | 12487 |
| p.Ser296Pro | 296 | P | 0.26748221 | 0.34012209 | 2.37475864 | 0.75783947 | 0.935050603 | 4 | 0.967917 | 49872.25 |
| p.Ser296Val | 296 | V | 0.27361576 | 0.11606958 | 1.71739401 | 1.81184411 | 0.979730865 | 4 | 0.827023 | 29753.5 |
| p.Ser296Leu | 296 | L | 0.69953459 | 0.43979155 | 1.7574297 | 1.41464626 | 1.077850523 | 4 | 0.375167 | 15492.5 |
| p.Ser296Glu | 296 | E | -0.0747678 | 0.05707558 | 3.09754895 | 2.10774986 | 1.296901653 | 4 | 2.439484 | 7131 |
| p.Ser296Thr | 296 | T | 0.10318191 | 0.68280437 | 1.81490936 | 2.64870197 | 1.312399402 | 4 | 1.298939 | 24666.75 |
| p.Ser296Lys | 296 | K | -0.2471546 | 0.58126947 | 3.42019211 | 1.72831672 | 1.370655913 | 4 | 2.522987 | 20216.25 |
| p.Ser296Ala | 296 | A | 0.14651832 | 0.04415332 | 2.80961797 | 3.07818398 | 1.519618399 | 4 | 2.718542 | 16673 |
| p.Ser296Gln | 296 | Q | 1.08051704 | -0.0388074 | 3.24547603 | 2.07584376 | 1.590757346 | 4 | 1.963076 | 7098 |
| p.Ser296Gly | 296 | G | 0.34246646 | 1.01121337 | 2.76836079 | 2.37316129 | 1.623800479 | 4 | 1.296213 | 28231.75 |
| p.Ser296Asp | 296 | D | 0.89289344 | 0.92709042 | 3.1948534 | 1.5179838 | 1.633205266 | 4 | 1.166227 | 5861 |
| p.Ser296Arg | 296 | R | 0.18280379 | 1.23293237 | 3.17326071 | 2.24670624 | 1.708925779 | 4 | 1.663034 | 9142.75 |
| p.Ser296Trp | 296 | W | 1.39406733 | 0.79041265 | 3.40839219 | 3.46159188 | 2.263616011 | 4 | 1.890701 | 4092.5 |
| p.Thr303Arg | 303 | R | 0.53822149 | 0.04783275 | 2.30433403 | 0.17521653 | 0.766401201 | 4 | 1.094381 | 14969.5 |
| p.Thr303Pro | 303 | P | 0.22245705 | 0.65353358 | 2.15757796 | 0.3681898 | 0.850439597 | 4 | 0.791437 | 39937.75 |
| p.Thr303Phe | 303 | F | -0.1027821 | 0.63255947 | 2.53273254 | 0.76497262 | 0.956870623 | 4 | 1.249402 | 7216.25 |
| p.Thr303Ala | 303 | A | 1.0963071 | 0.78984337 | 2.49892267 | -0.0598212 | 1.081312995 | 4 | 1.132328 | 11352.25 |
| p.Thr303Asp | 303 | D | 0.2932297 | 0.89587513 | 2.05664358 | 1.33237635 | 1.14453119 | 4 | 0.55126 | 16005.5 |
| p.Thr303Gly | 303 | G | 0.41994741 | 0.3491546 | 2.67850327 | 1.73976423 | 1.296842377 | 4 | 1.257408 | 11599.25 |
| p.Thr303Gln | 303 | Q | 0.54648835 | 0.88552999 | 2.4813925 | 1.29026894 | 1.300919947 | 4 | 0.711782 | 15276.25 |
| p.Thr303Asn | 303 | N | 0.0911079 | 0.25770096 | 2.54711915 | 2.93467375 | 1.457650441 | 4 | 2.225286 | 6504 |
| p.Thr303Ile | 303 | I | 0.72267637 | 0.44011546 | 2.27889071 | 2.40991372 | 1.462899065 | 4 | 1.052232 | 32107 |
| p.Thr303Leu | 303 | L | 0.12869423 | 0.55318398 | 3.11485571 | 2.3310044 | 1.531934579 | 4 | 2.023729 | 67381.25 |
| p.Thr303His | 303 | H | 0.43137456 | 0.86169063 | 2.6698974 | 2.28189343 | 1.561214007 | 4 | 1.171476 | 37506.25 |

| | | | | | | | | | | |
|---|---|---|---|---|---|---|---|---|---|---|
| p.Thr303Val | 303 | V | 1.03283611 | 0.54119432 | 1.96664911 | 2.83645109 | 1.594282658 | 4 | 1.035285 | 30519 |
| p.Thr303Tyr | 303 | Y | 0.6518069 | 0.09098755 | 3.29308841 | 2.61656716 | 1.663112505 | 4 | 2.353404 | 12956.25 |
| p.Thr303Ser | 303 | S | 0.23020931 | 0.85235952 | 2.74753076 | 3.36850361 | 1.799650798 | 4 | 2.240094 | 28910 |
| p.Thr303Met | 303 | M | 0.46456705 | 1.01996521 | 2.85138162 | 3.3315297 | 1.916860897 | 4 | 1.929399 | 24459.5 |
| p.Thr303Cys | 303 | C | 2.14834792 | 1.05030508 | 3.16099188 | 1.79795622 | 2.039400276 | 4 | 0.768814 | 1525.25 |
| p.Lys308Leu | 308 | L | 0.6052871 | 1.11058473 | 0.95053132 | 0.3280542 | 0.748614338 | 4 | 0.123069 | 9292.25 |
| p.Lys308Asn | 308 | N | -0.2453519 | 0.49729348 | 2.09202914 | 1.02836611 | 0.843084201 | 4 | 0.966152 | 17108.5 |
| p.Lys308Gln | 308 | Q | -0.7248928 | 1.01400686 | 1.97198562 | 1.68732422 | 0.987105985 | 4 | 1.463986 | 1728.5 |
| p.Lys308Arg | 308 | R | 1.57777964 | 1.07438477 | 1.82729791 | -0.2656326 | 1.053457433 | 4 | 0.871393 | 9049.75 |
| p.Lys308Ser | 308 | S | 0.9247583 | -0.0856894 | 2.56331394 | 0.91807571 | 1.080114623 | 4 | 1.203124 | 20912.25 |
| p.Lys308His | 308 | H | 0.59927644 | 0.23320601 | 2.90549601 | 1.15886089 | 1.224209838 | 4 | 1.401208 | 12429.25 |
| p.Lys308Met | 308 | M | 0.44231733 | 0.55104906 | 2.7132559 | 1.56834606 | 1.31874209 | 4 | 1.121481 | 30790.25 |
| p.Lys308Pro | 308 | P | 0.79438571 | 0.56302657 | 2.60090404 | 1.34519155 | 1.325876968 | 4 | 0.830164 | 19808.25 |
| p.Lys308Val | 308 | V | 0.19773388 | 0.32475534 | 2.69943958 | 2.46958772 | 1.422879131 | 4 | 1.810687 | 30079.5 |
| p.Lys308Ala | 308 | A | 0.63241882 | 0.51324315 | 2.66956041 | 1.97394701 | 1.447292346 | 4 | 1.10259 | 32314.75 |
| p.Lys308Glu | 308 | E | 0.32289723 | 0.53143275 | 3.06500999 | 2.22971507 | 1.537263762 | 4 | 1.766626 | 40587.75 |
| p.Lys308Thr | 308 | T | 0.87926499 | 0.70010134 | 2.9452997 | 1.74521775 | 1.567470944 | 4 | 1.051988 | 38162.75 |
| p.Lys308Gly | 308 | G | 0.95155238 | 0.47952131 | 3.53098971 | 2.62449086 | 1.896638568 | 4 | 2.034094 | 37103.25 |
| p.Lys308Cys | 308 | C | -0.1322172 | 1.67585408 | 3.4429435 | 3.00791759 | 1.9986245 | 4 | 2.583132 | 5817.25 |
| p.Lys308Asp | 308 | D | 1.14367036 | 1.24932926 | 3.37608386 | 2.43050489 | 2.049897095 | 4 | 1.12193 | 14201.75 |
| p.Arg313Tyr | 313 | Y | -0.9280139 | -0.0421162 | 2.9328858 | -0.2022631 | 0.440123169 | 4 | 2.910293 | 2819.75 |
| p.Arg313His | 313 | H | 0.09539765 | 0.95803383 | 2.66035557 | 0.2576296 | 0.992854164 | 4 | 1.375919 | 14355.75 |
| p.Arg313Thr | 313 | T | 0.94654881 | 0.02352764 | 1.52356196 | 1.6362142 | 1.032463153 | 4 | 0.543675 | 6038.5 |
| p.Arg313Ile | 313 | I | 0.21829857 | 0.07766066 | 3.09718042 | 0.80650497 | 1.049911155 | 4 | 1.96247 | 22354.25 |
| p.Arg313Glu | 313 | E | 0.79956522 | -0.5884985 | 3.33390074 | 0.78526115 | 1.082557147 | 4 | 2.676481 | 4682.25 |
| p.Arg313Ser | 313 | S | -0.4289121 | 0.5050307 | 2.04026387 | 2.36428655 | 1.120167244 | 4 | 1.72415 | 18813 |
| p.Arg313Gly | 313 | G | 0.54519441 | 0.55887743 | 3.12928302 | 0.77304641 | 1.251600318 | 4 | 1.57786 | 10872.25 |
| p.Arg313Leu | 313 | L | 0.32166356 | 0.68375933 | 2.33360741 | 1.85053777 | 1.297392018 | 4 | 0.902768 | 56486 |
| p.Arg313Pro | 313 | P | 0.42924001 | 0.47053398 | 2.94194096 | 2.56708993 | 1.602201222 | 4 | 1.794141 | 17733.5 |
| p.Arg313Asp | 313 | D | 0.27546556 | 0.91292244 | 2.49362219 | 2.83046698 | 1.628119292 | 4 | 1.511971 | 8838.75 |
| p.Arg313Val | 313 | V | 0.78587519 | 0.33641155 | 2.75126719 | 3.19556613 | 1.767280017 | 4 | 2.006257 | 21471.5 |
| p.Arg313Asn | 313 | N | 0.4022027 | 1.07228886 | 2.92184094 | 3.86133629 | 2.064417195 | 4 | 2.570456 | 7892.75 |
| p.Arg313Cys | 313 | C | 1.85059202 | 1.55184323 | 3.55799545 | 4.3286941 | 2.8222812 | 4 | 1.789583 | 2971 |
| p.Ala326Val | 326 | V | 0.5931519 | 0.1585359 | 1.84608732 | -0.9142701 | 0.420876246 | 4 | 1.304115 | 4592.75 |
| p.Arg333Leu | 333 | L | 0.71633075 | 1.00555239 | 3.01810181 | 1.59088733 | 1.582718071 | 4 | 1.048047 | 3442.75 |

| | | | | | | | | | |
|---|---|---|---|---|---|---|---|---|---|
| p.Ala335Val | 335 | V | 1.13683421 | 0.96538624 | 1.9361849 | 2.65354959 | 1.672988737 | 4 | 0.606312 | 2306.25 |
| p.Val336Ile | 336 | I | 0.82249063 | 1.08381999 | 2.28239973 | 3.23896665 | 1.85691925 | 4 | 1.252938 | 4693.25 |
| p.Thr341His | 341 | H | 0.77346388 | 0.40373441 | 0.52643642 | -0.1029572 | 0.400169383 | 4 | 0.136147 | 7571.5 |
| p.Thr341Ile | 341 | I | -0.0463556 | 0.17665966 | 3.15549793 | -0.2377676 | 0.762008576 | 4 | 2.57481 | 4702 |
| p.Thr341Lys | 341 | K | -0.2111168 | 0.08789879 | 2.45053618 | 1.08361964 | 0.852734464 | 4 | 1.44101 | 16869.25 |
| p.Thr341Val | 341 | V | -0.2049429 | 0.08693727 | 1.9588948 | 1.69915246 | 0.885010401 | 4 | 1.213658 | 19309 |
| p.Thr341Gly | 341 | G | -0.5094481 | 0.0892797 | 2.97476271 | 1.1693778 | 0.930993017 | 4 | 2.339058 | 8920.75 |
| p.Thr341Asp | 341 | D | 1.73011554 | -0.3807676 | 2.49304001 | 0.05588602 | 0.974568493 | 4 | 1.852507 | 4691 |
| p.Thr341Ala | 341 | A | 0.37944037 | 1.09616882 | 3.169925 | -0.3497774 | 1.073939197 | 4 | 2.300983 | 15993.75 |
| p.Thr341Glu | 341 | E | -0.0035701 | 0.31044797 | 2.35689781 | 1.78876219 | 1.113134457 | 4 | 1.298252 | 16585.5 |
| p.Thr341Arg | 341 | R | 0.04777971 | 0.31265342 | 2.57257878 | 1.88783226 | 1.205211043 | 4 | 1.490658 | 26115.25 |
| p.Thr341Met | 341 | M | 0.6867375 | 0.78803641 | 2.38891243 | 1.10314745 | 1.241708447 | 4 | 0.616362 | 28733.75 |
| p.Thr341Pro | 341 | P | 0.38920078 | -0.5578573 | 2.43686386 | 3.11788774 | 1.346523776 | 4 | 2.956569 | 3481 |
| p.Thr341Leu | 341 | L | -0.2969499 | 1.05610845 | 2.36767127 | 2.40122714 | 1.382014234 | 4 | 1.64515 | 15577.75 |
| p.Thr341Ser | 341 | S | 0.26475356 | 0.27876352 | 3.06258817 | 2.81874393 | 1.606212296 | 4 | 2.384298 | 20254 |
| p.Thr341Asn | 341 | N | 1.05780876 | 1.16252032 | 1.8756502 | 2.55012713 | 1.661526603 | 4 | 0.482981 | 16159.5 |
| p.Arg343Leu | 343 | L | -0.2955914 | -0.4770888 | 2.53787736 | 0.69848228 | 0.615919868 | 4 | 1.908753 | 9683 |
| p.Arg343Pro | 343 | P | -0.2651733 | -0.3973355 | 1.67305383 | 1.95935802 | 0.742475758 | 4 | 1.553768 | 7132.75 |
| p.Arg343Ser | 343 | S | -0.5269304 | 0.63270858 | 2.98089118 | 0.35614381 | 0.860703296 | 4 | 2.242429 | 4102 |
| p.Arg343Met | 343 | M | -0.1557426 | 0.37239709 | 2.12988028 | 1.39517708 | 0.935427952 | 4 | 1.048581 | 6862.25 |
| p.Arg343Glu | 343 | E | 1.11670522 | 0.05518292 | 1.91634207 | 2.00552123 | 1.273437858 | 4 | 0.819327 | 8081.25 |
| p.Arg343Cys | 343 | C | 0.1134864 | 0.93652094 | 3.34452952 | 1.27627698 | 1.417703462 | 4 | 1.888393 | 4845.25 |
| p.Arg343Ala | 343 | A | -0.0728329 | 0.27456066 | 3.33038967 | 2.35852015 | 1.4726594 | 4 | 2.686633 | 3046.25 |
| p.Arg343Gln | 343 | Q | 0.24419899 | 1.40355263 | 2.17703189 | 2.09769667 | 1.480620044 | 4 | 0.80015 | 5386.5 |
| p.Arg343Lys | 343 | K | 1.26216051 | 1.04309346 | 2.24285652 | 3.1989172 | 1.936756926 | 4 | 0.980153 | 10573.75 |
| p.Arg343Val | 343 | V | 0.58590698 | 1.11496571 | 3.79385066 | 3.06316202 | 2.139471344 | 4 | 2.351116 | 4064.25 |
| p.Ser344Gly | 344 | G | -0.522481 | 0.72317384 | 1.99291058 | 1.03115452 | 0.806189494 | 4 | 1.077058 | 4346.25 |
| p.Ser344Gln | 344 | Q | -0.6147098 | 0.57575302 | 2.02895137 | 1.72315979 | 0.928288587 | 4 | 1.449468 | 5581.75 |
| p.Ser344Pro | 344 | P | -0.5153016 | 0.57210171 | 2.26041846 | 1.6601718 | 0.994347587 | 4 | 1.501197 | 28485 |
| p.Ser344Arg | 344 | R | 0.4779946 | 0.47891006 | 0.97760548 | 2.14963608 | 1.021036554 | 4 | 0.621473 | 15836.5 |
| p.Ser344Met | 344 | M | 0.67136206 | 0.55044193 | 2.53465742 | 0.79413134 | 1.137648188 | 4 | 0.877291 | 27974 |
| p.Ser344Thr | 344 | T | 0.79554422 | 0.7773339 | 2.18255512 | 1.33057564 | 1.271502221 | 4 | 0.434748 | 7596 |
| p.Ser344Val | 344 | V | 0.7902852 | 0.25181334 | 2.72375128 | 1.47168241 | 1.309383056 | 4 | 1.138232 | 11670 |
| p.Ser344Leu | 344 | L | 0.35730496 | 0.75375279 | 2.87345989 | 2.15145234 | 1.533992495 | 4 | 1.389599 | 49498 |
| p.Ser344Ile | 344 | I | 0.36464244 | 0.74471123 | 3.23759483 | 2.16828278 | 1.62880782 | 4 | 1.75299 | 13793.5 |

| | | | | | | | | | | |
|---|---|---|---|---|---|---|---|---|---|---|
| p.Ser344His | 344 | H | 1.46298944 | 0.43851307 | 2.90138412 | 2.50599567 | 1.827220573 | 4 | 1.225245 | 4319.25 |
| p.Glu349Lys | 349 | K | 0.99729579 | 2.98244163 | 3.91560781 | 3.7062688 | 2.900403507 | 4 | 1.769536 | 1386.25 |
| p.His350Asn | 350 | N | 1.07846124 | -0.12755 | 2.357552 | -0.2783012 | 0.757540508 | 4 | 1.50646 | 890 |
| p.Tyr355Lys | 355 | K | -0.3066988 | 0.47139994 | 0.90223995 | -1.0256866 | 0.010313608 | 4 | 0.727309 | 3981 |
| p.Tyr355Cys | 355 | C | 0.87981817 | 1.2907308 | 1.17319272 | -1.3116197 | 0.508030486 | 4 | 1.501471 | 4770.75 |
| p.Tyr355Pro | 355 | P | -0.4386904 | -0.4763709 | 2.17875258 | 1.16882914 | 0.608130094 | 4 | 1.684405 | 21946.75 |
| p.Tyr355Gln | 355 | Q | -0.1860559 | 0.10155534 | 2.7080209 | 0.37555906 | 0.749769859 | 4 | 1.756911 | 6068.75 |
| p.Tyr355Arg | 355 | R | 0.62352608 | -0.0566135 | 2.56215891 | 1.35698545 | 1.121514224 | 4 | 1.255627 | 21457.25 |
| p.Tyr355Asp | 355 | D | 0.04794528 | 0.35928151 | 2.00971515 | 2.2059043 | 1.15571156 | 4 | 1.231225 | 5247.25 |
| p.Tyr355Leu | 355 | L | -0.0914477 | 0.4032457 | 2.51652804 | 2.04213057 | 1.217614142 | 4 | 1.581281 | 28295.5 |
| p.Tyr355Ala | 355 | A | 0.09656258 | -0.0065207 | 2.75125155 | 2.33986955 | 1.295290752 | 4 | 2.11421 | 15718 |
| p.Tyr355Phe | 355 | F | 0.06458789 | 0.27724645 | 2.24619288 | 2.69412726 | 1.320538618 | 4 | 1.803151 | 15647.75 |
| p.Tyr355Val | 355 | V | 0.50916715 | -0.4751493 | 2.31948077 | 3.15912634 | 1.378156228 | 4 | 2.749277 | 10232.75 |
| p.Tyr355Thr | 355 | T | 0.52552523 | 0.65049447 | 2.67155019 | 1.66560522 | 1.378293776 | 4 | 1.003989 | 14923.75 |
| p.Tyr355Ser | 355 | S | -0.2744345 | 0.83757433 | 2.36748557 | 2.61855057 | 1.38729398 | 4 | 1.846767 | 11534.75 |
| p.Tyr355Gly | 355 | G | 0.01241799 | 1.38877772 | 2.88014574 | 1.47293588 | 1.438569332 | 4 | 1.371903 | 21419 |
| p.Tyr355Glu | 355 | E | 0.70939971 | 0.8106794 | 3.55136953 | 1.291317 | 1.590691409 | 4 | 1.772993 | 6564.25 |
| p.Tyr355Ile | 355 | I | 0.50138933 | 0.40263021 | 2.72656631 | 3.91701481 | 1.886900166 | 4 | 2.983034 | 14367.75 |
| p.Lys356Pro | 356 | P | -0.0567519 | 0.06796331 | 2.04847707 | 0.04460262 | 0.526072778 | 4 | 1.033026 | 18022.25 |
| p.Lys356Val | 356 | V | 0.20732008 | 0.53440467 | 2.68667344 | -1.049961 | 0.594609298 | 4 | 2.411654 | 10629 |
| p.Lys356Leu | 356 | L | -0.1512417 | -0.0647091 | 1.91753784 | 1.97179086 | 0.918344472 | 4 | 1.406182 | 11054.75 |
| p.Lys356Met | 356 | M | 0.18226431 | 0.812373 | 2.31520223 | 0.43050062 | 0.935085038 | 4 | 0.913709 | 5104.75 |
| p.Lys356Arg | 356 | R | -0.0860746 | 0.65443674 | 2.81799431 | 1.50271409 | 1.222267638 | 4 | 1.553062 | 14889.25 |
| p.Lys356Tyr | 356 | Y | 0.28871572 | -0.0072225 | 3.13065636 | 1.98531399 | 1.349365887 | 4 | 2.180912 | 6364.75 |
| p.Lys356Gly | 356 | G | 0.70803622 | 0.67898661 | 2.87821152 | 1.73490991 | 1.500036065 | 4 | 1.085306 | 16740.5 |
| p.Lys356His | 356 | H | 0.45902751 | 0.71401494 | 2.45802067 | 2.70550288 | 1.584141501 | 4 | 1.348039 | 11591 |
| p.Lys356Gln | 356 | Q | 0.41663124 | 1.4922805 | 2.87552642 | 1.75980218 | 1.636060084 | 4 | 1.019756 | 10599.5 |
| p.Lys356Thr | 356 | T | 0.11482247 | 0.30532046 | 3.21551724 | 2.99884105 | 1.658625307 | 4 | 2.811617 | 27343 |
| p.Lys356Trp | 356 | W | 0.63582888 | 0.27801529 | 2.29726604 | 3.60510015 | 1.704052589 | 4 | 2.380189 | 5710.5 |
| p.Lys356Ser | 356 | S | 1.17016764 | 0.9826588 | 2.84678201 | 1.96543994 | 1.741262097 | 4 | 0.724686 | 17579 |
| p.Lys356Ala | 356 | A | 0.8656268 | 1.20804139 | 3.19686554 | 3.93014595 | 2.300169921 | 4 | 2.237181 | 6893 |
| p.Gln361Lys | 361 | K | 0.11023556 | 0.98369819 | 1.00898878 | 2.52679163 | 1.157428541 | 4 | 1.007995 | 3040 |
| p.Arg363Trp | 363 | W | 0.69170931 | 0.19902893 | 2.67903083 | 2.7589919 | 1.582190243 | 4 | 1.764671 | 5487.5 |
| p.Arg368Phe | 368 | F | 0.62792434 | 0.60979435 | 1.74631277 | 0.42331867 | 0.851837533 | 4 | 0.364145 | 10457.5 |
| p.Arg368Ser | 368 | S | 0.03242533 | 0.30713317 | 2.73965134 | 1.71927775 | 1.199621899 | 4 | 1.600206 | 31156 |

| | | | | | | | | | | |
|---|---|---|---|---|---|---|---|---|---|---|
| p.Arg368Glu | 368 | E | -0.1392886 | 0.10743859 | 2.62549499 | 2.26579184 | 1.214859201 | 4 | 2.051483 | 31197 |
| p.Arg368Tyr | 368 | Y | 0.59763132 | 0.63241807 | 2.25107777 | 1.42019857 | 1.225331431 | 4 | 0.611894 | 19382.5 |
| p.Arg368Ile | 368 | I | 0.67287076 | 0.65578149 | 2.72842049 | 0.88223282 | 1.234826387 | 4 | 1.002078 | 19108.25 |
| p.Arg368Leu | 368 | L | 0.13421622 | 0.78934642 | 2.76859362 | 1.27693591 | 1.242273041 | 4 | 1.254596 | 47457.5 |
| p.Arg368Ala | 368 | A | 0.50352594 | 0.63049972 | 3.16758486 | 0.69376286 | 1.248843346 | 4 | 1.64251 | 39707.5 |
| p.Arg368His | 368 | H | 0.61129379 | 1.39297709 | 2.72603603 | 0.43218834 | 1.290623811 | 4 | 1.089762 | 12816.75 |
| p.Arg368Pro | 368 | P | 0.34289439 | 0.3077063 | 2.79106984 | 1.77410633 | 1.303944216 | 4 | 1.449567 | 44250.5 |
| p.Arg368Thr | 368 | T | 0.32248826 | 0.67270745 | 2.48047195 | 1.90749285 | 1.345790125 | 4 | 1.0344 | 47196.25 |
| p.Arg368Val | 368 | V | 0.40693967 | 0.36078291 | 2.51092918 | 2.24333726 | 1.380497253 | 4 | 1.336667 | 56207 |
| p.Arg368Cys | 368 | C | 0.46970506 | 1.02407608 | 2.83806479 | 1.25550073 | 1.396836664 | 4 | 1.031879 | 10054.25 |
| p.Arg368Met | 368 | M | 0.64383592 | 0.38628146 | 2.71962307 | 2.15772488 | 1.476866333 | 4 | 1.297109 | 38599 |
| p.Arg368Asp | 368 | D | -0.0791795 | 0.662196 | 2.27571169 | 3.05808974 | 1.479204475 | 4 | 2.074455 | 24989.75 |
| p.Arg368Gly | 368 | G | 0.43663252 | 0.55136731 | 2.64358859 | 2.41299174 | 1.511145039 | 4 | 1.390502 | 80149.75 |
| p.Arg368Lys | 368 | K | 0.28157668 | 0.6276489 | 3.15272426 | 2.0498846 | 1.527958612 | 4 | 1.758765 | 13455.25 |
| p.Arg368Trp | 368 | W | 0.83186295 | 0.24992156 | 2.41622344 | 2.77328201 | 1.567822489 | 4 | 1.483805 | 12696.75 |
| p.Arg368Asn | 368 | N | 0.20978554 | 1.21458654 | 2.56355547 | 2.71212152 | 1.675012267 | 4 | 1.407995 | 10702.5 |
| p.Arg368Gln | 368 | Q | 0.25444474 | 0.64888162 | 3.01108918 | 2.99727536 | 1.727922727 | 4 | 2.197746 | 9592 |
| p.Arg369Phe | 369 | F | -0.6482318 | -1.4663911 | 2.06377591 | 0.18601199 | 0.033791265 | 4 | 2.28657 | 6535.75 |
| p.Arg369Leu | 369 | L | -0.2058231 | -0.6957739 | 1.67807191 | 0.96437609 | 0.435212737 | 4 | 1.17159 | 1445.75 |
| p.Arg369Met | 369 | M | 0.8322909 | 0.06492412 | 2.79881319 | 0.57291294 | 1.067235284 | 4 | 1.434181 | 9892 |
| p.Arg369Glu | 369 | E | 0.55252019 | 0.43839713 | 2.34952146 | 0.95727913 | 1.074429477 | 4 | 0.77217 | 10244 |
| p.Arg369Gln | 369 | Q | 0.63516331 | 0.47353315 | 2.187627 | 1.53051472 | 1.206709546 | 4 | 0.643754 | 1670.5 |
| p.Arg369Asp | 369 | D | 0.42387938 | -0.1421372 | 2.44890095 | 2.35198533 | 1.27065712 | 4 | 1.75685 | 5960.25 |
| p.Arg369Val | 369 | V | 0.4469724 | 0.44256018 | 1.84165098 | 2.68548938 | 1.354168233 | 4 | 1.221363 | 29165.5 |
| p.Arg369Asn | 369 | N | 0.09915808 | 0.07577045 | 3.52083216 | 1.85687506 | 1.388158937 | 4 | 2.717292 | 5996.25 |
| p.Arg369Gly | 369 | G | 0.83270525 | 0.58928136 | 2.78199935 | 1.83650127 | 1.510121808 | 4 | 1.010345 | 16196 |
| p.Arg369Ala | 369 | A | -0.0753904 | 1.08916193 | 2.67318968 | 2.9946862 | 1.670411844 | 4 | 2.048314 | 23409.25 |
| p.Arg369Lys | 369 | K | 0.69131821 | 0.34922356 | 2.91501848 | 2.989625 | 1.736296312 | 4 | 1.992056 | 7138.25 |
| p.Arg369His | 369 | H | 1.45343361 | 0.68390435 | 3.34948207 | 2.74269696 | 2.057379248 | 4 | 1.463458 | 16719.75 |
| p.Asp370Trp | 370 | W | 1.11005355 | 1.41815807 | 1.45720695 | -1.627088 | 0.589582648 | 4 | 2.207943 | 2820.25 |
| p.Asp370Met | 370 | M | -0.8170701 | 1.33682299 | 1.61996895 | 1.26939769 | 0.852279888 | 4 | 1.261615 | 10874.75 |
| p.Asp370Thr | 370 | T | -0.1214523 | 0.23034579 | 1.48542683 | 1.82629649 | 0.855154208 | 4 | 0.894836 | 3751.5 |
| p.Asp370Lys | 370 | K | 0.44360665 | 0.54343639 | 2.82710588 | 0.39385695 | 1.052001468 | 4 | 1.404311 | 5712.25 |
| p.Asp370Gly | 370 | G | 0.94049899 | -0.1362296 | 1.19555081 | 2.20977034 | 1.052397628 | 4 | 0.928453 | 2306 |
| p.Asp370Arg | 370 | R | 0.17272752 | 0.81595226 | 2.15882729 | 1.11478158 | 1.065572163 | 4 | 0.685703 | 11901.25 |

| | | | | | | | | | | | |
|---|---|---|---|---|---|---|---|---|---|---|---|
| p.Asp370Val | 370 | V | 0.50543384 | 0.54455101 | 2.49642583 | 0.75857941 | 1.076247521 | | 4 | 0.908783 | 5678 |
| p.Asp370Ile | 370 | I | 0.70114461 | 0.94260269 | 2.55019708 | 3.65068668 | 1.961157766 | | 4 | 1.942188 | 5169.25 |
| p.Arg373Ser | 373 | S | 1.01602202 | 0.1162731 | 2.46795989 | 0.53689249 | 1.034286875 | | 4 | 1.048634 | 10304.75 |
| p.Arg373Glu | 373 | E | 0.13996441 | -0.1681001 | 2.85066609 | 1.99184148 | 1.203592966 | | 4 | 2.115678 | 8745.25 |
| p.Arg373Met | 373 | M | 0.23178862 | 0.09280392 | 1.80041331 | 2.73474636 | 1.214938054 | | 4 | 1.626122 | 30686.5 |
| p.Arg373Gly | 373 | G | 0.67678754 | 0.15048294 | 2.40639261 | 1.64613662 | 1.219949927 | | 4 | 1.009355 | 26280.5 |
| p.Arg373Thr | 373 | T | 0.40178056 | 0.45646148 | 3.09505885 | 1.47030189 | 1.355900693 | | 4 | 1.585698 | 24997.75 |
| p.Arg373Lys | 373 | K | 0.49584489 | 0.8943134 | 2.51862966 | 1.62699693 | 1.38394622 | | 4 | 0.791681 | 19431.25 |
| p.Arg373Val | 373 | V | 0.55630374 | 0.50132989 | 3.24727838 | 1.9531023 | 1.564503576 | | 4 | 1.709848 | 11326 |
| p.Arg373Gln | 373 | Q | 0.86766023 | 0.4385312 | 3.15754128 | 2.04629365 | 1.627506591 | | 4 | 1.502473 | 2122.75 |
| p.Arg373Asn | 373 | N | 0.72450391 | 0.96633378 | 1.88797338 | 3.14607826 | 1.681222332 | | 4 | 1.204975 | 7974.25 |
| p.Arg373Leu | 373 | L | 1.21182483 | 0.70927143 | 2.47508488 | 2.75716916 | 1.788337575 | | 4 | 0.969002 | 6802.75 |
| p.Arg373Asp | 373 | D | 0.79316335 | 0.92154384 | 3.52808148 | 2.39626029 | 1.909762241 | | 4 | 1.693002 | 8224.75 |
| p.Arg373Pro | 373 | P | 0.86640337 | 1.17796232 | 3.73260605 | 3.02414235 | 2.200278523 | | 4 | 1.950378 | 10111.25 |
| p.Tyr376Lys | 376 | K | -0.2347454 | -0.0432484 | 2.25986713 | 1 | 0.74546835 | | 4 | 1.313694 | 11126.25 |
| p.Tyr376Val | 376 | V | -0.1245905 | 0.0841334 | 1.85022673 | 1.72472324 | 0.88362321 | | 4 | 1.09915 | 24084.5 |
| p.Tyr376Gly | 376 | G | 0.39373659 | 0.69204645 | 1.99848377 | 0.61134251 | 0.92390233 | | 4 | 0.529084 | 17715.75 |
| p.Tyr376His | 376 | H | 0.73991233 | -0.2646824 | 2.15405728 | 1.17945784 | 0.952186257 | | 4 | 1.007325 | 6629.5 |
| p.Tyr376Asn | 376 | N | 0.27516408 | 0.01072419 | 2.39336323 | 1.619616 | 1.074716876 | | 4 | 1.269036 | 19887 |
| p.Tyr376Glu | 376 | E | -0.1500453 | -0.0210152 | 2.52699243 | 2.10295862 | 1.114722628 | | 4 | 1.953552 | 21038.25 |
| p.Tyr376Pro | 376 | P | 0.66275396 | 0.50733435 | 2.86668427 | 0.44824416 | 1.121254186 | | 4 | 1.362196 | 32655 |
| p.Tyr376Ile | 376 | I | -0.4674328 | 0.05577977 | 3.28876123 | 1.78092488 | 1.164508269 | | 4 | 2.92831 | 6975.5 |
| p.Tyr376Arg | 376 | R | 1.00754526 | 1.16502525 | 2.10080064 | 0.92080783 | 1.298544746 | | 4 | 0.296269 | 7581.5 |
| p.Tyr376Leu | 376 | L | -0.1134246 | 0.91769701 | 2.97702451 | 1.47470042 | 1.313999345 | | 4 | 1.662024 | 17578.25 |
| p.Tyr376Ser | 376 | S | 0.75382351 | 0.01882118 | 2.64842892 | 2.01589137 | 1.359241245 | | 4 | 1.418817 | 7474.75 |
| p.Tyr376Thr | 376 | T | 0.22052606 | 1.25802713 | 2.66478258 | 1.58653028 | 1.432466514 | | 4 | 1.013856 | 36139.75 |
| p.Tyr376Met | 376 | M | 0.55531573 | 0.66831341 | 2.61033906 | 2.59593565 | 1.607475965 | | 4 | 1.323951 | 26925 |
| p.Tyr376Trp | 376 | W | 1.21681139 | 0.09515723 | 3.4484605 | 1.88827492 | 1.66217601 | | 4 | 1.965277 | 1049.25 |
| p.Tyr376Ala | 376 | A | 0.36316587 | 0.65660998 | 3.04935134 | 2.80960255 | 1.719682432 | | 4 | 1.975402 | 27555.25 |
| p.Tyr376Asp | 376 | D | 0.44413403 | 1.01098631 | 2.9229891 | 3.0679206 | 1.861507509 | | 4 | 1.771503 | 12790.5 |
| p.Tyr376Cys | 376 | C | 2.77529371 | -0.5680887 | 2.45943162 | 3.24792751 | 1.978641041 | | 4 | 2.987578 | 677 |
| p.Tyr376Gln | 376 | Q | 0.9068906 | 3.44625623 | 1.79518021 | 3.57568469 | 2.43100293 | | 4 | 1.689408 | 2266 |
| p.Arg380Pro | 380 | P | 0.44804461 | -0.5404526 | 1.20353339 | 2.45169597 | 0.890705335 | | 4 | 1.592905 | 2680.5 |
| p.Arg380Asp | 380 | D | 0.47704716 | 1.17261337 | 2.23551334 | -0.2479275 | 0.909311591 | | 4 | 1.118065 | 4682.5 |
| p.Arg380Ile | 380 | I | 0.54133557 | -0.7382589 | 2.84130225 | 2.53380841 | 1.294546831 | | 4 | 2.875949 | 3723 |

| | | | | | | | | | | |
|---|---|---|---|---|---|---|---|---|---|---|
| p.Arg380Ser | 380 | S | 0.39729818 | -0.3061324 | 2.01088832 | 3.48456782 | 1.396655488 | 4 | 2.878287 | 4235.75 |
| p.Arg380Asn | 380 | N | -0.0058926 | 0.56385347 | 2.63807384 | 3.09953567 | 1.573892604 | 4 | 2.325323 | 2826.5 |
| p.Arg380Gly | 380 | G | 0.80535257 | 2.02469583 | 1.75002175 | 2.77173101 | 1.837950289 | 4 | 0.66027 | 2658 |
| p.Arg380Val | 380 | V | 1.56061264 | 2.49114048 | 3 | 3.08423654 | 2.533997414 | 4 | 0.489745 | 2376.5 |
| p.Leu383Pro | 383 | P | 1.27270479 | 0.68929916 | 2.8708501 | 1.37745766 | 1.552577928 | 4 | 0.864029 | 5589 |
| p.Leu404Ile | 404 | I | 2.23963397 | 0.03380694 | 1.27085391 | 2.13707485 | 1.42034242 | 4 | 1.043257 | 2249 |
| p.Thr405Ile | 405 | I | 0.38862146 | 0.25802753 | 2.16369218 | 0.78060342 | 0.897736145 | 4 | 0.761597 | 29116.5 |
| p.Thr405Gln | 405 | Q | 0.42941252 | -1.0759489 | 1.9648962 | 2.41561911 | 0.933494745 | 4 | 2.517481 | 3446 |
| p.Thr405Ser | 405 | S | 0.38961334 | -0.0517761 | 2.69172835 | 0.7306449 | 0.940052617 | 4 | 1.466309 | 12588.5 |
| p.Thr405Asn | 405 | N | -0.2179362 | 0.33861 | 1.73258715 | 1.93227895 | 0.946384972 | 4 | 1.105045 | 9209 |
| p.Thr405His | 405 | H | -0.3907216 | 1.0682296 | 2.21529864 | 1.30005121 | 1.048214476 | 4 | 1.165482 | 12266 |
| p.Thr405Val | 405 | V | 0.60814775 | 0.51730555 | 2.1469986 | 1.07375304 | 1.086551237 | 4 | 0.559208 | 14719 |
| p.Thr405Gly | 405 | G | 0.01179029 | 0.81163239 | 2.39789846 | 1.4282325 | 1.162388412 | 4 | 1.014688 | 23317.25 |
| p.Thr405Cys | 405 | C | 0.57480263 | 0.37958866 | 2.72280753 | 1.10737197 | 1.1961427 | 4 | 1.13047 | 15700.75 |
| p.Thr405Tyr | 405 | Y | 0.72327166 | 0.54518627 | 2.24406913 | 1.99907133 | 1.377899596 | 4 | 0.752684 | 17900.75 |
| p.Thr405Arg | 405 | R | 0.34124647 | 0.48887706 | 2.87588959 | 2.2184133 | 1.481106607 | 4 | 1.590947 | 24975.25 |
| p.Thr405Pro | 405 | P | -0.0937113 | 0.61278076 | 2.83116886 | 2.86283542 | 1.553268426 | 4 | 2.315018 | 21851.5 |
| p.Thr405Leu | 405 | L | 0.42237041 | 0.38769977 | 2.34757035 | 3.33166035 | 1.622325218 | 4 | 2.137333 | 45136.25 |
| p.Thr405Asp | 405 | D | 0.96879004 | 0.89846197 | 3.10852446 | 1.70525673 | 1.670258301 | 4 | 1.052521 | 1329.75 |
| p.Thr405Met | 405 | M | 1.36015186 | 0.35813288 | 2.94100997 | 2.91928399 | 1.894644675 | 4 | 1.597106 | 12536 |
| p.Ser411Met | 411 | M | 0.14615684 | -0.1679245 | 1.99460674 | 1.17563642 | 0.787118865 | 4 | 0.977304 | 16742.75 |
| p.Ser411Ala | 411 | A | 0.41431561 | 0.45128407 | 2.78124208 | 0.19909359 | 0.961483836 | 4 | 1.484152 | 9034.75 |
| p.Ser411His | 411 | H | -0.705039 | 1.72646746 | 1.95419631 | 1.7328612 | 1.177121503 | 4 | 1.585667 | 5700 |
| p.Ser411Asn | 411 | N | 0.92354184 | 0.62829891 | 1.72662365 | 2.01056924 | 1.322258409 | 4 | 0.425946 | 3421.75 |
| p.Ser411Val | 411 | V | 0.62334515 | 1.0182756 | 3.04209133 | 0.78300457 | 1.366679164 | 4 | 1.273871 | 18468.5 |
| p.Ser411Arg | 411 | R | 0.14264606 | 0.37618503 | 2.61711302 | 2.34179603 | 1.369435038 | 4 | 1.664581 | 24530.25 |
| p.Ser411Tyr | 411 | Y | 1.05629189 | 0.95761269 | 1.76671294 | 1.76024642 | 1.385215985 | 4 | 0.192408 | 3932 |
| p.Ser411Pro | 411 | P | 0.78494349 | 0.72056783 | 2.30676829 | 1.92188533 | 1.433541236 | 4 | 0.643339 | 19324.75 |
| p.Ser411Gln | 411 | Q | 0.57436355 | 0.86046441 | 2.3079632 | 2.400102 | 1.53572329 | 4 | 0.907897 | 25336.5 |
| p.Ser411Gly | 411 | G | 0.41615682 | 0.67498536 | 3.31816618 | 2.13873449 | 1.637010713 | 4 | 1.831329 | 11929.25 |
| p.Ser411Leu | 411 | L | 0.39599174 | 0.70423765 | 2.72576173 | 2.73190449 | 1.639473903 | 4 | 1.598114 | 23739 |
| p.Ser411Glu | 411 | E | 0.36156823 | 0.50900102 | 3.2410081 | 2.69364835 | 1.701306423 | 4 | 2.190638 | 12137.25 |
| p.Ser411Thr | 411 | T | 1.15508134 | 0.9657052 | 2.83805172 | 4.25937351 | 2.304552944 | 4 | 2.406581 | 17869.5 |
| p.Ser411Phe | 411 | F | 2.41037612 | 1.33893913 | 1.67349918 | 5.1300189 | 2.638208331 | 4 | 2.959931 | 4135.75 |
| p.Met413Pro | 413 | P | 0.44929057 | -0.4903004 | 1.81323149 | 0.07416602 | 0.461596907 | 4 | 0.961093 | 4610.75 |

| | | | | | | | | | | | |
|---|---|---|---|---|---|---|---|---|---|---|---|
| p.Met413Gln | 413 | Q | 0.04540194 | 0.03978705 | 2.53520947 | 0.71404245 | 0.833610227 | | 4 | 1.387055 | 6895.25 |
| p.Met413Lys | 413 | K | 0.59448527 | 0.2821763 | 1.17218098 | 2.79986816 | 1.212177678 | | 4 | 1.256269 | 2947.25 |
| p.Met413Ala | 413 | A | 0.77514634 | 0.61013823 | 1.33498425 | 3.23022551 | 1.487623582 | | 4 | 1.445855 | 4684.75 |
| p.Met413Gly | 413 | G | -0.4602333 | 1.74598475 | 2.54326321 | 2.34076404 | 1.542444665 | | 4 | 1.897033 | 10497.25 |
| p.Met413Ile | 413 | I | 0.09644187 | 0.5482563 | 2.6799109 | 2.93441166 | 1.564755184 | | 4 | 2.102915 | 3394.75 |
| p.Met413Leu | 413 | L | 1.2066867 | 0.25340069 | 3.26394491 | 1.96019899 | 1.671057822 | | 4 | 1.615428 | 8822.5 |
| p.Tyr415Ala | 415 | A | -0.7403266 | 0.00952277 | 1.80397228 | 0.40765797 | 0.370206616 | | 4 | 1.140154 | 1628.5 |
| p.Tyr415Arg | 415 | R | -0.2318528 | 1.02876558 | 0.93859946 | 0.64458648 | 0.595024693 | | 4 | 0.330786 | 3830.5 |
| p.Tyr415Pro | 415 | P | -0.8542731 | 0.43922087 | 1.01921792 | 2.1402476 | 0.686103316 | | 4 | 1.55307 | 7601.5 |
| p.Tyr415Thr | 415 | T | 0.54410737 | -0.1828641 | 1.55458885 | 1.01329682 | 0.732282246 | | 4 | 0.542687 | 3462 |
| p.Tyr415Ile | 415 | I | 0.86833342 | 1.00943558 | 3.52479135 | -0.3032467 | 1.274828418 | | 4 | 2.596109 | 2352 |
| p.Tyr415His | 415 | H | 0.36369062 | 0.08947039 | 2.87731748 | 2.33571191 | 1.4165476 | | 4 | 1.949451 | 3698.25 |
| p.Tyr415Gln | 415 | Q | 0.67438417 | 0.82689127 | 2.67829508 | 3.31141342 | 1.872745984 | | 4 | 1.749519 | 16742.25 |
| p.Tyr415Val | 415 | V | 0.50455148 | 1.04292632 | 3.04867385 | 3.98841203 | 2.14614092 | | 4 | 2.706809 | 6290 |
| p.Tyr415Ser | 415 | S | 2.25153877 | 1.44360665 | 3.5187288 | 4.39231742 | 2.901547911 | | 4 | 1.717137 | 1008.75 |
| p.Glu417Thr | 417 | T | -0.1152615 | 0.49321654 | 1.18234252 | -0.0235967 | 0.384175202 | | 4 | 0.354892 | 19601.25 |
| p.Glu417Val | 417 | V | 0.51526786 | 0.74310382 | 0.50392593 | 1.52810016 | 0.822599444 | | 4 | 0.233352 | 9405.5 |
| p.Glu417Gln | 417 | Q | 0.42510432 | 0.07644472 | 2.35295499 | 0.73593011 | 0.897608533 | | 4 | 1.013914 | 9007.5 |
| p.Glu417Cys | 417 | C | -0.9920687 | 1.35124997 | 1.70189283 | 2.61978787 | 1.170215483 | | 4 | 2.364062 | 3452.75 |
| p.Glu417Asp | 417 | D | 0.25803543 | -0.0517814 | 3.05952081 | 1.46723371 | 1.183252133 | | 4 | 1.994121 | 14924 |
| p.Glu417Ser | 417 | S | 0.8647832 | -0.7660869 | 3.29930979 | 1.39844099 | 1.19911177 | | 4 | 2.808115 | 6774.75 |
| p.Glu417Ala | 417 | A | 1.14582044 | 0.73194886 | 2.60827567 | 0.88252716 | 1.342143033 | | 4 | 0.741739 | 4472 |
| p.Glu417Gly | 417 | G | 0.43371122 | 0.13664141 | 2.54880284 | 2.3249944 | 1.361037466 | | 4 | 1.56636 | 9744.75 |
| p.Glu417Asn | 417 | N | 0.57535731 | 0.78607652 | 2.03965254 | 2.39415642 | 1.448810695 | | 4 | 0.81497 | 12374.25 |
| p.Glu417Pro | 417 | P | 0.24934169 | 1.18270872 | 1.69446087 | 2.78315951 | 1.477417697 | | 4 | 1.115698 | 8881 |
| p.Glu417Ile | 417 | I | -0.3121891 | 0.50383141 | 2.61218397 | 3.22930355 | 1.508282458 | | 4 | 2.834517 | 2740.5 |
| p.Glu417Arg | 417 | R | 0.76896908 | 0.11090931 | 2.31860774 | 2.83720485 | 1.508922746 | | 4 | 1.640632 | 7801.25 |
| p.Glu417Leu | 417 | L | 0.33884229 | 0.67489323 | 2.7087687 | 2.43837603 | 1.540220063 | | 4 | 1.454763 | 12252.75 |
| p.Glu417Lys | 417 | K | 0.03964908 | 0.63037118 | 2.34872815 | 3.68871043 | 1.676864709 | | 4 | 2.758182 | 2769.75 |
| p.Tyr428Val | 428 | V | -0.3262365 | 0.57858865 | 3.3380314 | 1.63395516 | 1.306084672 | | 4 | 2.476677 | 6055.5 |
| p.Tyr428Leu | 428 | L | 1.30878429 | -0.2439027 | 2.65326851 | 2.22635043 | 1.486125148 | | 4 | 1.644868 | 8840.75 |
| p.Tyr428Thr | 428 | T | 0.96945612 | 0.00127503 | 2.40731959 | 2.98423268 | 1.590570856 | | 4 | 1.840339 | 9165.75 |
| p.Tyr428Met | 428 | M | 1.7617487 | 0.16333618 | 1.01177717 | 3.76553475 | 1.675599198 | | 4 | 2.367617 | 1188.25 |
| p.Tyr428Asn | 428 | N | 1.46014039 | 1.02185943 | 3.78727068 | 2.2953773 | 2.14116195 | | 4 | 1.483362 | 4367.25 |
| p.Tyr430Asp | 430 | D | 0.03507277 | 0.11458393 | 0.10128334 | -0.3703684 | -0.029857104 | | 4 | 0.052742 | 4947.5 |

| | | | | | | | | | | |
|---|---|---|---|---|---|---|---|---|---|---|
| p.Tyr430Pro | 430 | P | 1.05517671 | 0.75261557 | 1.99627691 | -1.2746557 | 0.632353368 | 4 | 1.896738 | 7903.75 |
| p.Tyr430His | 430 | H | 0.86144244 | 0.86304842 | 1.78849589 | 0.35084079 | 0.965956885 | 4 | 0.358817 | 3277.75 |
| p.Tyr430Ser | 430 | S | -0.0632713 | 0.47269807 | 2.07683745 | 1.78059762 | 1.066715464 | 4 | 1.053234 | 13717.25 |
| p.Tyr430Gly | 430 | G | 0.08756303 | 0.36076174 | 1.99087861 | 1.8575953 | 1.07419967 | 4 | 0.978818 | 6703.5 |
| p.Tyr430Lys | 430 | K | 0.44974589 | 0.03107967 | 2.03271558 | 1.89058021 | 1.101030335 | 4 | 1.020131 | 7123 |
| p.Tyr430Thr | 430 | T | -0.03354 | 0.4962159 | 2.6691755 | 2.81060546 | 1.485614222 | 4 | 2.147719 | 4870.75 |
| p.Tyr430Asn | 430 | N | 1.07649814 | 0.12858421 | 2.59523084 | 2.58206261 | 1.595593947 | 4 | 1.464657 | 2890.25 |
| p.Tyr430Leu | 430 | L | 1.53526303 | 0.54461283 | 2.36176836 | 4.55458885 | 2.249058267 | 4 | 2.914271 | 5824.75 |
| p.Ser435Ala | 435 | A | 0.14723126 | 0.26278959 | 2.69317172 | 1.23145901 | 1.083662895 | 4 | 1.3877 | 15391 |
| p.Ser435Pro | 435 | P | -0.1502074 | 0.33767553 | 3.0599378 | 1.09432738 | 1.085433323 | 4 | 1.994899 | 24704.25 |
| p.Ser435Asp | 435 | D | 0.37610711 | -0.343224 | 2.18919503 | 2.45121111 | 1.16832231 | 4 | 1.866787 | 15232.75 |
| p.Ser435Gly | 435 | G | 0.96841224 | 1.11945205 | 1.94928918 | 0.89374818 | 1.232725412 | 4 | 0.237021 | 22127.5 |
| p.Ser435Ile | 435 | I | 0.68413331 | 0.7126747 | 2.20601476 | 2.0593803 | 1.415550769 | 4 | 0.689452 | 28593 |
| p.Ser435Arg | 435 | R | 0.33718716 | 0.15852392 | 2.34157534 | 3.07318606 | 1.477618118 | 4 | 2.11095 | 34764.5 |
| p.Ser435Thr | 435 | T | 0.23504156 | 0.30269015 | 3.2196046 | 2.45304565 | 1.55259549 | 4 | 2.295981 | 15369.25 |
| p.Ser435Met | 435 | M | 0.34303184 | 0.22304573 | 3.07052839 | 2.58038493 | 1.554247724 | 4 | 2.197069 | 21180.75 |
| p.Ser435Gln | 435 | Q | 0.80830775 | 1.21763233 | 1.95364277 | 2.28581978 | 1.566350659 | 4 | 0.454622 | 32592.75 |
| p.Ser435Asn | 435 | N | 0.02737211 | 0.25457283 | 2.7033572 | 3.35141939 | 1.584180383 | 4 | 2.855733 | 4875 |
| p.Ser435Trp | 435 | W | 0.62514058 | 0.04145736 | 3.39464435 | 2.35334919 | 1.603647871 | 4 | 2.389212 | 7033.5 |
| p.Ser435Phe | 435 | F | 0.31661279 | 0.13185339 | 2.50497437 | 3.52024954 | 1.618422522 | 4 | 2.769172 | 19082 |
| p.Ser435Leu | 435 | L | 0.53969146 | 0.71307239 | 2.90303827 | 2.37363239 | 1.632358625 | 4 | 1.401041 | 79933.5 |
| p.Ser435Tyr | 435 | Y | 0.76100778 | 0.4308426 | 2.71400445 | 2.79672402 | 1.675644713 | 4 | 1.573701 | 11546 |
| p.Ser435Val | 435 | V | 0.21485329 | 0.67159683 | 2.90424102 | 2.92184832 | 1.678134863 | 4 | 2.068157 | 12326 |
| p.Ser435Cys | 435 | C | 1.06182663 | 0.55618924 | 3.11855661 | 2.01502821 | 1.687900173 | 4 | 1.275509 | 6598 |
| p.Ser435His | 435 | H | 0.76037462 | 0.56633538 | 3.89343594 | 1.67724062 | 1.72434664 | 4 | 2.3258 | 15451 |
| p.Tyr446Met | 446 | M | 0.38549322 | 1.29886248 | 1.87800948 | -1.2107106 | 0.587913642 | 4 | 1.815273 | 2588 |
| p.Tyr446Ser | 446 | S | -0.3101745 | 0.06519924 | 1.95770828 | 1.50047834 | 0.803302841 | 4 | 1.201111 | 11158.5 |
| p.Tyr446Thr | 446 | T | 0.46408136 | 0.4139245 | 2.31631449 | 0.64205169 | 0.959093013 | 4 | 0.82827 | 16113 |
| p.Tyr446Lys | 446 | K | 0.43230283 | 0.17970172 | 1.6043679 | 2.41025697 | 1.156657355 | 4 | 1.083696 | 10783.25 |
| p.Tyr446Gln | 446 | Q | 0.33422937 | 1.16932525 | 2.76807844 | 0.424678 | 1.174077764 | 4 | 1.269269 | 16571 |
| p.Tyr446Arg | 446 | R | 0.26803917 | 0.20230179 | 2.5667066 | 1.67113322 | 1.177045196 | 4 | 1.317233 | 46458.25 |
| p.Tyr446Val | 446 | V | 0.24894625 | 0.46653518 | 2.96848131 | 1.20867019 | 1.223158234 | 4 | 1.522643 | 15690 |
| p.Tyr446Ile | 446 | I | -0.0578207 | 1.65844717 | 2.84021956 | 0.52230882 | 1.240788725 | 4 | 1.645072 | 5693.25 |
| p.Tyr446Pro | 446 | P | 0.17837498 | 0.4889592 | 2.26121167 | 2.29014161 | 1.304671863 | 4 | 1.27335 | 10696.25 |
| p.Tyr446Leu | 446 | L | 0.18916463 | 0.97830274 | 2.12413971 | 1.94337818 | 1.308746315 | 4 | 0.810093 | 25822.75 |

| | | | | | | | | | | |
|---|---|---|---|---|---|---|---|---|---|---|
| p.Tyr446His | 446 | H | 0.77535864 | -0.3353781 | 3.34872815 | 1.47547335 | 1.316045524 | 4 | 2.392253 | 2952 |
| p.Tyr446Gly | 446 | G | 0.52154903 | 0.08885993 | 3.15146467 | 1.92848823 | 1.422590465 | 4 | 1.945217 | 17084 |
| p.Tyr446Glu | 446 | E | 0.11305579 | 0.70574028 | 2.65400415 | 2.42506786 | 1.474467021 | 4 | 1.579777 | 10439.25 |
| p.Tyr446Ala | 446 | A | 1.06570583 | 1.06580871 | 3.09169983 | 2.40147939 | 1.906173442 | 4 | 1.021133 | 12866.75 |
| p.Tyr446Asp | 446 | D | -0.0468183 | 1.11947016 | 3.17951105 | 3.44723632 | 1.924849804 | 4 | 2.809316 | 8142.25 |
| p.Gln450His | 450 | H | 0.18359994 | 0.56125571 | 2.68650053 | 3.01774555 | 1.612275432 | 4 | 2.091687 | 5111 |
| p.Gln450Val | 450 | V | 1.2144275 | -0.120389 | 2.18442457 | 3.70604787 | 1.746127738 | 4 | 2.599994 | 3642.5 |
| p.Gln450Met | 450 | M | 0.5276953 | 1.94390544 | 4.01056924 | 1.52588327 | 2.002013312 | 4 | 2.145996 | 1845.25 |
| p.Ser455Thr | 455 | T | -0.3307774 | 1.02127907 | 1.98808668 | 0.29180811 | 0.742599111 | 4 | 0.994751 | 10062.75 |
| p.Ser455His | 455 | H | -0.3251584 | 1.2365366 | 2.21501289 | 0.70667646 | 0.958266886 | 4 | 1.122441 | 4733.5 |
| p.Ser455Phe | 455 | F | 0.73249756 | 0.26553892 | 2.61049759 | 0.24130088 | 0.962458739 | 4 | 1.258227 | 6788.75 |
| p.Ser455Pro | 455 | P | 0.45040732 | 0.34985755 | 1.55831566 | 1.99907962 | 1.089415038 | 4 | 0.667545 | 9667 |
| p.Ser455Tyr | 455 | Y | 0.69885347 | 0.52364438 | 2.55179564 | 0.69942123 | 1.118428677 | 4 | 0.919973 | 5092 |
| p.Ser455Ala | 455 | A | 0.11816273 | 0.12726415 | 1.68085793 | 2.64143226 | 1.141929267 | 4 | 1.538866 | 18864.25 |
| p.Ser455Val | 455 | V | -0.0002684 | 0.57612628 | 2.76832977 | 1.76646702 | 1.27766367 | 4 | 1.52876 | 18627.25 |
| p.Ser455Leu | 455 | L | 0.73914812 | 0.68371569 | 2.7749677 | 1.74073516 | 1.484641665 | 4 | 0.975923 | 40151.5 |
| p.Ser455Gly | 455 | G | 0.48263186 | 0.70115227 | 3.13634161 | 1.73455568 | 1.513670354 | 4 | 1.468359 | 28312.75 |
| p.Ser455Gln | 455 | Q | 0.47624076 | 0.63963712 | 2.62418381 | 2.71521889 | 1.613820145 | 4 | 1.492344 | 11099.75 |
| p.Ser455Met | 455 | M | 1.26775912 | 0.47660183 | 2.33120591 | 2.4018402 | 1.619351763 | 4 | 0.849506 | 7091.5 |
| p.Ser455Glu | 455 | E | 0.61754767 | 1.32274065 | 2.76469703 | 2.02345897 | 1.682111081 | 4 | 0.850318 | 13759.5 |
| p.Ser455Arg | 455 | R | 0.78287959 | 0.78669377 | 2.74700225 | 2.64889073 | 1.741366585 | 4 | 1.221667 | 36424.25 |
| p.Ser455Ile | 455 | I | 0.08614365 | 1.18831849 | 3.53881404 | 2.66870929 | 1.870496366 | 4 | 2.35657 | 9519 |
| p.Thr458Ser | 458 | S | 0.54096789 | 0.38884661 | 1.5849625 | -1.0846911 | 0.357521478 | 4 | 1.207074 | 9789.25 |
| p.Thr458Asp | 458 | D | -0.0802936 | 0.3971833 | 2.64483396 | 0.37676748 | 0.834622794 | 4 | 1.504974 | 8900 |
| p.Thr458Pro | 458 | P | 0.30001683 | -0.3867232 | 2.45198864 | 1.59472147 | 0.990000928 | 4 | 1.624847 | 11809 |
| p.Thr458Leu | 458 | L | 0.33398798 | 1.05801189 | 2.84834534 | 0.70583408 | 1.236544825 | 4 | 1.242013 | 50672.25 |
| p.Thr458Gly | 458 | G | 0.47553587 | 0.52984737 | 2.46408965 | 1.50383198 | 1.243326217 | 4 | 0.88556 | 19606.5 |
| p.Thr458His | 458 | H | 0.27155505 | 1.0847192 | 1.78781474 | 2.22958792 | 1.343419227 | 4 | 0.732867 | 10355.75 |
| p.Thr458Lys | 458 | K | 1.38462778 | -0.1340018 | 3.09953567 | 1.25642735 | 1.401647242 | 4 | 1.754141 | 5400 |
| p.Thr458Met | 458 | M | 0.34800589 | 0.63999135 | 2.62931817 | 2.10231276 | 1.42990704 | 4 | 1.228398 | 24872.5 |
| p.Thr458Asn | 458 | N | 0.89451974 | 0.30424418 | 2.1494609 | 2.7857501 | 1.533493729 | 4 | 1.288968 | 16074.75 |
| p.Thr458Val | 458 | V | 0.70651342 | 0.35160478 | 2.76081234 | 2.34326052 | 1.540547763 | 4 | 1.414197 | 22797.25 |
| p.Thr458Trp | 458 | W | 0.70644365 | 0.68252485 | 3.081665 | 1.93000266 | 1.600159037 | 4 | 1.314812 | 7786.75 |
| p.Thr458Glu | 458 | E | 1.2590701 | 0.93491992 | 2.64724678 | 1.90058471 | 1.685455375 | 4 | 0.572144 | 6327.75 |
| p.Thr458Gln | 458 | Q | 0.70815539 | 1.01161836 | 2.96533287 | 2.09731931 | 1.695606482 | 4 | 1.072159 | 15317.75 |

| | | | | | | | | | | |
|---|---|---|---|---|---|---|---|---|---|---|
| p.Thr458Ala | 458 | A | 0.63593077 | 0.79428388 | 3.18356116 | 2.30796642 | 1.730435557 | 4 | 1.506479 | 10802.5 |
| p.Thr458Ile | 458 | I | 0.14866399 | 1.00418942 | 2.78307473 | 3.72408146 | 1.915002398 | 4 | 2.658616 | 14259.5 |
| p.Thr458Arg | 458 | R | 0.79569672 | 0.8469477 | 2.95675441 | 3.18249346 | 1.945473073 | 4 | 1.693884 | 26912.75 |
| p.Ala460Val | 460 | V | -1.4680723 | 1.06534649 | 2.466318 | 1.50330073 | 0.89172324 | 4 | 2.817385 | 2237.25 |
| p.Lys469Pro | 469 | P | -0.1327153 | -0.4106408 | 2.22954898 | 0.87975932 | 0.641488054 | 4 | 1.428359 | 53770.5 |
| p.Lys469Phe | 469 | F | -0.619833 | 0.23425676 | 2.35363695 | 0.82781902 | 0.698969944 | 4 | 1.569908 | 1242.75 |
| p.Lys469Leu | 469 | L | 0.04108136 | 0.43442885 | 2.57648737 | 1.0403635 | 1.023090269 | 4 | 1.241402 | 17029 |
| p.Lys469Ser | 469 | S | 0.25470219 | -0.5060541 | 2.47533801 | 2.04773084 | 1.067929244 | 4 | 2.026524 | 11055.5 |
| p.Lys469Ile | 469 | I | -0.0588937 | 0.5573186 | 2.52356196 | 1.66002545 | 1.170503079 | 4 | 1.319271 | 5889.75 |
| p.Lys469Asp | 469 | D | -0.4331595 | 0.48855387 | 2.65458464 | 2.27850959 | 1.247122146 | 4 | 2.147828 | 11409 |
| p.Lys469Gly | 469 | G | 0.33875162 | 0.45865086 | 2.21154348 | 2.26997413 | 1.319730023 | 4 | 1.134024 | 52020.75 |
| p.Lys469His | 469 | H | 0.55887327 | 0.47167727 | 2.51699358 | 2.0811206 | 1.407166181 | 4 | 1.093557 | 8910.75 |
| p.Lys469Arg | 469 | R | 1.6410777 | 0.2592839 | 2.65679525 | 1.25792858 | 1.453771355 | 4 | 0.982502 | 9146.75 |
| p.Lys469Thr | 469 | T | 0.79830387 | 0.55871034 | 3.11492361 | 1.57808209 | 1.512504978 | 4 | 1.330618 | 42217.75 |
| p.Lys469Ala | 469 | A | 0.17135223 | 0.80387445 | 2.41940798 | 2.78517858 | 1.544953314 | 4 | 1.579602 | 46008.25 |
| p.Lys469Glu | 469 | E | 0.89258391 | 1.04338271 | 1.81312187 | 2.94135401 | 1.672610623 | 4 | 0.877941 | 5197 |
| p.Lys469Val | 469 | V | 0.57611967 | 0.57216655 | 2.86943915 | 3.32135013 | 1.834768872 | 4 | 2.152943 | 41972.5 |
| p.Phe470Leu | 470 | L | 0.77514634 | -0.1119812 | 2.83187724 | 3.17536181 | 1.667601037 | 4 | 2.530757 | 2879.75 |
| p.Ser474Pro | 474 | P | -0.3600836 | -0.4860879 | 1.98859524 | 0.56679183 | 0.427303885 | 4 | 1.303784 | 22965.25 |
| p.Ser474Gln | 474 | Q | 0.03759679 | 0.98817171 | 2.45094515 | 1.62343665 | 1.275037575 | 4 | 1.039231 | 4952 |
| p.Ser474His | 474 | H | 0.57835977 | 0.97415298 | 2.68222263 | 1.01096186 | 1.311424308 | 4 | 0.8735 | 11284.5 |
| p.Ser474Glu | 474 | E | 0.82960499 | 1.02545624 | 2.50542374 | 1.1786959 | 1.384795217 | 4 | 0.578549 | 12982.25 |
| p.Ser474Leu | 474 | L | -0.148026 | 1.40001831 | 2.94020692 | 1.5800506 | 1.443062459 | 4 | 1.597874 | 23408 |
| p.Ser474Tyr | 474 | Y | 0.46168759 | -0.1339651 | 3.01961102 | 2.6778186 | 1.506288029 | 4 | 2.481417 | 6260.25 |
| p.Ser474Arg | 474 | R | 0.36088326 | 0.85907912 | 3.10074144 | 1.93903 | 1.564933455 | 4 | 1.482207 | 28017 |
| p.Ser474Gly | 474 | G | 0.73784999 | 0.6668737 | 2.72838175 | 2.1516034 | 1.57117721 | 4 | 1.062739 | 16847.25 |
| p.Ser474Trp | 474 | W | 0.99222265 | 0.91093891 | 2.77649396 | 2.15745752 | 1.70927826 | 4 | 0.830443 | 9109.25 |
| p.Ser474Thr | 474 | T | 0.65549645 | 1.48139087 | 2.65721457 | 2.29390345 | 1.772001337 | 4 | 0.795674 | 10321.25 |
| p.Ser474Ile | 474 | I | 1.46678404 | 0.48360798 | 2.9429189 | 3.03268862 | 1.981499887 | 4 | 1.512646 | 13398.25 |
| p.Ser474Val | 474 | V | -0.0648588 | 1.55259844 | 3.88219839 | 2.68650053 | 2.014109636 | 4 | 2.825656 | 7260.5 |
| p.Ser474Met | 474 | M | 0.92329976 | 1.22057007 | 2.01435529 | 4.34250897 | 2.125183523 | 4 | 2.397222 | 8738.75 |
| p.Tyr477Pro | 477 | P | 0.42449783 | -0.0584674 | 1.3708377 | 1.48843843 | 0.806326633 | 4 | 0.559204 | 6872 |
| p.Tyr477Thr | 477 | T | -0.0382314 | 0.67927969 | 3.11769504 | 0.83596184 | 1.148676298 | 4 | 1.867969 | 2581.25 |
| p.Tyr477Ala | 477 | A | 0.39915198 | 0.09102006 | 3.15121881 | 1.95226739 | 1.398414557 | 4 | 2.028961 | 5027.5 |
| p.Thr490Pro | 490 | P | -0.2139179 | -0.3320578 | 2.32081361 | 0.65735616 | 0.60804853 | 4 | 1.498475 | 14511.75 |

| | | | | | | | | | | |
|---|---|---|---|---|---|---|---|---|---|---|
| p.Thr490Ala | 490 | A | 0.2516146 | 0.65436655 | 1.20461772 | 0.48946799 | 0.650016714 | 4 | 0.164034 | 6120.25 |
| p.Thr490Asn | 490 | N | 1.33498425 | 0.65829424 | 0.39366385 | 0.6520767 | 0.759754758 | 4 | 0.162267 | 1736.75 |
| p.Thr490Glu | 490 | E | 0.35776562 | 0.78064427 | 1.84052179 | 0.49797068 | 0.869225588 | 4 | 0.450228 | 8291.25 |
| p.Thr490Val | 490 | V | 0.17776197 | 1.27207145 | 2.71490867 | 0.03697524 | 1.050429334 | 4 | 1.536085 | 14746 |
| p.Thr490Met | 490 | M | -0.0510623 | 0.99222909 | 2.88227804 | 0.47950256 | 1.075736851 | 4 | 1.631912 | 12738.25 |
| p.Thr490His | 490 | H | 0.58691341 | -0.1255778 | 3.09580559 | 1.56872621 | 1.281466849 | 4 | 1.945507 | 7398 |
| p.Thr490Tyr | 490 | Y | 0.1133818 | 0.54646331 | 2.93859946 | 2.14153817 | 1.434995683 | 4 | 1.765393 | 3309.5 |
| p.Thr490Lys | 490 | K | 0.19860776 | -0.1113828 | 2.91753784 | 2.92180231 | 1.481641274 | 4 | 2.773254 | 4416.25 |
| p.Thr490Leu | 490 | L | 0.71363067 | 0.46712601 | 3.18120628 | 1.80533952 | 1.54182562 | 4 | 1.532632 | 13040 |
| p.Thr490Arg | 490 | R | 0.19712673 | 1.68145661 | 2.50250034 | 3.53378985 | 1.978718383 | 4 | 1.985009 | 4590.75 |
| p.Thr490Ser | 490 | S | 0.70454412 | 2.31787273 | 3.06683151 | 3.36276846 | 2.363004203 | 4 | 1.415809 | 4844.75 |
| p.Thr492Gln | 492 | Q | -0.3459274 | 0.61080251 | 1.98792717 | 3.35476468 | 1.401891734 | 4 | 2.612615 | 3594.25 |
| p.Thr499Pro | 499 | P | -0.355026 | 0.07780107 | 1.94678221 | 0.14614317 | 0.453925116 | 4 | 1.039741 | 78455.75 |
| p.Thr499His | 499 | H | 0.74100591 | 0.69586594 | 1.67229617 | 0.5492178 | 0.914596454 | 4 | 0.261862 | 24795.25 |
| p.Thr499Asn | 499 | N | -0.3716188 | 1.14573504 | 2.01693986 | 1.4351937 | 1.056562453 | 4 | 1.03778 | 9633.25 |
| p.Thr499Val | 499 | V | 0.88668594 | 0.45500219 | 2.19779818 | 0.69027391 | 1.057440055 | 4 | 0.609105 | 25619.75 |
| p.Thr499Glu | 499 | E | 0.42788438 | 0.53905083 | 2.15225011 | 1.23494479 | 1.088532528 | 4 | 0.630439 | 17928.75 |
| p.Thr499Ala | 499 | A | 0.33207184 | 0.66654806 | 2.42324577 | 1.75655481 | 1.29460512 | 4 | 0.936051 | 42738.25 |
| p.Thr499Met | 499 | M | 0.73177141 | 0.90778273 | 2.58593173 | 1.09953567 | 1.331255385 | 4 | 0.722206 | 29167.5 |
| p.Thr499Tyr | 499 | Y | 0.42851741 | 0.71513228 | 1.64481236 | 2.60298173 | 1.347860945 | 4 | 0.969682 | 4233 |
| p.Thr499Gln | 499 | Q | 0.46010974 | 0.4085464 | 2.99370001 | 1.92735837 | 1.44742863 | 4 | 1.558454 | 35456.25 |
| p.Thr499Arg | 499 | R | 0.55627222 | 0.1845548 | 2.66119809 | 2.43719119 | 1.459804074 | 4 | 1.613755 | 27262.25 |
| p.Thr499Asp | 499 | D | 1.39040913 | -0.1137487 | 2.75316282 | 1.90336753 | 1.483297684 | 4 | 1.449401 | 13328.25 |
| p.Thr499Leu | 499 | L | 0.09073687 | 0.21544991 | 3.15489366 | 2.4769047 | 1.484496285 | 4 | 2.442715 | 44287.75 |
| p.Thr499Gly | 499 | G | 0.59466533 | 0.31291064 | 2.96912306 | 2.11161199 | 1.497077753 | 4 | 1.587057 | 44629.25 |
| p.Thr499Ile | 499 | I | 0.73156768 | 0.16964716 | 2.76438303 | 2.45036754 | 1.52899135 | 4 | 1.619609 | 16431.5 |
| p.Thr499Ser | 499 | S | 0.50315873 | 0.97821131 | 2.58957913 | 2.05454065 | 1.531372457 | 4 | 0.918906 | 21770.75 |
| p.Thr499Lys | 499 | K | 1.38019305 | 1.43730409 | 2.77229378 | 2.74766498 | 2.084363976 | 4 | 0.609253 | 10223 |
| p.Ala500Gln | 500 | Q | -1.3523017 | -0.5693656 | 1.5849625 | 1.43295941 | 0.27406363 | 4 | 2.139311 | 1200.5 |
| p.Ala500Leu | 500 | L | 0.85573622 | -0.9991603 | 1.37151778 | 2.55814219 | 0.946558965 | 4 | 2.190621 | 13110.75 |
| p.Ala500Glu | 500 | E | 0.64780907 | 0.64931291 | 1.34577484 | 2.19626539 | 1.209790552 | 4 | 0.540528 | 2101 |
| p.Ala500Pro | 500 | P | 0.83346081 | 0.08924605 | 2.4937581 | 1.6984717 | 1.278734168 | 4 | 1.088538 | 10842.25 |
| p.Ala500Arg | 500 | R | 0.46886823 | -0.0149884 | 2.67444451 | 2.29762535 | 1.356487428 | 4 | 1.763855 | 18097.25 |
| p.Ala500Lys | 500 | K | 0.42618588 | 0.54541359 | 2.02399095 | 3.08249007 | 1.519520122 | 4 | 1.613876 | 10164 |
| p.Ala500His | 500 | H | -0.0580619 | 0.61883737 | 2.46221944 | 3.1502771 | 1.543317994 | 4 | 2.281927 | 6678.75 |

| | | | | | | | | | | |
|---|---|---|---|---|---|---|---|---|---|---|
| p.Ala500Ser | 500 | S | 0.2806797 | 0.43789411 | 2.89436868 | 2.56738536 | 1.545081964 | 4 | 1.896752 | 11854.5 |
| p.Ala500Thr | 500 | T | 0.14678538 | 1.03451733 | 1.55433088 | 3.84940012 | 1.646258429 | 4 | 2.494977 | 10448.75 |
| p.Ala500Val | 500 | V | 0.97360026 | 0.3211011 | 2.75969101 | 2.86221341 | 1.729151445 | 4 | 1.633102 | 19130.25 |
| p.Ala500Ile | 500 | I | 2.15192417 | 0.75550027 | 2.99548452 | 2.49569516 | 2.099651029 | 4 | 0.922947 | 4791.5 |
| p.Gly503Met | 503 | M | 0.10454569 | -0.4194852 | 2.0789105 | -0.7958041 | 0.242041737 | 4 | 1.635911 | 12080.75 |
| p.Gly503Pro | 503 | P | -0.058469 | -0.4454268 | 1.90520881 | -0.0886758 | 0.328159309 | 4 | 1.136251 | 24820.25 |
| p.Gly503Val | 503 | V | -0.1476784 | -0.1683622 | 1.44347151 | 0.20163386 | 0.332266174 | 4 | 0.577606 | 24531.5 |
| p.Gly503Ile | 503 | I | -0.3081728 | -0.274723 | 1.50826361 | 0.76393264 | 0.42232513 | 4 | 0.77182 | 25522.5 |
| p.Gly503Asp | 503 | D | -0.1253141 | -0.5620318 | 0.43035055 | 2.00164223 | 0.436161725 | 4 | 1.254136 | 10724.25 |
| p.Gly503Arg | 503 | R | -0.4056036 | -0.2719393 | 1.65555008 | 0.93901935 | 0.479256627 | 4 | 0.980774 | 44071 |
| p.Gly503Glu | 503 | E | 0.01820572 | 0.47005008 | 0.67408839 | 1.17182453 | 0.583542181 | 4 | 0.22892 | 23288.5 |
| p.Gly503Lys | 503 | K | -0.1358213 | -0.0956284 | 2.28801546 | 0.76778241 | 0.706087062 | 4 | 1.285954 | 35206.25 |
| p.Gly503Thr | 503 | T | -0.0172588 | 0.22434628 | 1.29956028 | 1.58352197 | 0.772542441 | 4 | 0.619914 | 13829.5 |
| p.Gly503Leu | 503 | L | 0.17319125 | -0.3791463 | 2.76602806 | 0.57148627 | 0.782889821 | 4 | 1.899863 | 16341.75 |
| p.Gly503Ala | 503 | A | 0.54155197 | 0.60817349 | 2.84056584 | 0.51023661 | 1.125131975 | 4 | 1.30954 | 23948.75 |
| p.Gly503Asn | 503 | N | -0.4443575 | 0.26634714 | 2.2621187 | 2.67324414 | 1.189338128 | 4 | 2.291236 | 11601.25 |
| p.Gly503Ser | 503 | S | 0.18355123 | 0.50563395 | 2.71944783 | 2.03133382 | 1.359991706 | 4 | 1.47092 | 17553.75 |
| p.Pro504Gln | 504 | Q | -1.4019611 | 0.11055838 | 2.05889369 | 0.8573956 | 0.406221632 | 4 | 2.097275 | 4331.25 |
| p.Pro504Asn | 504 | N | 0.42915037 | -0.4462289 | 2.23857235 | -0.0524079 | 0.542271481 | 4 | 1.407003 | 9362.5 |
| p.Pro504Ser | 504 | S | 0.53501611 | 0.08352699 | 1.28500153 | 0.83659744 | 0.685035516 | 4 | 0.25575 | 9680.75 |
| p.Pro504Met | 504 | M | -0.1643005 | 0.19477138 | 2.92396411 | 0.38813534 | 0.835642594 | 4 | 1.990651 | 10640.5 |
| p.Pro504Trp | 504 | W | -0.0409364 | 0.80773583 | 2.44057259 | 0.56390089 | 0.942818217 | 4 | 1.124289 | 4775.25 |
| p.Pro504Arg | 504 | R | 0.00226839 | 0.74789197 | 2.60462953 | 0.55848006 | 0.978317488 | 4 | 1.275641 | 11322 |
| p.Pro504His | 504 | H | 0.59282462 | 0.98580168 | 2.93598351 | 0.21021771 | 1.181206879 | 4 | 1.468813 | 5869.75 |
| p.Pro504Phe | 504 | F | 0.28216415 | -0.176184 | 3.17584984 | 1.46994584 | 1.187943959 | 4 | 2.237526 | 6333.75 |
| p.Pro504Val | 504 | V | 0.88003938 | 0.92286395 | 2.35605796 | 0.86877239 | 1.256933418 | 4 | 0.537465 | 26662 |
| p.Pro504Cys | 504 | C | 0.28657637 | 1.12777333 | 3.0896007 | 1.1220766 | 1.406506747 | 4 | 1.415214 | 18673 |
| p.Pro504Thr | 504 | T | 0.79915963 | 0.88848181 | 2.61074426 | 1.6250281 | 1.48085345 | 4 | 0.70435 | 23231.5 |
| p.Pro504Leu | 504 | L | 1.16361636 | 0.88524359 | 2.95463947 | 1.24072688 | 1.561056575 | 4 | 0.886455 | 7851.25 |
| p.Pro504Glu | 504 | E | 0.69823697 | 0.4169547 | 3.15432815 | 2.30651304 | 1.644008215 | 4 | 1.706708 | 5455.5 |
| p.Pro504Asp | 504 | D | 0.7916464 | 0.72412525 | 2.27052894 | 3.05269742 | 1.709749504 | 4 | 1.310784 | 4213.75 |
| p.Pro504Gly | 504 | G | 0.54959116 | 0.86433228 | 2.8505191 | 2.90298892 | 1.791857864 | 4 | 1.586302 | 14326 |
| p.Pro504Lys | 504 | K | 0.61917822 | 0.8675414 | 2.92005505 | 3.22884613 | 1.9089052 | 4 | 1.837501 | 10313.5 |
| p.Pro504Ile | 504 | I | 0.83885224 | 0.82087386 | 2.93128725 | 3.89433274 | 2.121336524 | 4 | 2.378502 | 4014.75 |
| p.Asn505Glu | 505 | E | -0.2695552 | -0.3893198 | 1.35587543 | 1.67456596 | 0.592891591 | 4 | 1.153572 | 8020.5 |

| | | | | | | | | | | |
|---|---|---|---|---|---|---|---|---|---|---|
| p.Asn505Lys | 505 | K | -0.1203967 | -0.1157863 | 1.34464817 | 1.30136941 | 0.602458635 | 4 | 0.692572 | 5655.75 |
| p.Asn505Asp | 505 | D | -0.3986515 | -0.3358302 | 1.00705476 | 2.15884984 | 0.607855711 | 4 | 1.489514 | 3228.75 |
| p.Asn505His | 505 | H | 0.05212712 | 0.28820317 | 1.77294779 | 0.89760841 | 0.752721624 | 4 | 0.589488 | 24829.5 |
| p.Asn505Pro | 505 | P | 0.00449701 | -0.6728723 | 2.52011136 | 1.33493652 | 0.796668155 | 4 | 2.015691 | 17523.5 |
| p.Asn505Arg | 505 | R | -0.046188 | 0.20802332 | 2.48313502 | 0.57639322 | 0.805340883 | 4 | 1.316433 | 19467 |
| p.Asn505Leu | 505 | L | -0.1206858 | 0.09397775 | 2.50250034 | 1.27622208 | 0.938003595 | 4 | 1.465082 | 28228.25 |
| p.Asn505Val | 505 | V | 0.31702335 | 1.28354392 | 1.45437839 | 0.72858888 | 0.945883633 | 4 | 0.271755 | 15716.25 |
| p.Asn505Tyr | 505 | Y | 0.38905371 | 0.72016324 | 2.04138392 | 0.8893142 | 1.009978768 | 4 | 0.515966 | 10596 |
| p.Asn505Gln | 505 | Q | 0.5157147 | 0.59063284 | 2.66429653 | 0.34133665 | 1.027995178 | 4 | 1.2009 | 27281.75 |
| p.Asn505Cys | 505 | C | 0.81126515 | 0.69477037 | 1.44651573 | 2.2707206 | 1.305817963 | 4 | 0.522932 | 4707.25 |
| p.Asn505Thr | 505 | T | 0.98961836 | 0.0051691 | 2.91488339 | 2.11683825 | 1.506627276 | 4 | 1.625739 | 4288 |
| p.Asn505Trp | 505 | W | 0.39684855 | 1.65595433 | 2.86716432 | 2.01655118 | 1.734129595 | 4 | 1.052654 | 2019.5 |
| p.Asn505Ser | 505 | S | 0.69058334 | 1.81969052 | 3.357552 | 1.19826963 | 1.766523873 | 4 | 1.338253 | 2739.5 |
| p.Asn505Ile | 505 | I | 1.4221578 | 0.8785555 | 3.6229904 | 1.8189305 | 1.935658549 | 4 | 1.413955 | 11054 |
| p.Gly506Phe | 506 | F | -0.7024169 | 0.3177115 | 2.18442457 | -1.5915352 | 0.05204601 | 4 | 2.629397 | 3093.25 |
| p.Gly506Lys | 506 | K | 0.61532075 | -1.3061201 | 1.3079889 | -0.1907783 | 0.106602802 | 4 | 1.262115 | 7240 |
| p.Gly506Ser | 506 | S | 0.06118352 | -0.4222982 | 1.25397507 | 1.43716279 | 0.582505797 | 4 | 0.820906 | 15269 |
| p.Gly506Leu | 506 | L | -0.4126451 | 0.36923381 | 1.73102804 | 1.3340819 | 0.755424665 | 4 | 0.933392 | 32577.75 |
| p.Gly506Asn | 506 | N | -0.2416089 | 0.20645088 | 2.15919859 | 1.11547722 | 0.80987944 | 4 | 1.127935 | 2725.5 |
| p.Gly506Val | 506 | V | 0.1621012 | 0.23876294 | 1.97139308 | 1.2077746 | 0.895007957 | 4 | 0.741413 | 12541 |
| p.Gly506Pro | 506 | P | -0.2925017 | 0.09519856 | 2.12254349 | 1.82424703 | 0.937371849 | 4 | 1.471008 | 24010.75 |
| p.Gly506Arg | 506 | R | 0.07335724 | 0.48035264 | 2.14522456 | 1.7242245 | 1.105789736 | 4 | 0.973325 | 19495.25 |
| p.Gly506His | 506 | H | 0.1680332 | 0.33900346 | 2.60510015 | 1.51333957 | 1.156369096 | 4 | 1.290381 | 5856.25 |
| p.Gly506Thr | 506 | T | -0.0864734 | 0.53737304 | 2.78572491 | 1.5849625 | 1.205396761 | 4 | 1.585564 | 3841.5 |
| p.Gly506Glu | 506 | E | 0.67730472 | 0.07364792 | 2.2735837 | 2.22117956 | 1.311428977 | 4 | 1.229201 | 7704.75 |
| p.Gly506Met | 506 | M | -0.3304901 | 0.7868415 | 2.68757757 | 2.9887263 | 1.533163814 | 4 | 2.493845 | 9639 |
| p.Gly506Cys | 506 | C | 1.06183925 | 1.64482088 | 2.10479619 | 3.42626475 | 2.05943027 | 4 | 1.012461 | 2199.75 |
| p.Ser507Pro | 507 | P | 0.00020057 | 0.1276142 | 1.94576609 | 0.29661956 | 0.592550103 | 4 | 0.828604 | 50937 |
| p.Ser507Asp | 507 | D | -0.2026703 | -0.2583543 | 2.63838699 | 1.58001813 | 0.939345129 | 4 | 2.011963 | 15634.5 |
| p.Ser507Trp | 507 | W | 1.64128391 | -0.0423315 | 1.98935276 | 0.56634682 | 1.038662999 | 4 | 0.886198 | 3166 |
| p.Ser507Val | 507 | V | 0.84486468 | 0.65525269 | 1.23883248 | 1.54387681 | 1.070706664 | 4 | 0.158588 | 32839.25 |
| p.Ser507Lys | 507 | K | 0.60638014 | 0.45388479 | 2.2067108 | 1.01707351 | 1.071012309 | 4 | 0.629817 | 32999.25 |
| p.Ser507Glu | 507 | E | 0.17915497 | 0.03060799 | 2.13422839 | 2.08237184 | 1.106590797 | 4 | 1.342021 | 19766.75 |
| p.Ser507Arg | 507 | R | 0.61116757 | -0.0310104 | 2.51523888 | 1.58265656 | 1.169513145 | 4 | 1.244891 | 17760.25 |
| p.Ser507Leu | 507 | L | 0.41798695 | 0.24568258 | 2.74171717 | 1.64630409 | 1.262922695 | 4 | 1.360836 | 37646.25 |

| | | | | | | | | | | |
|---|---|---|---|---|---|---|---|---|---|---|
| p.Ser507Tyr | 507 | Y | 0.55489103 | 1.01440177 | 2.97461468 | 0.51175858 | 1.263916515 | 4 | 1.352402 | 7553 |
| p.Ser507His | 507 | H | -0.0074264 | 0.51007325 | 2.63263606 | 2.11390779 | 1.312297689 | 4 | 1.590369 | 25479.25 |
| p.Ser507Met | 507 | M | -0.0526498 | 0.55962149 | 2.22513448 | 2.93672583 | 1.417207998 | 4 | 1.952539 | 18269.5 |
| p.Ser507Ile | 507 | I | 0.17558287 | 0.91242698 | 2.84670359 | 1.87101356 | 1.45143175 | 4 | 1.34705 | 7607.25 |
| p.Ser507Thr | 507 | T | 0.24613235 | 0.32732091 | 2.3585777 | 2.88711937 | 1.454787582 | 4 | 1.866813 | 35400.25 |
| p.Ser507Gln | 507 | Q | 0.73320557 | 0.40059289 | 2.49610911 | 2.70718593 | 1.584273375 | 4 | 1.405931 | 32891 |
| p.Ser507Ala | 507 | A | 0.95944495 | 0.54400747 | 2.6493236 | 2.51173304 | 1.666127264 | 4 | 1.146759 | 35123 |
| p.Ser507Gly | 507 | G | 0.3731434 | 0.59835749 | 3.55057951 | 2.57114083 | 1.773305307 | 4 | 2.378734 | 24970.5 |
| p.Ser507Asn | 507 | N | 0.26559813 | 1.07103425 | 2.45361664 | 3.66235253 | 1.863150387 | 4 | 2.255133 | 5825.75 |
| p.Gly508Asp | 508 | D | -1.0507746 | -0.2836207 | 1.3016557 | -0.220842 | -0.063395396 | 4 | 0.970524 | 6158.75 |
| p.Gly508Trp | 508 | W | -1.0836212 | -0.9312313 | 1.83834674 | 0.82778682 | 0.162820264 | 4 | 2.000045 | 9546.75 |
| p.Gly508Thr | 508 | T | -0.4775211 | -0.0396595 | 2.11872022 | 0.00838779 | 0.402481855 | 4 | 1.356893 | 17967 |
| p.Gly508His | 508 | H | 0.05412641 | -1.0292307 | 2.46481147 | 0.51568765 | 0.501348707 | 4 | 2.132691 | 8205 |
| p.Gly508Val | 508 | V | 0.10406103 | -0.5590778 | 2.60613652 | 0.15298514 | 0.576026216 | 4 | 1.937175 | 15237.75 |
| p.Gly508Pro | 508 | P | -0.6449553 | -0.2775724 | 2.35478178 | 1.05402101 | 0.621568774 | 4 | 1.86786 | 16264.75 |
| p.Gly508Gln | 508 | Q | -0.4104006 | -0.5044459 | 1.533682 | 1.92477572 | 0.635902795 | 4 | 1.620782 | 12215.5 |
| p.Gly508Leu | 508 | L | 0.03794989 | -0.2860421 | 1.96532444 | 0.91889754 | 0.659032433 | 4 | 1.017613 | 40626 |
| p.Gly508Arg | 508 | R | -0.645514 | -0.205919 | 2.39073361 | 1.25363837 | 0.698234734 | 4 | 1.93206 | 19392.75 |
| p.Gly508Glu | 508 | E | -0.1573935 | -0.2252129 | 2.6169729 | 0.82542649 | 0.764948244 | 4 | 1.754929 | 4661.25 |
| p.Gly508Ala | 508 | A | -0.0798329 | 0.18720833 | 2.69442848 | 1.31038647 | 1.028047587 | 4 | 1.596983 | 15809.5 |
| p.Gly508Phe | 508 | F | 0.0833431 | 1.25728061 | 1.96271898 | 1.2558215 | 1.139791046 | 4 | 0.606853 | 8357.75 |
| p.Gly508Lys | 508 | K | -0.3368745 | 0.02816353 | 2.88566732 | 2.12055892 | 1.174378829 | 4 | 2.47382 | 5023.75 |
| p.Gly508Ser | 508 | S | 0.46862191 | 0.22580408 | 2.37974417 | 2.59253799 | 1.416677039 | 4 | 1.542378 | 30018.5 |
| p.Gly508Tyr | 508 | Y | 0.04107594 | 0.1459667 | 2.78290188 | 3.16448086 | 1.533606347 | 4 | 2.791227 | 1969.75 |
| p.Lys509Tyr | 509 | Y | -0.3806955 | -0.5933676 | 2.33135751 | -1.2565958 | 0.025174648 | 4 | 2.502913 | 9574.25 |
| p.Lys509Pro | 509 | P | -0.6446702 | -0.5787907 | 2.10332725 | 0.44728211 | 0.331787121 | 4 | 1.644771 | 41928.5 |
| p.Lys509Leu | 509 | L | -0.474097 | 0.04508192 | 1.27187177 | 0.70530001 | 0.387039173 | 4 | 0.580903 | 26845.25 |
| p.Lys509Asp | 509 | D | -0.7473261 | -0.2921808 | 1.82919931 | 0.76904557 | 0.389684505 | 4 | 1.324617 | 4862.75 |
| p.Lys509Ser | 509 | S | 0.02578868 | 0.16602582 | 1.66164768 | -0.2498301 | 0.400908018 | 4 | 0.73627 | 29697.5 |
| p.Lys509Glu | 509 | E | 0.08712061 | 0.08307107 | 1.71829638 | 0.12677868 | 0.503816685 | 4 | 0.655927 | 13802.75 |
| p.Lys509Phe | 509 | F | -0.580816 | 0.03765585 | 1.82560429 | 1.12285675 | 0.60132523 | 4 | 1.162012 | 5156.75 |
| p.Lys509Val | 509 | V | -0.6080972 | -0.3732268 | 2.16513199 | 2.08001803 | 0.815956503 | 4 | 2.286737 | 24683.25 |
| p.Lys509Gln | 509 | Q | 0.11519288 | -0.404859 | 3.30455217 | 0.40703771 | 0.855480941 | 4 | 2.778511 | 14263.75 |
| p.Lys509Gly | 509 | G | 0.07416929 | 0.20936878 | 2.69934052 | 0.47996628 | 0.865711218 | 4 | 1.522773 | 17055.5 |
| p.Lys509Ile | 509 | I | 0.85611499 | -0.6695985 | 2.27321989 | 1.08935308 | 0.887272366 | 4 | 1.462168 | 11573 |

| | | | | | | | | | | |
|---|---|---|---|---|---|---|---|---|---|---|
| p.Lys509Met | 509 | M | -0.3610126 | -0.1851436 | 2.87403904 | 1.2973395 | 0.906305565 | 4 | 2.27408 | 21166.75 |
| p.Lys509Thr | 509 | T | -0.0042432 | 0.69223057 | 1.47476578 | 2.18654314 | 1.087324067 | 4 | 0.902004 | 32091 |
| p.Lys509Arg | 509 | R | -0.2293012 | 0.32429173 | 2.33386536 | 1.96626329 | 1.098779785 | 4 | 1.547198 | 21647.75 |
| p.Lys509Ala | 509 | A | 0.07685389 | -0.0734533 | 2.43129478 | 2.10616172 | 1.135214268 | 4 | 1.734523 | 41437.25 |
| p.Lys509Trp | 509 | W | 0.36963347 | -0.2267401 | 2.5987683 | 2.00603639 | 1.186924522 | 4 | 1.776886 | 4249.25 |
| p.Lys509His | 509 | H | 0.53085152 | 0.18912843 | 2.24519358 | 1.9946125 | 1.239946507 | 4 | 1.062359 | 19974 |
| p.Lys509Asn | 509 | N | 0.5849625 | 0.3737202 | 1.42576391 | 3.36797546 | 1.438105518 | 4 | 1.861773 | 3788.5 |
| p.Lys509Cys | 509 | C | -0.241586 | 1.71353538 | 3.422233 | 0.982298 | 1.469120097 | 4 | 2.345967 | 3217.75 |
| p.Ser510Gln | 510 | Q | -0.2679211 | 0.36693006 | 1.86175006 | -1.6675876 | 0.073292876 | 4 | 2.143965 | 7319.25 |
| p.Ser510Glu | 510 | E | -0.0829916 | -0.5121935 | 2.24776765 | -0.0232126 | 0.407342486 | 4 | 1.552839 | 37967.5 |
| p.Ser510Val | 510 | V | -0.2738115 | -0.2433601 | 1.96187948 | 0.20486538 | 0.412393319 | 4 | 1.114955 | 46533 |
| p.Ser510Arg | 510 | R | 0.70765694 | -1.0632351 | 1.20456496 | 0.98748278 | 0.459117398 | 4 | 1.071397 | 26364.75 |
| p.Ser510Tyr | 510 | Y | 0.0305717 | 0.5333869 | 1.27257818 | 0.05907923 | 0.473904001 | 4 | 0.336681 | 3386.75 |
| p.Ser510Ile | 510 | I | -0.3887892 | -0.0556176 | 1.6016799 | 0.8512626 | 0.502133933 | 4 | 0.811908 | 22661.75 |
| p.Ser510Leu | 510 | L | -0.2989338 | -0.3302754 | 2.3230614 | 0.38964631 | 0.520874626 | 4 | 1.55388 | 28223 |
| p.Ser510Ala | 510 | A | 0.03578138 | -0.1053702 | 1.42177233 | 0.88069051 | 0.558218518 | 4 | 0.521001 | 48179.25 |
| p.Ser510Asp | 510 | D | -0.5430564 | -0.9237823 | 2.49827844 | 1.21250329 | 0.560985753 | 4 | 2.533674 | 27883.5 |
| p.Ser510Asn | 510 | N | -0.7525744 | -0.143119 | 1.84488093 | 1.30451104 | 0.563424638 | 4 | 1.474132 | 10703.5 |
| p.Ser510Pro | 510 | P | 0.30735917 | -0.3708881 | 1.83263604 | 0.65216618 | 0.60531832 | 4 | 0.850087 | 43362.5 |
| p.Ser510Phe | 510 | F | 0.54370798 | 1.80240995 | 1 | -0.8881714 | 0.614486631 | 4 | 1.274258 | 5368.75 |
| p.Ser510Cys | 510 | C | 0.15513076 | -0.3960262 | 1.83183503 | 1.50540373 | 0.774085841 | 4 | 1.135309 | 15455.25 |
| p.Ser510Met | 510 | M | 0.04347164 | 0.17860873 | 1.95849645 | 1.0600791 | 0.810163979 | 4 | 0.789268 | 40741.75 |
| p.Ser510Lys | 510 | K | -0.2547309 | 0.01414053 | 2.44038693 | 1.1456476 | 0.836361051 | 4 | 1.511695 | 25576.75 |
| p.Ser510His | 510 | H | 0.45724125 | -0.3101225 | 2.61956116 | 0.88989024 | 0.914142541 | 4 | 1.538875 | 6257.5 |
| p.Ser510Gly | 510 | G | -0.1010489 | 0.45268184 | 2.57027186 | 0.85339017 | 0.943823731 | 4 | 1.32883 | 51805.5 |
| p.Ser510Thr | 510 | T | 0.43119188 | 0.89052014 | 2.8194846 | 1.79866158 | 1.484964549 | 4 | 1.114384 | 63008 |
| p.Thr511Asp | 511 | D | -0.5700253 | -0.5483837 | 2.12567678 | -1.277781 | -0.067628312 | 4 | 2.252862 | 8914.5 |
| p.Thr511Pro | 511 | P | -0.8008293 | -0.0675932 | 1.79002684 | -0.647359 | 0.068561317 | 4 | 1.416788 | 13539.25 |
| p.Thr511Trp | 511 | W | 0.94997828 | -0.3015466 | 1.5849625 | 0.36838741 | 0.650445398 | 4 | 0.649629 | 3204.25 |
| p.Thr511Tyr | 511 | Y | -0.4107544 | 0.15779681 | 2.23562825 | 1.39231742 | 0.843747024 | 4 | 1.427521 | 4429.25 |
| p.Thr511Glu | 511 | E | -0.0418202 | 0.2754894 | 2.26085252 | 0.92702055 | 0.855385575 | 4 | 1.040576 | 8476.25 |
| p.Thr511Val | 511 | V | 0.3139727 | -0.3219281 | 2.20655023 | 1.28350917 | 0.870526001 | 4 | 1.229071 | 23627 |
| p.Thr511Arg | 511 | R | -0.2644759 | -0.0249057 | 2.7604593 | 1.04502043 | 0.879024542 | 4 | 1.897345 | 35132.75 |
| p.Thr511Lys | 511 | K | 0.13022989 | 0.41232311 | 2.24110027 | 1.63582982 | 1.104870771 | 4 | 1.000827 | 16683.25 |
| p.Thr511Met | 511 | M | 0.20049575 | -0.2715238 | 3.19736126 | 1.49702563 | 1.155839708 | 4 | 2.411422 | 24916.5 |

| | | | | | | | | | | |
|---|---|---|---|---|---|---|---|---|---|---|
| p.Thr511Ser | 511 | S | -0.0629267 | 1.11717355 | 2.3479233 | 1.33952215 | 1.185423078 | 4 | 0.979396 | 13715.75 |
| p.Thr511Gly | 511 | G | -0.2209638 | 0.39380822 | 2.46588252 | 2.30116953 | 1.234974124 | 4 | 1.826408 | 12811.25 |
| p.Thr511Gln | 511 | Q | 0.34712828 | 0.61378367 | 2.88081413 | 1.23941852 | 1.270286152 | 4 | 1.292656 | 27297.25 |
| p.Thr511Asn | 511 | N | 0.89265575 | 0.07153051 | 2.1181171 | 2.23866081 | 1.330241045 | 4 | 1.073936 | 12084.5 |
| p.Thr511His | 511 | H | -0.1500177 | 0.57717115 | 3.09895848 | 1.90164493 | 1.356939222 | 4 | 2.070098 | 18666 |
| p.Thr511Ala | 511 | A | 0.24729124 | 0.44168495 | 1.97352779 | 2.97077277 | 1.408319185 | 4 | 1.68103 | 10446.75 |
| p.Thr511Leu | 511 | L | 0.73768751 | 0.61884121 | 2.43717881 | 2.34582548 | 1.534883252 | 4 | 0.98214 | 56656.5 |
| p.Thr511Ile | 511 | I | 0.89572952 | 0.5343998 | 2.07194984 | 3.50589093 | 1.751992523 | 4 | 1.798084 | 2348.5 |
| p.Thr511Cys | 511 | C | 0.83487935 | 0.75739157 | 2.53743413 | 3.26678654 | 1.849122899 | 4 | 1.568037 | 2335.75 |
| p.Tyr520Phe | 520 | F | 0.12297404 | 0.1298035 | 2.19417255 | 0.01559686 | 0.615636736 | 4 | 1.110191 | 2881.5 |
| p.Tyr520Pro | 520 | P | 0.15776236 | -0.163556 | 2.22080617 | 0.42327365 | 0.659571545 | 4 | 1.140881 | 9226.75 |
| p.Tyr520Asp | 520 | D | -0.7920397 | 0.61118897 | 2.38773015 | 0.63251443 | 0.709848455 | 4 | 1.695556 | 5634.75 |
| p.Tyr520Val | 520 | V | -1.2646473 | 1.14868698 | 2.142019 | 1.21044016 | 0.809124721 | 4 | 2.117831 | 10791.75 |
| p.Tyr520Leu | 520 | L | 0.44873702 | -0.3042307 | 2.49381461 | 1.02452232 | 0.915710809 | 4 | 1.402858 | 2692.5 |
| p.Tyr520Gln | 520 | Q | 0.03233065 | 0.82126654 | 1.38109017 | 1.76782656 | 1.00062848 | 4 | 0.567705 | 2054.25 |
| p.Tyr520Thr | 520 | T | 0.14303558 | -0.0899558 | 2.58419897 | 1.86588707 | 1.125791446 | 4 | 1.706182 | 16766.5 |
| p.Tyr520Ala | 520 | A | 0.84623957 | 1.32529607 | 3.1122095 | -0.4845955 | 1.1997874 | 4 | 2.211751 | 3780 |
| p.Tyr520Glu | 520 | E | 1.26711249 | 0.27106473 | 2.13993026 | 2.12905272 | 1.451790051 | 4 | 0.786813 | 5935.25 |
| p.Tyr520Met | 520 | M | 0.07430382 | 1.28606343 | 1.56884284 | 3.59387055 | 1.630770159 | 4 | 2.133003 | 5614.75 |
| p.Tyr520Lys | 520 | K | 0.78708253 | 0.1082163 | 3.02718483 | 2.68528956 | 1.651943303 | 4 | 2.030057 | 3387.75 |
| p.Tyr520Gly | 520 | G | 0.16187615 | 1.74164208 | 3.0052943 | 1.98732087 | 1.72403335 | 4 | 1.383865 | 4589 |
| p.Thr523Glu | 523 | E | 1.11588071 | 1.60554338 | 0.92405115 | 0.57411512 | 1.054897589 | 4 | 0.185067 | 1413.5 |
| p.Thr523Pro | 523 | P | 0.73031389 | 0.69863619 | 2.91194382 | 1.20836554 | 1.387314858 | 4 | 1.087482 | 3737.5 |
| p.Thr523Lys | 523 | K | 0.00910221 | 0.48542683 | 3.24158599 | 1.83953533 | 1.393912587 | 4 | 2.118508 | 2162.25 |
| p.Thr523Leu | 523 | L | 0.59019915 | 0.06039067 | 2.63742992 | 2.98935276 | 1.569343124 | 4 | 2.130966 | 2427.5 |
| p.Thr523Ser | 523 | S | 1.02893589 | 0.25319408 | 2.26946067 | 3.49605685 | 1.761911874 | 4 | 2.026116 | 3279 |
| p.Pro533Gln | 533 | Q | 0.35880503 | 0.73941173 | 1.51488883 | 1.68013114 | 1.073309182 | 4 | 0.395076 | 26606 |
| p.Pro533Ala | 533 | A | 0.17967942 | 0.33833846 | 2.27583351 | 1.72383118 | 1.129420643 | 4 | 1.065135 | 57032.25 |
| p.Pro533Lys | 533 | K | 0.0983252 | 0.86257803 | 1.99674702 | 1.88679992 | 1.211112543 | 4 | 0.811182 | 25829.25 |
| p.Pro533Glu | 533 | E | 0.67478076 | 0.27914803 | 2.70180915 | 1.21660071 | 1.218084664 | 4 | 1.126074 | 35219.25 |
| p.Pro533Met | 533 | M | 0.78197888 | 0.39218424 | 2 | 1.85931689 | 1.258370002 | 4 | 0.629459 | 14677.5 |
| p.Pro533Ser | 533 | S | 0.25705225 | 0.51544859 | 2.12811961 | 2.26786636 | 1.292121702 | 4 | 1.10852 | 28272.5 |
| p.Pro533Thr | 533 | T | 0.27870704 | 0.28578383 | 2.69927283 | 1.96940573 | 1.308292356 | 4 | 1.492489 | 79476.75 |
| p.Pro533Asn | 533 | N | 0.60873795 | 0.54132074 | 2.82110589 | 1.38410564 | 1.338817558 | 4 | 1.122749 | 19623 |
| p.Pro533Val | 533 | V | 0.51306329 | 0.29696616 | 3.09288546 | 1.53985982 | 1.360693681 | 4 | 1.627527 | 70588.25 |

| | | | | | | | | | | | |
|---|---|---|---|---|---|---|---|---|---|---|---|
| p.Pro533Arg | 533 | R | 0.69571284 | 0.77281959 | 2.68367484 | 1.72831053 | 1.470129451 | | 4 | 0.875104 | 57236.75 |
| p.Pro533Asp | 533 | D | 0.83501227 | 0.63083365 | 2.36207524 | 2.40378877 | 1.557927483 | | 4 | 0.914748 | 23402.5 |
| p.Pro533Leu | 533 | L | 0.27008478 | 0.80231 | 2.35429215 | 2.83734966 | 1.56600915 | | 4 | 1.500118 | 36027.25 |
| p.Pro533Trp | 533 | W | 0.86657892 | 0.55801145 | 2.82489722 | 2.20848977 | 1.614494338 | | 4 | 1.16448 | 23114.25 |
| p.Pro533Gly | 533 | G | 0.26518843 | 0.66858693 | 2.96426098 | 2.74565734 | 1.660923421 | | 4 | 1.936048 | 67985.75 |
| p.Pro533His | 533 | H | 0.62612101 | 1.15841942 | 1.44999734 | 3.64927904 | 1.720954201 | | 4 | 1.768986 | 12407 |
| p.Pro533Tyr | 533 | Y | 0.72000443 | 0.62865909 | 3.66479667 | 1.96213784 | 1.743899508 | | 4 | 2.009865 | 10784.5 |
| p.Pro533Cys | 533 | C | 0.48372827 | 0.81694029 | 2.96829114 | 2.93888849 | 1.801962045 | | 4 | 1.786978 | 9564.25 |
| p.Pro533Ile | 533 | I | 0.49852421 | 1.21108867 | 3.49389802 | 2.32326951 | 1.881695104 | | 4 | 1.71902 | 7566.25 |
| p.Tyr537Lys | 537 | K | -0.0711297 | -0.486418 | 1.70564398 | -0.0421063 | 0.276497501 | | 4 | 0.948951 | 39990.5 |
| p.Tyr537Glu | 537 | E | -0.5686082 | -0.3078397 | 1.74631145 | 0.60915808 | 0.369755416 | | 4 | 1.097294 | 68848 |
| p.Tyr537Ser | 537 | S | -0.613097 | -0.1686904 | 1.96307523 | 0.44403048 | 0.406329566 | | 4 | 1.264919 | 44860.25 |
| p.Tyr537Asp | 537 | D | -0.8226256 | -0.5885347 | 2.67117821 | 0.78570571 | 0.511430884 | | 4 | 2.576455 | 19934.25 |
| p.Tyr537Gly | 537 | G | -0.3617413 | -0.3533328 | 1.75261568 | 1.10859909 | 0.536535174 | | 4 | 1.134958 | 94413 |
| p.Tyr537Gln | 537 | Q | -0.973683 | 0.08345282 | 1.95264992 | 1.28449861 | 0.586729587 | | 4 | 1.680265 | 19643 |
| p.Tyr537Arg | 537 | R | -0.1996017 | 0.00736934 | 2.1471673 | 0.76769253 | 0.680656875 | | 4 | 1.1288 | 77406 |
| p.Tyr537Pro | 537 | P | -0.4502443 | -0.0214408 | 2.7105613 | 1.04135495 | 0.820057791 | | 4 | 1.981588 | 58374 |
| p.Tyr537Thr | 537 | T | 0.03229442 | 0.18582107 | 2.21311129 | 0.96295016 | 0.848544234 | | 4 | 0.993533 | 67677.75 |
| p.Tyr537Ala | 537 | A | -0.4084398 | 0.28599671 | 2.33073041 | 1.69915317 | 0.976860113 | | 4 | 1.583673 | 60256.5 |
| p.Tyr537Asn | 537 | N | 0.09247037 | -0.0088282 | 2.29624145 | 1.62434855 | 1.001058041 | | 4 | 1.303798 | 39048.25 |
| p.Tyr537Leu | 537 | L | 0.08992697 | -0.0206865 | 2.78331779 | 1.67285018 | 1.131352116 | | 4 | 1.811323 | 49714.25 |
| p.Tyr537Val | 537 | V | 0.18040954 | 0.43894457 | 2.30375477 | 1.71881125 | 1.160480034 | | 4 | 1.033321 | 97554 |
| p.Tyr537Met | 537 | M | 0.36564291 | 0.35059994 | 2.41680789 | 1.80228662 | 1.233834343 | | 4 | 1.085475 | 36644 |
| p.Tyr537Ile | 537 | I | 0.58200793 | 0.31426077 | 2.50513962 | 2.1003296 | 1.375434481 | | 4 | 1.185774 | 20624.75 |
| p.Tyr537Trp | 537 | W | 0.72496523 | 0.56888104 | 3.14684139 | 2.38175833 | 1.705611496 | | 4 | 1.596047 | 22062.25 |
| p.Tyr537His | 537 | H | 0.59518594 | 0.37780459 | 3.32633776 | 2.67513239 | 1.743615172 | | 4 | 2.185688 | 8791.25 |
| p.Tyr541Gln | 541 | Q | -0.8601014 | -0.0646969 | 1.46937638 | 2.12817254 | 0.668187658 | | 4 | 1.882082 | 8120.25 |
| p.Tyr541Gly | 541 | G | 0.63189818 | 0.88506759 | 1.8423365 | -0.0260313 | 0.833317743 | | 4 | 0.599949 | 12022.5 |
| p.Tyr541Pro | 541 | P | 0.1256079 | 0.74955105 | 1.99743064 | 0.64356123 | 0.879037705 | | 4 | 0.630225 | 13027 |
| p.Tyr541Thr | 541 | T | 0.00319939 | 0.17832681 | 3.23691775 | 0.42048017 | 0.959731029 | | 4 | 2.333972 | 12486.75 |
| p.Tyr541Arg | 541 | R | 0.07084204 | 0.36701508 | 2.64879411 | 0.79619138 | 0.970710654 | | 4 | 1.340211 | 16431.75 |
| p.Tyr541Glu | 541 | E | -0.297934 | 0.50823671 | 2.59553174 | 2.12429798 | 1.232533115 | | 4 | 1.839982 | 6943.25 |
| p.Tyr541Leu | 541 | L | 0.35282011 | 0.23164802 | 2.97365961 | 1.7283242 | 1.321612987 | | 4 | 1.673752 | 39058.25 |
| p.Tyr541Ala | 541 | A | -0.0375247 | 1.00152103 | 2.86074213 | 1.57700122 | 1.350434921 | | 4 | 1.460178 | 7932.75 |
| p.Tyr541Phe | 541 | F | 0.47862932 | 0.94615832 | 3.98272201 | 1.02778316 | 1.608823201 | | 4 | 2.563155 | 4934 |

| | | | | | | | | | | |
|---|---|---|---|---|---|---|---|---|---|---|
| p.Ser557Thr | 557 | T | -0.2316991 | -0.1815544 | 0.94991477 | 1.52229488 | 0.514739045 | 4 | 0.748847 | 14487 |
| p.Ser557Ile | 557 | I | 0.49595749 | 0.82105827 | 1.85194507 | -0.6164021 | 0.638139679 | 4 | 1.033625 | 10411.5 |
| p.Ser557Met | 557 | M | 0.03182948 | 0.40296027 | 1.95041797 | 0.42199755 | 0.701801319 | 4 | 0.725167 | 10875.75 |
| p.Ser557Asn | 557 | N | -0.352761 | -0.7452599 | 2.32818709 | 1.93530939 | 0.791368877 | 4 | 2.44689 | 6151 |
| p.Ser557Lys | 557 | K | 0.58913818 | 0.64198245 | 1.09987193 | 1.50403523 | 0.958756945 | 4 | 0.184735 | 10259 |
| p.Ser557Pro | 557 | P | 0.09389787 | 0.02300862 | 2.76660489 | 1.13216361 | 1.003918746 | 4 | 1.637944 | 25246.75 |
| p.Ser557Val | 557 | V | -0.0574615 | 0.71088196 | 1.3821632 | 2.34307978 | 1.094665851 | 4 | 1.038627 | 16993.5 |
| p.Ser557Asp | 557 | D | 0.45409882 | 0.8645847 | 2.0694214 | 1.08885339 | 1.119239579 | 4 | 0.470343 | 5110.75 |
| p.Ser557Leu | 557 | L | -0.3445992 | 0.72230819 | 3.03525085 | 1.15167474 | 1.141158635 | 4 | 1.990203 | 21249.25 |
| p.Ser557Arg | 557 | R | 0.28409764 | 0.59419935 | 1.95279948 | 1.85930331 | 1.172599944 | 4 | 0.734752 | 31740.25 |
| p.Ser557Trp | 557 | W | -0.3875292 | 0.18767883 | 3.42066205 | 1.68167414 | 1.225621458 | 4 | 2.901922 | 7329.5 |
| p.Ser557Glu | 557 | E | 0.20362438 | -0.4489716 | 3.24345404 | 2.21850518 | 1.304152992 | 4 | 2.960513 | 6518.75 |
| p.Ser557Gly | 557 | G | 0.28295996 | 0.61965919 | 3.21723072 | 2.99493288 | 1.778695689 | 4 | 2.376402 | 24065.5 |
| p.Ser557Ala | 557 | A | 0.98500962 | 0.4164597 | 2.88719006 | 3.25762161 | 1.886570247 | 4 | 1.951686 | 25092.25 |
| p.Ser557Tyr | 557 | Y | 1.13726461 | 1.21175211 | 2.33184356 | 3.10186886 | 1.945682283 | 4 | 0.892694 | 14965.5 |
| p.Ser557His | 557 | H | 1.19624731 | 0.98082769 | 2.39689015 | 4.05311134 | 2.156769124 | 4 | 1.986404 | 1832.75 |
| p.Ser559Val | 559 | V | -0.2023319 | 0.12296002 | 2.31688664 | 0.02480339 | 0.565579545 | 4 | 1.381705 | 13361 |
| p.Ser559Arg | 559 | R | 0.65945014 | 0.45337877 | 2.3315461 | 1.06865713 | 1.128258035 | 4 | 0.708899 | 18081 |
| p.Ser559Gly | 559 | G | -0.2295371 | 0.55019708 | 2.56786007 | 2.07745318 | 1.241493297 | 4 | 1.699966 | 13262.25 |
| p.Ser559Met | 559 | M | 0.47963891 | 0.55413586 | 2.68499132 | 1.96247399 | 1.420310017 | 4 | 1.17616 | 16941.25 |
| p.Ser559Leu | 559 | L | 0.81328672 | 0.23135459 | 2.56157666 | 2.12862692 | 1.43371122 | 4 | 1.195192 | 30608 |
| p.Ser559Thr | 559 | T | 0.29688363 | 0.69457324 | 2.7844224 | 2.1031658 | 1.469761268 | 4 | 1.368698 | 8942.75 |
| p.Ser559Ile | 559 | I | 0.5570385 | 0.53522227 | 3.05703095 | 1.74616732 | 1.473864759 | 4 | 1.434061 | 6005.25 |
| p.Ser559Gln | 559 | Q | 0.61478171 | 0.3860928 | 2.63787403 | 2.5222636 | 1.540253035 | 4 | 1.452567 | 10669 |
| p.Ser559Lys | 559 | K | 0.2929117 | 0.11658214 | 3.15110113 | 2.61915119 | 1.544936538 | 4 | 2.447155 | 8070 |
| p.Ser559Pro | 559 | P | 0.81947051 | 0.60595105 | 2.75357656 | 2.30556562 | 1.621140932 | 4 | 1.141378 | 26636.75 |
| p.Ser559Ala | 559 | A | 0.9457514 | 1.0050511 | 2.42893449 | 2.39034661 | 1.6925209 | 4 | 0.686515 | 16224.25 |
| p.Ser559Cys | 559 | C | 0.47257591 | 2.2217143 | 1.85561009 | 2.94868122 | 1.874645381 | 4 | 1.080057 | 4477.75 |
| p.Ser559Asn | 559 | N | 2.12432814 | 0.41693453 | 3.89812039 | 1.21584772 | 1.913807694 | 4 | 2.236531 | 3672 |
| p.Tyr566Phe | 566 | F | 0.11395619 | 0.59717845 | 1.87890818 | -0.5691085 | 0.505233568 | 4 | 1.067582 | 2009.75 |
| p.Tyr566Leu | 566 | L | 0.18795727 | 0.74338202 | 1.28640891 | 0.6671706 | 0.721229703 | 4 | 0.202407 | 14970.25 |
| p.Tyr566Arg | 566 | R | -0.3938766 | 0.33870381 | 2.03380271 | 1.2692092 | 0.811959773 | 4 | 1.126663 | 20662 |
| p.Tyr566Pro | 566 | P | 0.44507754 | -0.221618 | 2.41878963 | 0.67033544 | 0.828146157 | 4 | 1.267932 | 15836.25 |
| p.Tyr566Asn | 566 | N | -0.4859132 | 0.70427698 | 2.37223318 | 0.81738428 | 0.851995304 | 4 | 1.374714 | 6904 |
| p.Tyr566Asp | 566 | D | -0.2971855 | 0.51346337 | 1.78916181 | 1.64852763 | 0.913491832 | 4 | 0.977613 | 11330 |

| | | | | | | | | | | |
|---|---|---|---|---|---|---|---|---|---|---|
| p.Tyr566Glu | 566 | E | -0.6223983 | 1.08686084 | 2.72892828 | 0.70998115 | 0.97584299 | 4 | 1.903564 | 6881.5 |
| p.Tyr566Lys | 566 | K | 0.7010299 | 0.73645163 | 2.816419 | 0.01069989 | 1.066150106 | 4 | 1.473143 | 6680 |
| p.Tyr566Ile | 566 | I | 0.75027995 | -0.2445518 | 2.31078754 | 1.97945127 | 1.19899173 | 4 | 1.376789 | 12506.5 |
| p.Tyr566Thr | 566 | T | -0.1922579 | 0.20662869 | 3.14532303 | 1.89205209 | 1.262936485 | 4 | 2.390847 | 15466.25 |
| p.Tyr566Gln | 566 | Q | 0.5628343 | 1.0009999 | 2.28540222 | 1.24549452 | 1.273682736 | 4 | 0.534677 | 19380.5 |
| p.Tyr566Ala | 566 | A | 0.2559881 | 0.80086158 | 1.82318434 | 2.32017299 | 1.300051754 | 4 | 0.884525 | 13086.25 |
| p.Tyr566His | 566 | H | 0.48643624 | 0.40218096 | 2.01851058 | 2.38484875 | 1.322994134 | 4 | 1.053001 | 29039.75 |
| p.Tyr566Ser | 566 | S | 0.08271105 | 0.51971132 | 2.72815309 | 2.0180853 | 1.33716519 | 4 | 1.546795 | 25248 |
| p.Tyr566Gly | 566 | G | 0.65647568 | -0.1597876 | 2.17508671 | 3.34756792 | 1.504835675 | 4 | 2.445195 | 10051.5 |
| p.Tyr566Val | 566 | V | 0.89050996 | 0.88166398 | 2.4664096 | 2.73395684 | 1.743135097 | 4 | 0.991319 | 13890.5 |
| p.Tyr597Pro | 597 | P | 0.49335271 | 0.45680807 | 2.16273341 | 1.16217942 | 1.068768402 | 4 | 0.637028 | 60402.25 |
| p.Tyr597Glu | 597 | E | 0.4805505 | 0.65847864 | 2.51063739 | 0.7999963 | 1.112415709 | 4 | 0.885981 | 15366.25 |
| p.Tyr597Arg | 597 | R | 0.32043066 | 0.45309543 | 2.54181688 | 1.5998503 | 1.228798315 | 4 | 1.096181 | 17100.75 |
| p.Tyr597Ala | 597 | A | 0.42488137 | 0.75340572 | 3.09365911 | 0.86057732 | 1.283130881 | 4 | 1.491255 | 17662.75 |
| p.Tyr597Met | 597 | M | 0.05033647 | 1.80846854 | 2.53303471 | 0.86209789 | 1.313484403 | 4 | 1.177202 | 3684.5 |
| p.Tyr597Ile | 597 | I | -0.0801703 | 1.07613796 | 1.84600835 | 2.42857298 | 1.317637237 | 4 | 1.175181 | 7981.5 |
| p.Tyr597Trp | 597 | W | 0.63226822 | 1.67011062 | 3.10852446 | 0.17891135 | 1.397453661 | 4 | 1.69082 | 2862.25 |
| p.Tyr597Gln | 597 | Q | 0.27508752 | 0.64947698 | 2.0981399 | 2.89339303 | 1.479024357 | 4 | 1.507119 | 24148.25 |
| p.Tyr597Gly | 597 | G | 0.56676938 | 0.48017856 | 1.60090404 | 3.27749702 | 1.481337253 | 4 | 1.69308 | 6381.5 |
| p.Tyr597Asn | 597 | N | 0.72796279 | 0.75038847 | 2.53032724 | 2.05735809 | 1.516509147 | 4 | 0.84303 | 13563.5 |
| p.Tyr597Val | 597 | V | 0.75045178 | 1.1769706 | 2.71416421 | 1.52807567 | 1.542415567 | 4 | 0.711319 | 12883.75 |
| p.Tyr597Thr | 597 | T | 0.87299472 | 0.15327503 | 3.44251824 | 1.72694796 | 1.548933988 | 4 | 2.007369 | 7372 |
| p.Tyr597Leu | 597 | L | 0.47476384 | 0.74097197 | 3.20623027 | 2.50690655 | 1.732218158 | 4 | 1.778871 | 9215 |
| p.Tyr597Lys | 597 | K | 0.20261542 | 1.20358717 | 3.16773744 | 2.92961067 | 1.875887677 | 4 | 2.010345 | 14196 |
| p.Tyr597Ser | 597 | S | 1.04329826 | 0.65643426 | 3.55521516 | 2.38238831 | 1.909333995 | 4 | 1.750827 | 9402.75 |
| p.Tyr597His | 597 | H | 1.53319444 | 0.66279664 | 2.44131682 | 3.53728725 | 2.043648789 | 4 | 1.518804 | 6320.5 |
| p.Thr598Tyr | 598 | Y | 1.06608919 | -1.130124 | 0.97982212 | -1.3457748 | -0.10749688 | 4 | 1.712888 | 1900.75 |
| p.Thr598Lys | 598 | K | -0.0202658 | 0.37263994 | 3.43063435 | -0.2093165 | 0.893422974 | 4 | 2.919839 | 6911 |
| p.Thr598Pro | 598 | P | -0.1193878 | 0.29700302 | 2.65111484 | 1.03137593 | 0.965026485 | 4 | 1.489835 | 20660.75 |
| p.Thr598Met | 598 | M | 0.26009718 | 0.01272857 | 3.36042612 | 0.58599632 | 1.054812048 | 4 | 2.417718 | 11487 |
| p.Thr598Ala | 598 | A | 0.78943552 | 0.61606448 | 2.29893645 | 1.06556587 | 1.192500579 | 4 | 0.578351 | 19747 |
| p.Thr598Ser | 598 | S | 0.88809647 | 0.15017822 | 2.55585291 | 1.17878183 | 1.193227355 | 4 | 1.012671 | 9152 |
| p.Thr598Leu | 598 | L | 0.02685783 | 0.56165166 | 2.42635769 | 1.95726766 | 1.243033711 | 4 | 1.284584 | 50317.25 |
| p.Thr598Glu | 598 | E | -0.1622776 | 0.3334464 | 3.03369693 | 1.87557455 | 1.270110057 | 4 | 2.1353 | 17292.25 |
| p.Thr598Gln | 598 | Q | -0.1265205 | 0.04334891 | 2.89254282 | 2.35549652 | 1.291216929 | 4 | 2.421363 | 27137 |

| | | | | | | | | | | |
|---|---|---|---|---|---|---|---|---|---|---|
| p.Thr598Asp | 598 | D | 0.75509868 | 0.18259141 | 2.14615684 | 2.94872612 | 1.508143264 | 4 | 1.602168 | 4387.75 |
| p.Thr598Ile | 598 | I | 1.09778589 | 0.87538089 | 0.85581641 | 3.32950131 | 1.539621127 | 4 | 1.435898 | 3904.75 |
| p.Thr598Trp | 598 | W | 0.19565961 | 0.69824216 | 2.67430999 | 2.77199298 | 1.585051185 | 4 | 1.770718 | 6083 |
| p.Thr598His | 598 | H | 1.01231155 | 0.2373419 | 2.63282214 | 2.47486698 | 1.589335644 | 4 | 1.344625 | 10959.25 |
| p.Thr598Gly | 598 | G | 1.27266762 | 0.85102642 | 3.11922156 | 1.42240949 | 1.666331271 | 4 | 0.996694 | 7654 |
| p.Thr598Val | 598 | V | 0.82483521 | 1.54158331 | 3.02533578 | 1.86680611 | 1.814640102 | 4 | 0.840926 | 11276.75 |
| p.Thr598Cys | 598 | C | 0.47861864 | 1.57359474 | 3.02926459 | 4.04370002 | 2.281294497 | 4 | 2.472004 | 4381.75 |
| p.Lys603Trp | 603 | W | -0.1009289 | 0.33126047 | 1.40525648 | -0.9440436 | 0.172886111 | 4 | 0.955442 | 2917 |
| p.Lys603His | 603 | H | 0.22299551 | 0.49626063 | 2.99282597 | -0.8321756 | 0.719976639 | 4 | 2.62402 | 9278.25 |
| p.Lys603Pro | 603 | P | 0.0379466 | -0.1713465 | 2.12281241 | 1.74329953 | 0.933178015 | 4 | 1.364313 | 12765 |
| p.Lys603Thr | 603 | T | 0.30550441 | 1.00691389 | 1.67312269 | 1.39552857 | 1.095267388 | 4 | 0.351868 | 24077.75 |
| p.Lys603Leu | 603 | L | -0.0294295 | 0.24366386 | 2.51216668 | 2.05380435 | 1.195051353 | 4 | 1.62558 | 27835.25 |
| p.Lys603Gln | 603 | Q | 0.24421239 | 0.65200633 | 3.56592093 | 0.32760531 | 1.197436242 | 4 | 2.524152 | 17649.25 |
| p.Lys603Glu | 603 | E | 0.55668354 | 0.60995732 | 2.14775362 | 2.17583527 | 1.372557439 | 4 | 0.831131 | 16279.25 |
| p.Lys603Ser | 603 | S | 0.4276756 | 0.61570529 | 2.01329682 | 2.47541459 | 1.383023076 | 4 | 1.030677 | 17375 |
| p.Lys603Met | 603 | M | 0.32508982 | 0.27673521 | 2.57278332 | 2.45094232 | 1.406387669 | 4 | 1.632298 | 35396.75 |
| p.Lys603Asn | 603 | N | 0.95596723 | 2.06756328 | 2.44864103 | 0.19401062 | 1.416545541 | 4 | 1.065256 | 5432 |
| p.Lys603Arg | 603 | R | 0.56451814 | 0.85695684 | 2.33659054 | 2.06536457 | 1.455857522 | 4 | 0.766786 | 24546.75 |
| p.Lys603Gly | 603 | G | 0.90201972 | 0.05874421 | 3.01819711 | 2.00224777 | 1.495302204 | 4 | 1.663962 | 31361.75 |
| p.Lys603Asp | 603 | D | 0.08929223 | 1.07385485 | 2.7589919 | 2.16558607 | 1.521931262 | 4 | 1.399279 | 4670.75 |
| p.Lys603Ile | 603 | I | 2.1276142 | 0.91766507 | 1.07038933 | 2.52025681 | 1.658981351 | 4 | 0.619134 | 3069.25 |
| p.Lys603Ala | 603 | A | 0.00137671 | 0.82155606 | 2.97336487 | 2.99972797 | 1.699006405 | 4 | 2.322577 | 22478.5 |
| p.Lys603Val | 603 | V | 0.45757745 | 0.53066794 | 3.28631909 | 2.67930375 | 1.738467057 | 4 | 2.126825 | 26524.25 |
| p.Ala608Thr | 608 | T | 3.01143058 | -0.5131718 | 2.0630098 | 2.3616378 | 1.730726591 | 4 | 2.394581 | 1261.5 |
| p.Lys612Pro | 612 | P | -0.2114629 | -0.1738507 | 1.08746284 | -0.834631 | -0.033120425 | 4 | 0.649912 | 6200 |
| p.Lys612Asp | 612 | D | -0.525911 | -0.7698508 | 1.77811158 | 0.62426611 | 0.276653973 | 4 | 1.371497 | 32412 |
| p.Lys612Glu | 612 | E | -0.0402317 | -0.6145232 | 2.62359935 | 1.52938387 | 0.874557061 | 4 | 2.180715 | 21208.25 |
| p.Lys612Ile | 612 | I | 1.25499709 | 0.65426702 | 1.32365484 | 0.33836498 | 0.892820984 | 4 | 0.22704 | 13295 |
| p.Lys612His | 612 | H | -0.6731087 | 0.03633281 | 2.79541531 | 1.50172562 | 0.915091249 | 4 | 2.391451 | 12689.5 |
| p.Lys612Leu | 612 | L | -0.3864037 | 0.74364394 | 2.49557618 | 0.98842048 | 0.960309213 | 4 | 1.406138 | 10681.5 |
| p.Lys612Met | 612 | M | 0.28144605 | 0.38639929 | 2.36094176 | 1.02156801 | 1.012588775 | 4 | 0.91494 | 46399 |
| p.Lys612Trp | 612 | W | 0.24499922 | 0.03262452 | 2.32577016 | 1.529269 | 1.033165724 | 4 | 1.179745 | 9713.75 |
| p.Lys612Ala | 612 | A | 0.08925988 | 0.52647467 | 2.09345976 | 1.42584085 | 1.033758792 | 4 | 0.808703 | 30255.25 |
| p.Lys612Gly | 612 | G | 0.00922103 | 0.2671557 | 2.21193385 | 1.78778871 | 1.069024821 | 4 | 1.196347 | 69200.5 |
| p.Lys612Val | 612 | V | 0.22582968 | 0.32958247 | 2.20571368 | 1.63132583 | 1.098112918 | 4 | 0.954204 | 21046.75 |

| | | | | | | | | | | | |
|---|---|---|---|---|---|---|---|---|---|---|---|
| p.Lys612Thr | 612 | T | -0.2779994 | -0.068896 | 2.28092369 | 2.51532831 | 1.112339151 | | 4 | 2.220775 | 29888.75 |
| p.Lys612Tyr | 612 | Y | 0.44212053 | 0.3189788 | 2.1204716 | 1.66731885 | 1.137222445 | | 4 | 0.800157 | 13780.25 |
| p.Lys612Asn | 612 | N | 0.67030866 | 0.11833829 | 2.12222826 | 2.0876329 | 1.249627028 | | 4 | 1.02637 | 10106.25 |
| p.Lys612Arg | 612 | R | 0.34610102 | 0.41889899 | 1.66636904 | 2.66970117 | 1.275267555 | | 4 | 1.231374 | 21756.75 |
| p.Lys612Ser | 612 | S | 0.57168712 | 0.19079859 | 2.96323899 | 1.68407065 | 1.352448835 | | 4 | 1.554546 | 34396.25 |
| p.Lys612Gln | 612 | Q | 0.24361185 | 0.93978621 | 2.98445915 | 1.36708625 | 1.383735864 | | 4 | 1.353189 | 21019.25 |
| p.Gln613Lys | 613 | K | 0.86622615 | -0.3307613 | 2.13796526 | 1.60222757 | 1.068914409 | | 4 | 1.142489 | 5119.25 |
| p.Arg614Cys | 614 | C | 0.38441222 | 0.5849625 | 2.56247849 | 0.74082049 | 1.068168426 | | 4 | 1.01371 | 8276 |
| p.Ala616Val | 616 | V | -0.253033 | -0.2462726 | 2.50382574 | 1.24972005 | 0.81356004 | | 4 | 1.769367 | 3570.75 |
| p.Ala618Thr | 618 | T | -1.6407942 | 0.05658353 | 0.36709441 | -1.1732377 | -0.597588505 | | 4 | 0.926068 | 3498.75 |
| p.Ala618Asp | 618 | D | 0.39221473 | -1.0068277 | -0.0155969 | -1.1543281 | -0.446134499 | | 4 | 0.568036 | 3304 |
| p.Ala618Val | 618 | V | -0.2308839 | -0.8297563 | 2.40209844 | 1.19264508 | 0.633525823 | | 4 | 2.109621 | 862.75 |
| p.Ala618Arg | 618 | R | 0.22746753 | 0.46120911 | 1.830075 | 0.84285676 | 0.840402099 | | 4 | 0.499645 | 8807.25 |
| p.Ala618Leu | 618 | L | -0.2662801 | 0.39104202 | 1.15754128 | 2.41063232 | 0.923233888 | | 4 | 1.321809 | 1573.75 |
| p.Ala618Pro | 618 | P | -0.0104678 | -0.0692947 | 2.22443157 | 2.09472985 | 1.059849721 | | 4 | 1.615925 | 12741.25 |
| p.Ala618Gly | 618 | G | -0.4010398 | 1.03250743 | 3.22119367 | 0.74436615 | 1.149256857 | | 4 | 2.291303 | 6242.75 |
| p.Ala618Asn | 618 | N | 0.23886548 | -0.0692904 | 2.52356196 | 2.58816493 | 1.320325496 | | 4 | 2.051928 | 2528.5 |
| p.Ala618Cys | 618 | C | 0.6630957 | -0.5677587 | 2.52356196 | 2.97705758 | 1.398989137 | | 4 | 2.721533 | 3373.75 |
| p.Ala618Ser | 618 | S | 1.40901371 | 0.94832397 | 4.10991769 | 0.82415016 | 1.822851383 | | 4 | 2.388045 | 3108.75 |
| p.Ala618Met | 618 | M | 1.96834268 | 0.9798971 | 3.46512273 | 2.09848986 | 2.127963093 | | 4 | 1.044133 | 3564.25 |
| p.Pro625Ser | 625 | S | 0.44277055 | 0.1492702 | 1.5321458 | 0.74000765 | 0.71604855 | | 4 | 0.354169 | 7124.75 |
| p.Pro625Gln | 625 | Q | -0.7289562 | 0.49786393 | 3.0630098 | 0.97481167 | 0.951682308 | | 4 | 2.496245 | 4599.75 |
| p.Pro625Lys | 625 | K | 0.83428003 | 0.11590897 | 3.32192809 | 1.48542683 | 1.43938598 | | 4 | 1.887943 | 1656.25 |
| p.Thr634Arg | 634 | R | -0.3730822 | 0.43295941 | 1.63471554 | 1.13437742 | 0.707242546 | | 4 | 0.761661 | 3109.5 |
| p.Thr634Leu | 634 | L | 1.2108508 | 1.11933882 | 2.75607442 | -0.107294 | 1.244742507 | | 4 | 1.376334 | 3241.75 |
| p.Thr634Val | 634 | V | 0.3958415 | 0.66186168 | 2.78427131 | 3.80632406 | 1.912074637 | | 4 | 2.736968 | 3876.25 |
| p.Thr634Ser | 634 | S | 2.69839073 | 1.02779845 | 2.84799691 | 3.32736198 | 2.475387016 | | 4 | 1.003314 | 2169.75 |
| p.Ser635Gln | 635 | Q | 0.00479117 | 0.91817578 | 2.58084639 | -0.0280304 | 0.86894574 | | 4 | 1.494786 | 12507.75 |
| p.Ser635Gly | 635 | G | -0.0563825 | -0.1275599 | 2.23878686 | 2.00411025 | 1.01473868 | | 4 | 1.643099 | 9598.25 |
| p.Ser635Arg | 635 | R | 0.06223834 | 0.17016676 | 2.83163648 | 1.28544983 | 1.087372851 | | 4 | 1.657953 | 8536.75 |
| p.Ser635Glu | 635 | E | -0.0529489 | 0.37101876 | 2.98114097 | 1.44131682 | 1.185131919 | | 4 | 1.828968 | 6092 |
| p.Ser635Thr | 635 | T | 0.50315951 | -0.4556545 | 3.24260251 | 1.78173061 | 1.26795952 | | 4 | 2.57298 | 13257 |
| p.Ser635Asp | 635 | D | 0.47529802 | -0.0942393 | 3.18476004 | 1.55667346 | 1.280623049 | | 4 | 2.080245 | 5050.5 |
| p.Ser635Met | 635 | M | 0.85431445 | -0.5379139 | 3.28540222 | 2.02633415 | 1.407034222 | | 4 | 2.666707 | 4442.5 |
| p.Ser635Leu | 635 | L | 0.21924438 | 0.31762188 | 3.2098046 | 2.1069152 | 1.463396516 | | 4 | 2.108257 | 11420.25 |

| | | | | | | | | | | |
|---|---|---|---|---|---|---|---|---|---|---|
| p.Ser635His | 635 | H | 2.21824078 | 0.57746796 | 1.8251662 | 2.27177426 | 1.723162299 | 4 | 0.623033 | 3303.25 |
| p.Ser635Ala | 635 | A | 1 | 1.0816838 | 3.1827948 | 3.05498923 | 2.079866958 | 4 | 1.443265 | 4178.25 |
| p.Asp638Tyr | 638 | Y | 0.09123237 | 0.00596567 | 2.50901365 | 0.60448506 | 0.802674186 | 4 | 1.363922 | 5182 |
| p.Asp638Met | 638 | M | 0.49538441 | 0.63844483 | 2.16308757 | 0.79475904 | 1.022918963 | 4 | 0.592718 | 7201 |
| p.Asp638Gln | 638 | Q | -0.7282484 | 1.40178731 | 3.28703811 | 0.98647162 | 1.236762164 | 4 | 2.718259 | 4302.5 |
| p.Asp638Glu | 638 | E | 0.4426477 | 1.46318831 | 2.96962635 | 1.80096659 | 1.669107239 | 4 | 1.085114 | 3074 |
| p.Asp638Val | 638 | V | 0.94117303 | 0.43063435 | 2.73360658 | 3.3758669 | 1.870320216 | 4 | 1.982648 | 5932.75 |
| p.Thr655Lys | 655 | K | 0.01282404 | -0.3221935 | 2.65662349 | 0.66834365 | 0.753899415 | 4 | 1.778282 | 6147.75 |
| p.Thr655His | 655 | H | 0.24135976 | 0.73222629 | 2.02384674 | 0.30896342 | 0.826599052 | 4 | 0.684253 | 15488.25 |
| p.Thr655Arg | 655 | R | 0.19913515 | -0.6340268 | 1.43989447 | 3.02966405 | 1.008666706 | 4 | 2.54139 | 32606.25 |
| p.Thr655Trp | 655 | W | 0.84981734 | 0.66774555 | 1.5212369 | 1.39231742 | 1.107779303 | 4 | 0.170694 | 1825 |
| p.Thr655Gln | 655 | Q | 0.18350828 | 0.55799545 | 1.98374411 | 1.96751018 | 1.173189507 | 4 | 0.881959 | 31406 |
| p.Thr655Val | 655 | V | 1.12093702 | 0.48533176 | 3.06169232 | 0.82232914 | 1.37257256 | 4 | 1.33547 | 8443.5 |
| p.Thr655Ser | 655 | S | 0.65566052 | 1.11243951 | 3.09294722 | 0.70940987 | 1.39261428 | 4 | 1.3265 | 11304.75 |
| p.Thr655Pro | 655 | P | 0.57581029 | 0.28457201 | 2.56862376 | 2.22197482 | 1.41274522 | 4 | 1.321381 | 23622.25 |
| p.Thr655Leu | 655 | L | 0.13574314 | 0.53880682 | 2.84852643 | 2.66356139 | 1.546659445 | 4 | 1.982926 | 43602.25 |
| p.Thr655Met | 655 | M | 0.6276489 | 1.32281021 | 3.7825199 | 3.22760071 | 2.240144931 | 4 | 2.265212 | 11517.5 |
| p.Thr655Asn | 655 | N | 0.84298565 | 1.51641073 | 4.0768156 | 3.03467896 | 2.367722736 | 4 | 2.138461 | 3349.5 |
| p.Arg662Ser | 662 | S | -0.0976849 | 0.08096884 | 0.91593574 | 2.08920017 | 0.747104968 | 4 | 0.99571 | 5374.25 |
| p.Thr665Leu | 665 | L | 0.38619298 | 0.22351726 | 0.71111881 | 1.27130202 | 0.648032768 | 4 | 0.213739 | 13053.25 |
| p.Thr665Asn | 665 | N | -0.4694426 | 0.9101907 | 2.90189486 | 0.0846624 | 0.856826327 | 4 | 2.180127 | 10104 |
| p.Thr665Phe | 665 | F | 0.47180884 | 1.55774103 | 2.19041146 | -0.6742298 | 0.886432872 | 4 | 1.586199 | 3006 |
| p.Thr665Lys | 665 | K | 0.09462899 | 0.89798212 | 1.69940218 | 1.34063607 | 1.008162341 | 4 | 0.478345 | 11783.5 |
| p.Thr665Ala | 665 | A | 0.12192879 | 0.51303421 | 2.4768137 | 1.42969262 | 1.135367328 | 4 | 1.100154 | 22977 |
| p.Thr665Arg | 665 | R | 0.46139089 | 0.52175904 | 2.45243449 | 1.17259419 | 1.152044652 | 4 | 0.855233 | 16810.75 |
| p.Thr665His | 665 | H | 0.94939532 | 1.56921431 | 1.91822207 | 0.60198108 | 1.259703196 | 4 | 0.352778 | 6173.25 |
| p.Thr665Ser | 665 | S | -0.6450091 | 1.03600415 | 2.5834956 | 2.3190741 | 1.323391189 | 4 | 2.178813 | 13219.75 |
| p.Thr665Gly | 665 | G | 0.66711294 | 0.91668822 | 2.34221237 | 1.52423475 | 1.362562069 | 4 | 0.556102 | 26875.25 |
| p.Thr665Pro | 665 | P | 0.16578161 | 0.87130033 | 2.8704486 | 1.62539009 | 1.383230158 | 4 | 1.338238 | 13020 |
| p.Thr665Ile | 665 | I | 0.65109999 | 0.45424278 | 2.95450045 | 1.51461219 | 1.393613851 | 4 | 1.294917 | 31284.75 |
| p.Thr665Gln | 665 | Q | 0.14712687 | 0.21348137 | 2.7548875 | 2.5849625 | 1.425114561 | 4 | 2.071617 | 1504.75 |
| p.Thr665Val | 665 | V | 0.84174906 | 1.34603233 | 2.15484305 | 1.64650859 | 1.497283255 | 4 | 0.302418 | 9124 |
| p.Thr665Asp | 665 | D | 0.5207988 | 0.23564018 | 2.26220788 | 3.11179217 | 1.532609757 | 4 | 1.910674 | 18249 |
| p.Thr665Glu | 665 | E | 0.13455178 | 0.58466396 | 3.95020177 | 1.65745222 | 1.581717434 | 4 | 2.901286 | 7303.5 |
| p.Thr665Met | 665 | M | 0.00932432 | 1.08297115 | 2.91767834 | 2.59661035 | 1.651646039 | 4 | 1.838802 | 20396 |

| | | | | | | | | | | |
|---|---|---|---|---|---|---|---|---|---|---|
| p.Gln667Lys | 667 | K | -0.2791832 | 0.19382327 | 1.63005039 | -0.8519014 | 0.173197281 | 4 | 1.126107 | 1458.25 |
| p.Gln667Glu | 667 | E | -0.2319845 | 1.85173578 | 1.57531233 | 1.70373229 | 1.22469898 | 4 | 0.955835 | 3858.5 |
| p.Ala669Thr | 669 | T | 0.26047417 | 2.37772989 | 2.67670507 | 4.07140926 | 2.346579596 | 4 | 2.478942 | 2520.5 |
| p.Lys679Tyr | 679 | Y | 0.75073375 | 0.32570153 | 2.08618804 | -0.5238896 | 0.65968342 | 4 | 1.185198 | 11452.5 |
| p.Lys679Trp | 679 | W | 0.60494207 | 0.68413791 | 1.6520767 | -0.2016339 | 0.684880702 | 4 | 0.575922 | 3200 |
| p.Lys679Ile | 679 | I | 0.8004564 | 1.13464953 | 1.19793938 | 0.60279579 | 0.933960276 | 4 | 0.079151 | 5146.5 |
| p.Lys679Val | 679 | V | -0.2726427 | 0.45864873 | 2.74088064 | 1.38717071 | 1.078514355 | 4 | 1.68953 | 6524 |
| p.Lys679Gly | 679 | G | 0.07982268 | 1.00349603 | 2.41919894 | 0.96679763 | 1.117328822 | 4 | 0.935634 | 15302.75 |
| p.Lys679Asn | 679 | N | -0.5281765 | 3.42428981 | 1 | 0.85693695 | 1.188262572 | 4 | 2.697067 | 1311.75 |
| p.Lys679Thr | 679 | T | 0.22811088 | 0.33494286 | 2.2758017 | 2.0322764 | 1.217782962 | 4 | 1.180554 | 15512.75 |
| p.Lys679Gln | 679 | Q | -0.023004 | 1.17230608 | 2.29604946 | 1.55019708 | 1.248887149 | 4 | 0.936969 | 5256.5 |
| p.Lys679His | 679 | H | 0.47604373 | 0.17989549 | 2.65983042 | 2.1038113 | 1.354895236 | 4 | 1.472245 | 17236.5 |
| p.Lys679Arg | 679 | R | 1.11375829 | 0.55312339 | 2.80527161 | 1.58878493 | 1.515234556 | 4 | 0.918815 | 20488.75 |
| p.Lys679Ser | 679 | S | 0.77454013 | 1.0168559 | 2.91038381 | 1.37939563 | 1.520293864 | 4 | 0.9206 | 10868.25 |
| p.Lys679Ala | 679 | A | 1.19150964 | 0.40678278 | 2.08048992 | 2.96818355 | 1.661741475 | 4 | 1.22606 | 4411.5 |
| p.Lys679Glu | 679 | E | 0.36965485 | 0.85214317 | 3.44183756 | 2.18315726 | 1.711698209 | 4 | 1.918524 | 11635.25 |
| p.Lys679Pro | 679 | P | 0.71910761 | 0.91090983 | 2.90573134 | 2.627816 | 1.790891192 | 4 | 1.2888 | 20621.5 |
| p.Lys679Met | 679 | M | 0.87047996 | 0.66149528 | 2.48202898 | 3.37426093 | 1.847066286 | 4 | 1.698267 | 8866 |
| p.Lys682Asp | 682 | D | -0.6126833 | 0.77696964 | 3.09515723 | -0.4604805 | 0.699740787 | 4 | 2.937518 | 1959.25 |
| p.Lys682His | 682 | H | 0.52496349 | 0.34522754 | 1.9608294 | 1.47864121 | 1.077415412 | 4 | 0.594235 | 9792.25 |
| p.Lys682Met | 682 | M | 0.09633008 | 1.05889369 | 2.02578403 | 1.23446525 | 1.103868262 | 4 | 0.628047 | 8156.5 |
| p.Lys682Trp | 682 | W | 0.24363357 | 1.44018645 | 2.40880555 | 0.86522332 | 1.239462221 | 4 | 0.846461 | 3225.75 |
| p.Lys682Val | 682 | V | 1.20302336 | 0.8067381 | 1.66603411 | 1.73822269 | 1.353504563 | 4 | 0.189094 | 9381 |
| p.Lys682Pro | 682 | P | 0.76955745 | 0.21391476 | 1.95909461 | 2.67760086 | 1.405041918 | 4 | 1.249668 | 19883 |
| p.Lys682Ser | 682 | S | 0.59227471 | 0.37400289 | 2.74774543 | 2.26629923 | 1.495080564 | 4 | 1.41194 | 11209.75 |
| p.Lys682Thr | 682 | T | 0.72747761 | 0.15648798 | 3.52201616 | 1.57842837 | 1.496102529 | 4 | 2.165485 | 20234.75 |
| p.Lys682Ile | 682 | I | 0.16776391 | 1.86819944 | 1.55325364 | 2.70566946 | 1.573721614 | 4 | 1.115053 | 6637.5 |
| p.Lys682Gly | 682 | G | 1.64195787 | -0.1953209 | 3.59693514 | 1.2668235 | 1.57759891 | 4 | 2.440562 | 5721 |
| p.Lys682Tyr | 682 | Y | 0.58112773 | -0.0973144 | 2.32886414 | 3.59054353 | 1.600805243 | 4 | 2.80416 | 7068.75 |
| p.Lys682Ala | 682 | A | 0.38055155 | 0.48715288 | 3.5922305 | 2.03839829 | 1.624583307 | 4 | 2.294747 | 15371.75 |

Table S3 (TAP2 Synonymous variants)

| position | syn_or_mut | Score integration 1 | Score integration 2 | Score integration 3 | Score integration 4 | Score average | Number Surviving Replicates | Variance | Average read count |
|---|---|---|---|---|---|---|---|---|---|
| 2 | syn | 1.206127367 | 0.43484961 | 3.84419169 | 0.565597176 | 1.51269146 | 4 | 2.52953517 | 2768 |
| 16 | syn | 0.069231002 | 0.04270838 | 2.908638258 | -0.12417941 | 0.724099556 | 4 | 2.128311103 | 3322.75 |
| 16 | syn | 0.879342543 | 0.86943163 | 1.66930654 | 2.339850003 | 1.439482678 | 4 | 0.500731833 | 12384.75 |
| 34 | syn | 0.145339363 | 0.3408283 | 3.321928095 | 1.423373358 | 1.307867279 | 4 | 2.118806055 | 2094.25 |
| 34 | syn | 0.071960338 | 1.09594611 | 2.544320516 | 2.635492717 | 1.58692992 | 4 | 1.517426211 | 3356.25 |
| 43 | syn | 0.871034443 | 0.15437734 | 2.695993813 | -1.23224021 | 0.622291348 | 4 | 2.673448616 | 2146.5 |
| 43 | syn | -0.661831264 | 0.70092852 | 3.065095028 | -0.11594148 | 0.747062701 | 4 | 2.701720218 | 1830.25 |
| 84 | syn | 0.120899433 | 0.72286811 | 1.966833136 | 0.881203952 | 0.922951157 | 4 | 0.591584203 | 3426.75 |
| 101 | syn | 0.768166463 | 0.25162964 | 3.117266052 | 3.529253068 | 1.916578806 | 4 | 2.711091589 | 3320.5 |
| 111 | syn | 0.173594816 | 0.94208002 | 1.777367109 | 1.974148122 | 1.216797518 | 4 | 0.683853261 | 11753.5 |
| 111 | syn | 0.135728991 | 0.77039179 | 3.285402219 | 2.184066947 | 1.593897487 | 4 | 2.004635081 | 13554.25 |
| 132 | syn | 0.725108466 | 0.15923725 | 2.950787026 | 2.38332864 | 1.554615346 | 4 | 1.757074256 | 2381.25 |
| 135 | syn | 0.533795071 | -0.84118616 | 2.567040593 | 2.58138926 | 1.210259692 | 4 | 2.795628385 | 11021.5 |
| 141 | syn | -0.074887212 | -0.18795892 | 2.544320516 | 2.64385619 | 1.231332643 | 4 | 2.479919596 | 2278.5 |
| 144 | syn | 0.944529701 | 0.61257116 | 2.72762773 | 2.360834589 | 1.661390795 | 4 | 1.079998424 | 10037.75 |
| 145 | syn | -0.094226762 | 0.36768169 | 2.670538232 | 1.721084794 | 1.166269489 | 4 | 1.599079249 | 13215.25 |
| 173 | syn | 0.792287474 | 0.54105675 | 1.653771659 | -0.30092488 | 0.67154775 | 4 | 0.647357604 | 10118.25 |
| 199 | syn | 0.199308808 | 1.91488339 | 1.64385619 | 4.169925001 | 1.981993346 | 4 | 2.694616524 | 904.25 |
| 199 | syn | 1.027829677 | 0.53833083 | 2.72662365 | 3.974004791 | 2.066697238 | 4 | 2.496158218 | 11200.5 |
| 200 | syn | 0.566218778 | 0.09866264 | 2.662965013 | 2.02358682 | 1.337858312 | 4 | 1.452388283 | 2911 |
| 200 | syn | 0.534990812 | 1.51650406 | 2.704701745 | 4.28757659 | 2.260943302 | 4 | 2.61242194 | 5551 |
| 204 | syn | 0.918115819 | 0.67334949 | 2.906890596 | 1.283792966 | 1.445537219 | 4 | 1.012054036 | 3095 |
| 218 | syn | 1.42324003 | 1.48219834 | 2.485426827 | 2.558318511 | 1.987295926 | 4 | 0.382494606 | 5200.25 |
| 219 | syn | 0.765070932 | 0.91391102 | 3.724026538 | 1.428843299 | 1.707962948 | 4 | 1.887327998 | 3517.25 |
| 220 | syn | 1.080919995 | -0.74865468 | 2.740880644 | 0.724546335 | 0.949423074 | 4 | 2.053549757 | 5766 |
| 231 | syn | 0.591436265 | 0.68531228 | 2.03312678 | 2.610222453 | 1.480024445 | 4 | 1.001475368 | 13934 |
| 244 | syn | 0.150491627 | -0.64254999 | 2.678071905 | 0.537537155 | 0.680887673 | 4 | 2.014033816 | 1388 |
| 244 | syn | 0.270360812 | 0.63182684 | 2.846308556 | 2.245727238 | 1.498555861 | 4 | 1.544794891 | 44032.75 |
| 246 | syn | 0.14199251 | 0.95682098 | 1.4471074 | 1.50676777 | 1.013172165 | 4 | 0.398021952 | 20492.25 |
| 246 | syn | 0.519282082 | 0.59666654 | 2.743194558 | 2.404125777 | 1.56581724 | 4 | 1.374489155 | 29005.75 |
| 250 | syn | 1.240358968 | 1.21403335 | 2.353636955 | 2.325585118 | 1.783403599 | 4 | 0.412735609 | 2906.25 |
| 251 | syn | 0.610588308 | -1.28403149 | 1.900464326 | 2.083416008 | 0.827609289 | 4 | 2.411397774 | 1035 |

| | | | | | | | | | |
|---|---|---|---|---|---|---|---|---|---|
| 253 | syn | 0.607559391 | 1.94085402 | 1.787270676 | -0.17870589 | 1.03924455 | 4 | 1.014066021 | 4054.5 |
| 254 | syn | 0.294560078 | 1.13719196 | 3.795322367 | 1.938376214 | 1.791362655 | 4 | 2.235274922 | 7304.5 |
| 255 | syn | 0.573931464 | 0.38310221 | 2.55307844 | 0.589836475 | 1.024987147 | 4 | 1.046628869 | 4154.75 |
| 257 | syn | 0.83275837 | 0.75341327 | 2.417702741 | 1.581760069 | 1.396408611 | 4 | 0.602847157 | 10574.25 |
| 273 | syn | 0.705081533 | 0.70949576 | 3.682994584 | 2.892220907 | 1.997448196 | 4 | 2.323572633 | 6720.5 |
| 274 | syn | 0.729910837 | 0.34103692 | 1.750021747 | 2.678071905 | 1.374760352 | 4 | 1.107952385 | 1305.5 |
| 277 | syn | 0.512675562 | 0.10141052 | 2.891897393 | 2.824130635 | 1.582528528 | 4 | 2.198106178 | 14998.5 |
| 293 | syn | 0.632082791 | -0.54260849 | 1.818974005 | 1.728739573 | 0.90929697 | 4 | 1.227958549 | 3528 |
| 295 | syn | 0.570426408 | 0.79842843 | 1.449610648 | 3.581456557 | 1.599980511 | 4 | 1.883775269 | 6409.75 |
| 296 | syn | 0.416287127 | 0.45232811 | 2.23339142 | -0.70292584 | 0.599770204 | 4 | 1.473713489 | 9120.25 |
| 303 | syn | -0.577715975 | -0.26447782 | 0.265420999 | 2.689997971 | 0.528306293 | 4 | 2.197937167 | 6248.5 |
| 303 | syn | 0.770919997 | 0.67829013 | 4.020171001 | 2.315753927 | 1.946283763 | 4 | 2.475601441 | 13359 |
| 312 | syn | 1.594963893 | 1.07285434 | 2.917251107 | 1.426422263 | 1.752872901 | 4 | 0.649902375 | 7337.25 |
| 313 | syn | 1.366095678 | 0.08719844 | 2.178212973 | 3.925095191 | 1.889150571 | 4 | 2.583081821 | 3569.75 |
| 341 | syn | 0.415373127 | 1.68528339 | 3.104869118 | 0.287281952 | 1.373201897 | 4 | 1.730908101 | 5743.25 |
| 341 | syn | 0.16073583 | -0.12235376 | 3.296837114 | 2.207044457 | 1.385565911 | 4 | 2.700605021 | 3515.5 |
| 341 | syn | 0.205232523 | 1.05343001 | 2.402098444 | 2.485747818 | 1.536627199 | 4 | 1.218653911 | 12100.5 |
| 341 | syn | 0.519492709 | 3.05557996 | 4.070389328 | 4.053681121 | 2.92478578 | 4 | 2.796451352 | 3707.75 |
| 342 | syn | 3.010124217 | 0.99405073 | 0.144389909 | 3.229867542 | 1.844608101 | 4 | 2.29718709 | 1718.5 |
| 343 | syn | 0.235152579 | 0.44825614 | 2.081941614 | 1.91041441 | 1.168941186 | 4 | 0.92490014 | 11197.25 |
| 354 | syn | 1.301437569 | 0.92648221 | 1.710835862 | 0.947919165 | 1.221668701 | 4 | 0.135907153 | 7540.25 |
| 367 | syn | 0.532252916 | 0.55365801 | 3.431732104 | 3.195370018 | 1.928253263 | 4 | 2.568120856 | 22239.75 |
| 368 | syn | 0.610078324 | 0.63434005 | 3.098919005 | 2.328373504 | 1.667927722 | 4 | 1.557091167 | 17699 |
| 368 | syn | -0.356593764 | 2.43010975 | 2.342392197 | 2.364572432 | 1.695120153 | 4 | 1.872289051 | 2344 |
| 369 | syn | -0.067097851 | 0.13612691 | 2.491016509 | 2.618251819 | 1.294574346 | 4 | 2.126582514 | 15095 |
| 370 | syn | 0.07113995 | 0.25546119 | 2.592457037 | 2.380852236 | 1.324977602 | 4 | 1.812449861 | 5488 |
| 372 | syn | 0.740613371 | 0.56860789 | 2.126108769 | 1.887004023 | 1.330583513 | 4 | 0.623711946 | 12691.25 |
| 373 | syn | 1.007331315 | 0.11907586 | 1.142957954 | -0.44141837 | 0.456986689 | 4 | 0.564917057 | 4088.25 |
| 387 | syn | 0.337406987 | -0.14300772 | 2.466906739 | 2.773724144 | 1.358757538 | 4 | 2.176193659 | 3927.75 |
| 405 | syn | 1.063127979 | -0.04397673 | 2.95187805 | 1.432334323 | 1.350840905 | 4 | 1.53275203 | 11410.5 |
| 411 | syn | 0.671706955 | 0.48034819 | 3.647614842 | 0.354885933 | 1.288638979 | 4 | 2.490200332 | 6238.25 |
| 411 | syn | 1.674489348 | 0.90116044 | 3.404390255 | 3.079546799 | 2.264896711 | 4 | 1.390152592 | 5151 |
| 428 | syn | -0.374056566 | 1.14494264 | 2.428236997 | 3.481869008 | 1.670248021 | 4 | 2.770548289 | 4784.25 |
| 434 | syn | 0.471268118 | 0.09945093 | 0.833990049 | 0.22650853 | 0.407804407 | 4 | 0.104537302 | 4131 |
| 434 | syn | 0.847165513 | 1.25523745 | 2.942856093 | 3.547065893 | 2.148081238 | 4 | 1.692792294 | 19693.75 |

| | | | | | | | | | |
|---|---|---|---|---|---|---|---|---|---|
| 435 | syn | 0.105109117 | 0.40887921 | 2.39410073 | 3.366222474 | 1.568577883 | 4 | 2.46655196 | 13375.25 |
| 446 | syn | 1.298810072 | 2.30122682 | 3.085391491 | 3.508239071 | 2.548416863 | 4 | 0.944073491 | 4393.5 |
| 448 | syn | 0.77289904 | 0.42580758 | 1.381249186 | -0.13394312 | 0.611503172 | 4 | 0.402910194 | 4820.25 |
| 454 | syn | -0.227362096 | -0.05628913 | 1.5334322 | -1.2438704 | 0.001477643 | 4 | 1.317827048 | 3193 |
| 455 | syn | 1.587741591 | 0.7603607 | 2.966992894 | 1.56157268 | 1.719166967 | 4 | 0.839495863 | 8412.25 |
| 455 | syn | 0.560969645 | 1.22920081 | 3.975752454 | 2.280107919 | 2.011507707 | 4 | 2.215489416 | 2550 |
| 455 | syn | 0.8165002 | 0.72670361 | 2.611434712 | 4.210752506 | 2.091347757 | 4 | 2.74995229 | 4461.75 |
| 468 | syn | 0.079530998 | 0.1624747 | 2.365181293 | 0.409170492 | 0.75408937 | 4 | 1.173207712 | 6413.75 |
| 473 | syn | 0.525332786 | -0.31081892 | 1.530514717 | 0.786642722 | 0.632917826 | 4 | 0.577175038 | 3890.25 |
| 474 | syn | 1.418225777 | 1.50854939 | 3.939805218 | 2.546801506 | 2.353345471 | 4 | 1.380803012 | 2953 |
| 490 | syn | 0.369476753 | 0.41105342 | 2.286881148 | 0.256131992 | 0.830885828 | 4 | 0.946473976 | 9694.25 |
| 499 | syn | 0.348873243 | 0.79327165 | 2.888599447 | 2.215850475 | 1.561648705 | 4 | 1.416668613 | 24220.25 |
| 503 | syn | 0.91771537 | 1.46142347 | 2.509674373 | 0.350057468 | 1.309717669 | 4 | 0.845841415 | 2957.5 |
| 503 | syn | 0.163852811 | 1.00611053 | 3.014355293 | 1.159813685 | 1.336033081 | 4 | 1.443558065 | 13044.25 |
| 503 | syn | -0.076930243 | 0.62230612 | 3.757937599 | 1.674625237 | 1.494484677 | 4 | 2.795236681 | 6724 |
| 504 | syn | 0.096588883 | 0.79675981 | 2.203816692 | 0.809777584 | 0.976735742 | 4 | 0.780217505 | 15914 |
| 504 | syn | 0.827323338 | 3.64934173 | 1.304854582 | 1.442222329 | 1.805935494 | 4 | 1.579732549 | 1152.25 |
| 506 | syn | 0.970455724 | 1.18916827 | 1.852774047 | 0.453289868 | 1.116421977 | 4 | 0.336185578 | 3564 |
| 507 | syn | -0.556948125 | 1.71648766 | 2.945818962 | 3.125530882 | 1.807722345 | 4 | 2.877291149 | 7636 |
| 508 | syn | 0.450736434 | 1.63377753 | 3.279864035 | 2.756865152 | 2.030310787 | 4 | 1.580519443 | 4806.5 |
| 510 | syn | 0.944848334 | 0.93078079 | 2.217753527 | 2.07788711 | 1.542817439 | 4 | 0.491331397 | 8586.25 |
| 511 | syn | 1.191265372 | 1.45024454 | 3.658211483 | 2.327237006 | 2.156739599 | 4 | 1.238254313 | 8478.75 |
| 523 | syn | 1.26358833 | 0.6918777 | 3.434628228 | 3.909487707 | 2.324895492 | 4 | 2.511853073 | 2346.25 |
| 533 | syn | 0.288653412 | 0.76642466 | 2.172734535 | 1.441357198 | 1.167292451 | 4 | 0.672908939 | 11060 |
| 533 | syn | 0.617553751 | 0.66478998 | 3.218046951 | 2.155303917 | 1.66392365 | 4 | 1.583303952 | 24664.5 |
| 541 | syn | 0.396207095 | 0.70252089 | 3.404983835 | 2.252045695 | 1.688939378 | 4 | 1.968691833 | 5464.25 |
| 556 | syn | 1.345450818 | 2.08746284 | 2.321928095 | 3.247927513 | 2.250692317 | 4 | 0.615219536 | 1055.5 |
| 557 | syn | 0.305469672 | 0.31059226 | 2.569641701 | 1.296981738 | 1.120671344 | 4 | 1.150460778 | 9754.25 |
| 558 | syn | 0.845762984 | -0.20195883 | 1.961525852 | 0.920829787 | 0.881539948 | 4 | 0.781054279 | 3977.75 |
| 597 | syn | -0.143302478 | 0.75788625 | 3.201633861 | 1.362570079 | 1.294696927 | 4 | 1.999007744 | 948.25 |
| 598 | syn | -0.346687825 | -0.68477373 | 3.026967048 | 0.775354423 | 0.692714978 | 4 | 2.811131781 | 2770.75 |
| 598 | syn | 0.607167916 | 0.49903649 | 3.089375807 | 1.891805244 | 1.521846363 | 4 | 1.492264947 | 11185 |
| 606 | syn | 0.636995276 | 0.70736621 | 2.112474729 | 2.32414592 | 1.445245534 | 4 | 0.805131687 | 5786 |
| 612 | syn | -0.232022179 | 0.49007379 | 2.961623328 | 1.404159708 | 1.155958662 | 4 | 1.897307374 | 22895.25 |
| 634 | syn | 0.452190498 | 0.28633586 | 2.936541402 | 0.832419483 | 1.12687181 | 4 | 1.507767255 | 3429.75 |

| | | | | | | | | | |
|---|---|---|---|---|---|---|---|---|---|
| 634 | syn | 0.362278596 | 0.2781413 | 3.036525876 | 2.719388821 | 1.599083648 | 4 | 2.198633089 | 5649.75 |
| 634 | syn | 0.755077643 | 0.56495254 | 2.794415866 | 2.384049807 | 1.624623964 | 4 | 1.274718345 | 4062.25 |
| 665 | syn | 0.730362605 | 0.46150776 | 2.031526506 | 2.025053325 | 1.312112549 | 4 | 0.695934165 | 22032.75 |
| 665 | syn | 0.684405109 | 0.72075695 | 2.421242649 | 3.637455377 | 1.86596502 | 4 | 2.051365562 | 15442 |
| | | | | | Mean | 1.431581287 | | | |
| | | | | | Standard Dev | 0.511555969 | | | |
| | | | | | Mean - 1 SD | 0.920025318 | | | |

Table S4 (TAP2 Nonsense variants)

| index | position | Score integration 1 | Score integration 2 | Score integration 3 | Score integration 4 | Score average | Surviving Replicates | Variance | Average read count |
|---|---|---|---|---|---|---|---|---|---|
| p.Asp16Ter | 16 | -0.1285321 | -0.7037518 | 1.83399005 | 1.57150024 | 0.643301596 | 4 | 1.5631906 | 6071.5 |
| p.Ser78Ter | 78 | -0.0777515 | 0.496019 | 1.59367972 | 2.90516592 | 1.229278288 | 4 | 1.7291279 | 5626.5 |
| p.Leu111Ter | 111 | -0.2870419 | -0.9478245 | 1.17778712 | -0.7502217 | -0.201825258 | 4 | 0.9226153 | 10425 |
| p.Trp113Ter | 113 | 0.32862275 | -0.1040356 | 1.10128334 | 2.55458885 | 0.970114841 | 4 | 1.3643581 | 2594.75 |
| p.Gln129Ter | 129 | 0.12447358 | 1.89683946 | 2.11005355 | -0.2175914 | 0.978443787 | 4 | 1.4279189 | 3748.25 |
| p.Tyr172Ter | 172 | -0.0171966 | -0.0599848 | 1.73614797 | 3.20090572 | 1.214968068 | 4 | 2.4531038 | 14994.5 |
| p.Met196Ter | 196 | 0.04258804 | 0.6865834 | 2.93816976 | -0.5522012 | 0.778785001 | 4 | 2.328318 | 7481.25 |
| p.Ser200Ter | 200 | 1.41119543 | 0.56907113 | 2.26497221 | 1.09192249 | 1.334290315 | 4 | 0.5054619 | 1231.25 |
| p.Ser204Ter | 204 | -0.059501 | 2.31034012 | 2.39356597 | 1.95733354 | 1.650434653 | 4 | 1.3352621 | 2208.5 |
| p.Ser219Ter | 219 | -0.27483 | -0.3733654 | 2.57870351 | 1.67204812 | 0.900639045 | 4 | 2.1385957 | 5543.75 |
| p.Ser232Ter | 232 | -0.2099892 | 0.914008 | 0.86875547 | 0.23132555 | 0.451024963 | 4 | 0.2913532 | 6715.75 |
| p.Thr244Ter | 244 | -0.7214174 | 1.13289427 | 0.10128334 | 0.38384288 | 0.224150772 | 4 | 0.5868373 | 3009.5 |
| p.Lys245Ter | 245 | 0.03485876 | -0.157991 | 2.06889508 | 0.15200309 | 0.524441481 | 4 | 1.0764842 | 3761.75 |
| p.Thr246Ter | 246 | -0.6577678 | 0.04681054 | 2.49669305 | -0.4814992 | 0.35105915 | 4 | 2.1357324 | 12526.8 |
| p.Ser255Ter | 255 | -0.8606106 | 0.65014389 | 2.3616378 | 1.28324015 | 0.858602812 | 4 | 1.8128603 | 11144 |
| p.Arg273Ter | 273 | -0.1322526 | -0.4503775 | 1.04083372 | 1.17232858 | 0.407633048 | 4 | 0.6711203 | 8803.25 |
| p.Lys277Ter | 277 | -0.6567952 | -0.302425 | 1.59991284 | -0.0625845 | 0.144527019 | 4 | 1.0009755 | 2643.5 |
| p.Leu295Ter | 295 | -0.6927507 | 0.22758912 | 1.92913602 | 1.01591956 | 0.61997349 | 4 | 1.2492967 | 11863.8 |
| p.Thr303Ter | 303 | 0.69489956 | -0.56474 | 2.57968756 | 0.51733608 | 0.806795808 | 4 | 1.7068544 | 9129 |
| p.Arg343Ter | 343 | -0.2488271 | -1.221819 | 1.56408377 | -0.7538747 | -0.165109257 | 4 | 1.4867992 | 7965.75 |
| p.Ser344Ter | 344 | 0.76553475 | -0.0627589 | 2.53238581 | 1.48471103 | 1.179968173 | 4 | 1.2126758 | 8952.25 |
| p.Arg368Ter | 368 | -0.5305988 | 0.6382901 | 1.74393755 | -1.2319785 | 0.154912607 | 4 | 1.7173489 | 10408.3 |
| p.Asp370Ter | 370 | 0.78249973 | 0.12868202 | 3.38904229 | 0.19585226 | 1.124019075 | 4 | 2.366385 | 2375.75 |
| p.Arg373Ter | 373 | -0.5662882 | 0.19521041 | 2.0135976 | 0.27180462 | 0.478581104 | 4 | 1.190361 | 4859.75 |
| p.Gln397Ter | 397 | 0.48602065 | 0.09206055 | 3.56884284 | 0.26686119 | 1.103446306 | 4 | 2.7273901 | 1582.5 |
| p.Thr405Ter | 405 | 0.38724643 | -0.8204506 | 2.75002175 | 0.39854938 | 0.678841744 | 4 | 2.2337519 | 1426.75 |
| p.Ser411Ter | 411 | -0.5080664 | -0.2617944 | 2.04064198 | 1.55107213 | 0.70546332 | 4 | 1.6353328 | 7928.25 |
| p.Ser435Ter | 435 | -0.1383475 | -1.0795413 | 2.6811195 | 1.03046528 | 0.623424003 | 4 | 2.6267267 | 7081 |
| p.Tyr446Ter | 446 | -1.4412921 | -0.9268336 | 1 | -1.4888577 | -0.714245865 | 4 | 1.3708174 | 1514 |
| p.Ser455Ter | 455 | -0.534512 | 0.53568558 | 1.73758464 | -0.682443 | 0.264078806 | 4 | 1.2595468 | 15553.5 |
| p.Thr458Ter | 458 | -0.7114197 | -0.3778567 | 1.36678233 | 1.81149009 | 0.522249 | 4 | 1.5691693 | 7426.5 |
| p.Lys469Ter | 469 | 0.11028391 | -0.8625334 | 2.1229815 | 0.64711898 | 0.504462745 | 4 | 1.5546697 | 10088 |

| | | | | | | | | | |
|---|---|---|---|---|---|---|---|---|---|
| p.Thr499Ter | 499 | -0.5291522 | -0.0113407 | 1.12653241 | -0.8888244 | -0.075696238 | 4 | 0.7720984 | 12163.5 |
| p.Gly503Ter | 503 | 0.0426052 | -0.9123527 | 1.53713398 | 2.21386835 | 0.720313702 | 4 | 2.0075964 | 14776.5 |
| p.Pro504Ter | 504 | -0.1230485 | -0.0374747 | 3.34965258 | 1.57978226 | 1.192227897 | 4 | 2.6822667 | 6451 |
| p.Asn505Ter | 505 | -1.5389479 | 0.47613922 | 2.45204453 | 1.48026512 | 0.717375233 | 4 | 2.9134225 | 5736.75 |
| p.Gly506Ter | 506 | -1.4587724 | -0.2725259 | 1.35033181 | 0.87714325 | 0.124044201 | 4 | 1.5778386 | 2833.5 |
| p.Ser507Ter | 507 | -0.2746433 | -0.9318714 | 2.72246602 | 0.59335517 | 0.527326607 | 4 | 2.5318038 | 7767.25 |
| p.Lys509Ter | 509 | -0.2155298 | -0.2355946 | 1.48308289 | -0.3541513 | 0.169451798 | 4 | 0.7706868 | 10832.8 |
| p.Ser510Ter | 510 | -0.8344504 | 0.12619591 | 1.39005082 | -0.606356 | 0.018860098 | 4 | 1.0035729 | 13773.8 |
| p.Pro533Ter | 533 | -0.1934919 | -0.144468 | 2.1859834 | 1.20883517 | 0.764214664 | 4 | 1.3206733 | 21353 |
| p.Tyr537Ter | 537 | 0.36852698 | -0.5573749 | 1.30220986 | -0.3820027 | 0.182839809 | 4 | 0.7181446 | 12715 |
| p.Glu538Ter | 538 | 0.56478462 | 1.98142789 | 1.13750352 | 2.68072148 | 1.59110938 | 4 | 0.8662346 | 1111 |
| p.Ser557Ter | 557 | -0.6618053 | -1.1893206 | 1.60279579 | -1.0100677 | -0.314599448 | 4 | 1.6819232 | 5491.5 |
| p.Tyr566Ter | 566 | -1.0680949 | 0.60550608 | 2.24488706 | 0.37958866 | 0.540471731 | 4 | 1.840877 | 4272 |
| p.Tyr597Ter | 597 | 0.02004151 | -0.2064882 | 1.3811673 | 1.16938151 | 0.591025542 | 4 | 0.6402902 | 11303 |
| | | | | | Mean | 0.589036861 | | | |
| | | | | | St Dev | 0.498751696 | | | |

Table S5 (TAP2 ClinVar variants)

| VUS | functional score | benign | functional score | pathogenic | functional score | cancer assoc | functional score |
|---|---|---|---|---|---|---|---|
| G34R | 1.37955086 | R313H | 0.99285416 | R273* | 0.40763305 | R369Q | 1.20670955 |
| L111F | n/a | T665A | 1.13536733 | | | D370N | n/a |
| Y172F | 1.61920326 | | | | | D638A | n/a |
| S219F | 0.06740792 | | | | | | |
| R273Q | 0.9588095 | | | | | | |
| L295V | 1.3114607 | | | | | | |
| R343H | n/a | | | | | | |
| R368P | 1.30394422 | | | | | | |
| T499M | 1.33125539 | | | | | | |
| A500V | 1.72915145 | | | | | | |
| P533H | 1.7209542 | | | | | | |
| A618S | 1.82285138 | | | | | | |

Table S6 (DNA primers used)

| | |
|---|---|
| Barcode TOP strand: | TCGAGCTCAGACACTACTGAGGCGCGCCAGCCANNNNNNNNNNNNNNNNNNCTCATACAGGCGGTACGTCTGCTTTGGTCATGCA |
| Barcode BOTTOM strand: | TGACCAAAGCAGACGTACCGCCTGTATGAGNNNNNNNNNNNNNNNNNNTGGCTGGCGCGCCTCAGTAGTGTCTGAGC |
| Nest1-Fwd-N1: | ACACTCTTTCCCTACACGACGCTCTTCCGATCTNGCTCAGACACTACTGAGGCG |
| Nest1-Fwd-N2: | ACACTCTTTCCCTACACGACGCTCTTCCGATCTNNGCTCAGACACTACTGAGGCG |
| Nest1-Rev-N1: | GTGACTGGAGTTCAGACGTGTGCTCTTCCGATCTNCCAAAGCAGACGTACCGCCTG |
| Nest1-Rev-N2: | GTGACTGGAGTTCAGACGTGTGCTCTTCCGATCTNNCCAAAGCAGACGTACCGCCTG |
| Seq-i701 | CAAGCAGAAGACGGCATACGAGATCGAGTAATGTGACTGGAGTTCAGACG |
| Seq-i702 | CAAGCAGAAGACGGCATACGAGATTCTCCGGAGTGACTGGAGTTCAGACG |
| Seq-i703 | CAAGCAGAAGACGGCATACGAGATAATGAGCGGTGACTGGAGTTCAGACG |
| Seq-i704 | CAAGCAGAAGACGGCATACGAGATGGAATCTCGTGACTGGAGTTCAGACG |
| Seq-i705 | CAAGCAGAAGACGGCATACGAGATTTCTGAATGTGACTGGAGTTCAGACG |
| Seq-i501 | AATGATACGGCGACCACCGAGATCTACACTATAGCCTACACTCTTTCCCTACACG |
| Seq-i502 | AATGATACGGCGACCACCGAGATCTACACATAGAGGCACACTCTTTCCCTACACG |
| Seq-i503 | AATGATACGGCGACCACCGAGATCTACACCCTATCCTACACTCTTTCCCTACACG |

Primers for TAP2 sgRNA

| | |
|---|---|
| TAP2 sgRNA #11 Position 1 Fwd | GCTTTATATATCTTGTGGAAAGGACTGAGAGAAAGGAGGGTGAGTCGGTTTAAGAGCTAAGCTGGAAACAGCATAGCAAGTT |
| TAP2 sgRNA #12 Position 2 Fwd | ATATCCCTTGGAGAAAAGCCTTGTTGCAGGCTTCGCAAGAGCACATGTTTTAGAGCTAGAAATAGCAAGTT |
